# Beyond canonical: Hidden phosphoregulatory control in RAS signaling reveals context-dependent rewiring in Y1 tumor cells

**DOI:** 10.64898/2026.11.16.760319

**Authors:** Fabio Montoni, Toshiki Hanada, Doi Kentaro, Rosangela Aparecida Moreira Wailemann, Katia Andrea de Menezes Torres, Thompson Eusébio Pavan Torres, Jun Adachi, Marcelo da Silva Reis, Yayoi Natsume-Kitatani

## Abstract

Cell signaling pathways are not static. Regulatory mechanisms finely control the signal across complex networks to promote signal propagation. Among them, phosphorylation is one of the most important regulatory mechanisms, in which signal cascades propagate signals to the nucleus, enabling responses. Phosphorylation events can be regulated by guanine nucleotide exchange factors (GEFs) and GTPase-activating proteins (GAPs) to coordinate the activation state of small G-proteins. In the Y1 cell derived from the adrenal gland of a mouse, the tumor phenotype is driven by constitutive overexpression of non-mutated KRAS with high SOS (Son of Sevenless)-mediated KRAS-GTP levels, activating downstream ERK signaling. Our previous finding showed that SOS is not a sole GEF in KRAS activation, pointing that much of the GEF and GAP regulatory system remains unknown. This motivated the generation of a comprehensive time-resolved proteome (DIA) and TMT phosphoproteome (DDA) under FGF2 stimulation combined or not with the SOS inhibitor BI-3406 (SOSi) to investigate adaptive signaling dynamics. Our findings revealed ERK feedback relief through parallel pathway activation and differential phosphorylation dynamics consequent to SOS inhibition, revealing novel layers of phospho-signaling rewiring beyond canonical RAS activation.

## Introduction

Cellular dynamics are not just a set of isolated molecular effects, and cellular architecture is not static. Their dynamics are governed by regulatory mechanisms, which finely control the signal across the complex networks to promote signal propagation, response, and effect (**Kholodenko, 2006**).

Among the regulatory mechanisms, phosphorylation is one of the most relevant. The reversible addition of a phosphate group to serine, threonine, or tyrosine amino acids by protein kinases promotes conformational changes that alter the physicochemical properties of proteins (**Ardito et al., 2017**). This alteration enables phosphorylation cascades that propagate signals to the nucleus to promote multiple cellular responses. Phosphorylation events, in turn, are fine-tuned by several regulatory mechanisms. Among them, guanine nucleotide exchange factors (GEFs) and GTPase-activating proteins (GAPs) coordinate the activation state of small G-proteins (**Bos, Rehmann e Wittinghofer, 2007**), thereby regulating downstream phosphorylation cascades and playing important roles in cancer signaling (**Singh et al., 2017**), especially in tumors that have no mutations in their sequence.

That is the case of the Y1 adrenocortical tumor cell, a well-established model of RAS-dependent tumorigenesis (**Schimmer, 1979**). The tumor phenotype is driven by the combination of constitutive overexpression of non-mutated KRAS (**Schwab et al., 1983**), with high SOS-mediated KRAS-GTP levels, that activates downstream ERK signaling events (**Forti et al., 2002**). This cell line can be sensitized by Fibroblast Growth Factor 2 (FGF2) (**Costa et al., 2008; Dias et al., 2019**), which, combined with proteasome inhibitors, sustains hyperactivation of the ERK pathway, imposing an intolerable, death-leading stress signaling (**Dias et al., 2019**). This aligns with the growing premise that hyperactivation of cell signaling pathways surrounds tumor cells, due to their own lack of specific stress controls, compared to normal lineages (**Dias et al., 2024**). This shows that the Y1 cell line is still a relevant model to study carcinogenesis as a whole. Even though the importance and years of effort, much of the signaling of the Y1 cell is vastly unknown. A recent study from our research group revealed that SOS is not the sole GEF in KRAS activation, and RASGRP4 emerged as an indispensable factor for maintaining the Y1 cell tumor phenotype (**Montoni et al., 2026**).

This previously unknown factor, due to its conformational difference compared to SOS (**Ebinu et al., 1998**), suggests differential activation and response in this cell, showing that a largely unknown layer of consequent phosphorylation signaling is yet to be discovered. Even though targeting phosphoproteins can be an effective strategy in cancer therapy (**Radivojac et al., 2008; Singh et al., 2017)**, due to technical limitations and data-level issues, the majority of the phosphoproteome is still largely unknown, let alone their cell signaling effect (**Robichon, 2020; Houles, Yoon and Roux, 2024**). This motivated the generation of a comprehensive time-resolved proteome (DIA) and TMT phosphoproteome (DDA) across a 0–30-minute time course under FGF2 stimulation, a sensitizing factor in Y1 cells, combined or not with the SOS inhibitor BI-3406 (SOSi), to perturb one of the main KRAS activation axes and investigate adaptive signaling dynamics. Our findings revealed ERK feedback relief through parallel pathway activation, possibly enabled by differential phosphorylation site dynamics consequent to SOS inhibition. In addition, differential GEFs and GAPs, particularly from the RHO family, were reported, revealing novel layers of phospho-signaling rewiring beyond canonical Ras activation.

## Results

### Overview of differentially abundant proteins and phosphosites

Due to the long experimental design, especially for the phosphoproteome, precautions had to be taken to ensure the quality of the results. The pipeline: Log2 → keeping only <30% of missing rows → BPCA imputation → ComBat batch effect removal (for phosphoproteome) → zero mean-centering, ensured 8,786 proteins and 14,512 phosphosites with enough quality for downstream analysis. A full scheme of the experimental design can be observed in **Figure 1**.

**Figure 1:**
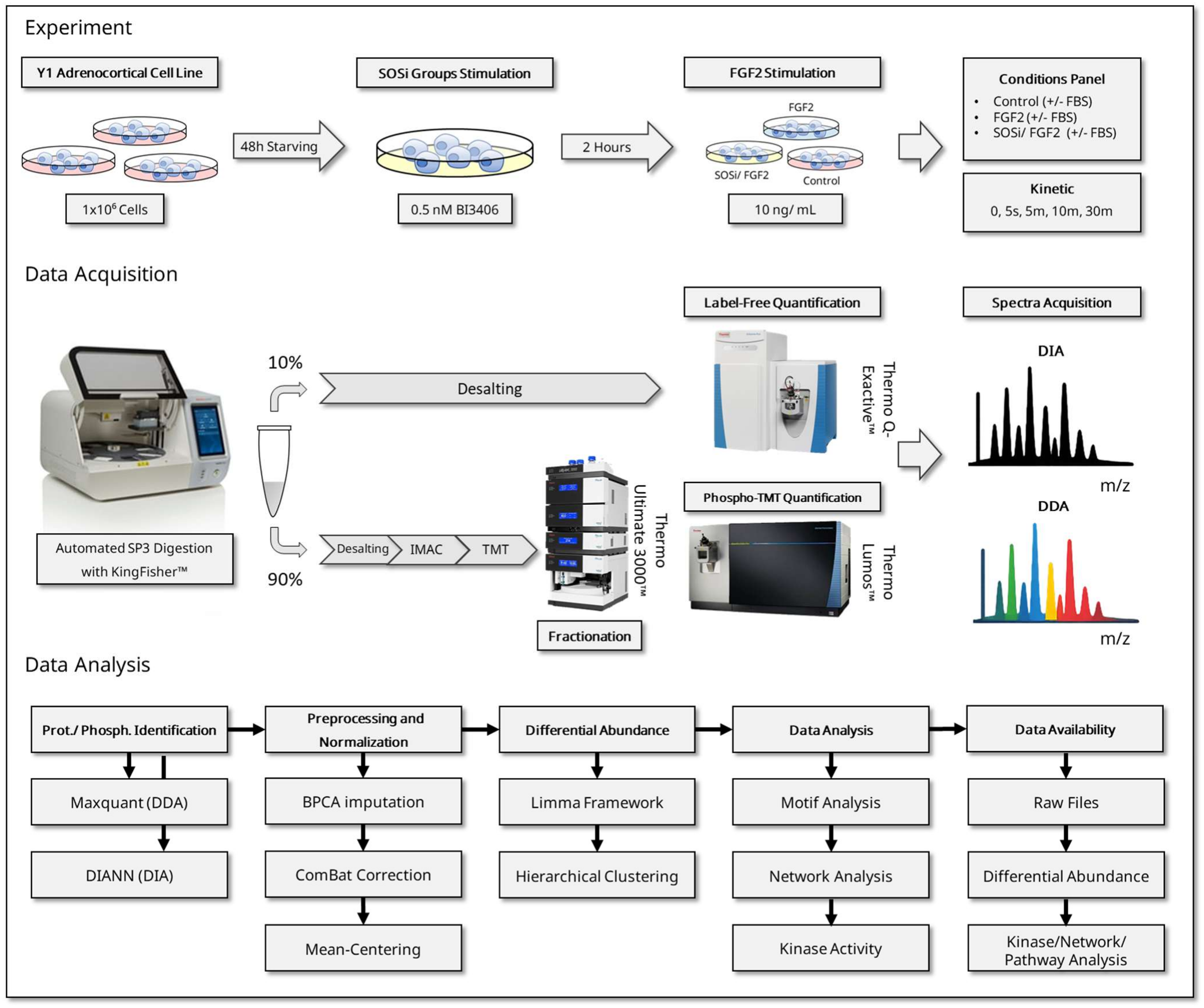
Experimental Design. Y1 mouse adrenocortical tumor cells were synchronized in G0/G1 by serum starvation and subjected to three experimental conditions: control, FGF2 stimulation (10 ng/mL), or SOS1 inhibition with BI-3406 (0.5 nM) in the presence or absence of serum. Samples were collected at 0 (control), 5 seconds, 5, 10, and 30 minutes in biological triplicates. Cell lysates were processed for proteomic and phosphoproteomic analyses, with the majority of the material allocated to phosphopeptide enrichment and the remainder to total proteome analysis. Phosphoproteome and proteome datasets were acquired using Orbitrap-based mass spectrometry platforms. Statistically significant changes were identified using linear models implemented in the limma framework.

A comprehensive proteome and phosphoproteome of Y1 adrenocortical tumor cells were investigated using high-resolution mass spectrometry. In total, 72 samples were analyzed across a 0–30 min time course under 10 ng/mL FGF2 (**Salotti et al., 2013**) alone or combined with the SOS inhibitor BI-3406 (0.5 nM), in the presence or absence of FBS stimulation (**Supplemental Figure S1**). DIA-NN analysis (precursor FDR 1.0%; protein group FDR 1.0%) identified 8,786 protein groups (**Supplemental Figure S2A**). MaxQuant phosphoproteome analysis (PIF ≥50%, FDR = 0.01, PTM localization probability >0.75) identified 14,512 phosphosites (**Supplemental Figure S2B**). A gene-centric analysis of both proteomes revealed that there are 3,377 genes in common on both proteomes, and 5,358 protein groups exclusively present in the proteome and 541 exclusive genes in the phosphoproteome (Figure 2A). The most frequently phosphorylated site in the phosphoproteome analysis is Serine (85.7%), followed by Tyrosine (13.3%) and Threonine (1.0%) (**Figure 2B**), and the motif identified for the significant phosphorylations was R-X-X-S-P (**Supplemental Figures S3 and S4**), which is highly correlated with MAPK (**Eblen, 2018**) and CDK activation (**Errico et al., 2010**). The significance analysis of the proteome using the Limma framework revealed 111 of the 8,786 protein groups identified (**Supplemental Figure S5**), corresponding to approximately 1.26% of the total identifications (**Figure 2C**). This demonstrates that the majority of the proteome remained stable throughout the time series established in this experiment. Additionally, it can be observed that the times where the largest changes occur among the 111 proteins are at the times with the SOS1 inhibitor, in which the inhibitory stimulus preceded the 30-minute FGF2 stimulation by 2 hours, such as JUNB (UNIPROT ID: P17275), SERPINE1 (UNIPROT ID: P05121) and CCN1 (UNIPROT ID: O00622), which remained increased at the times SOSi/FGF2–FBS 5 min (2) and SOSi/FGF2 + FBS. For the phosphoproteome, the Limma significance method revealed 10,416 of 14,512 phosphosites significant in at least one condition (**Supplemental Figure S6**), corresponding to approximately 72,2% of the total identifications (**Figure 2D**), showing higher complexity when compared to the proteome.

**Figure 2:**
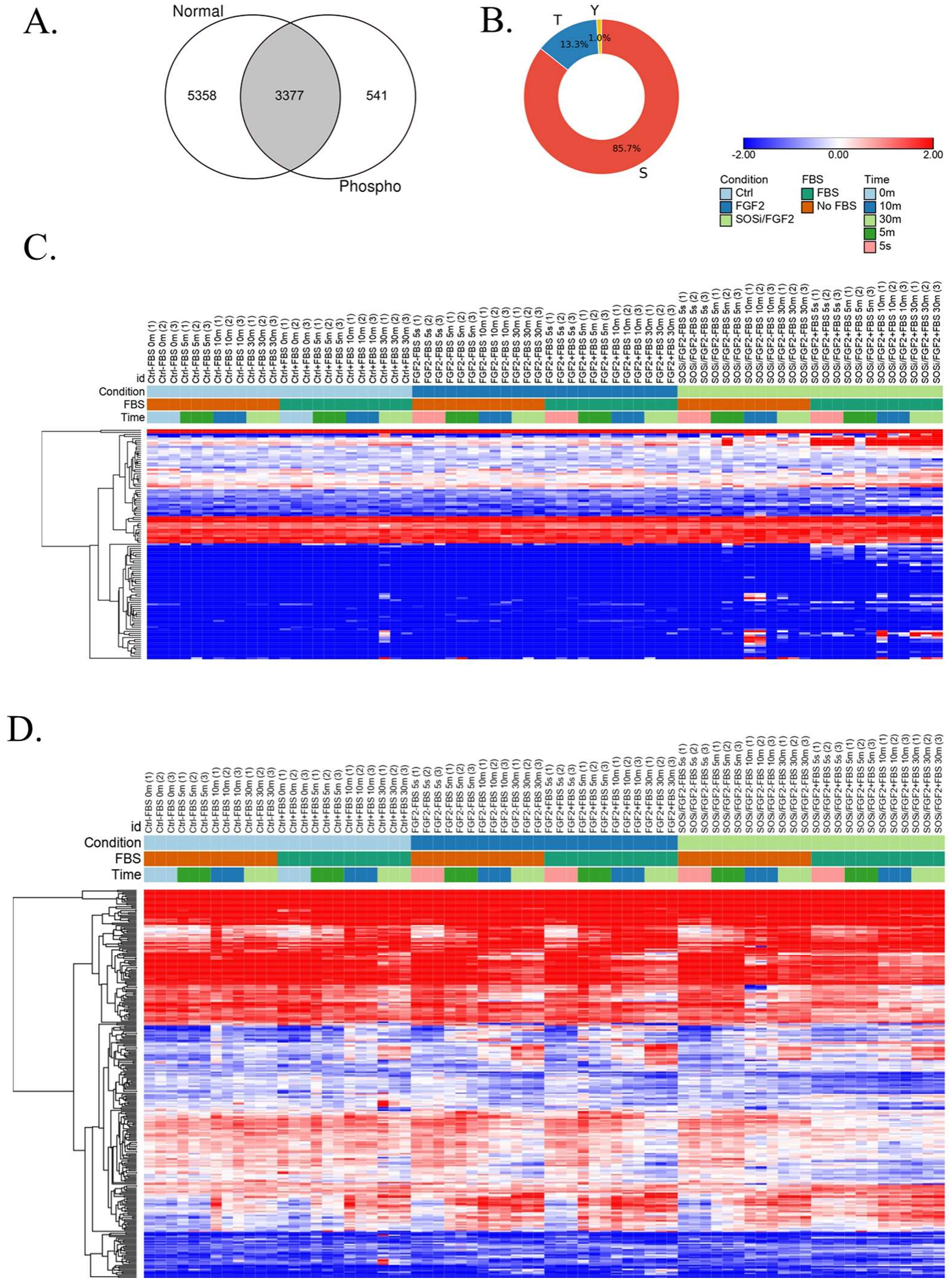
Overview of differentially abundant proteins and phosphosites. Overview of differentially abundant proteins and phosphosites. **A**. Venn diagram of Proteome and Phosphoproteome common significant hits in a gene-centric comparison. **B**. Distribution of phosphorylated amino acid residues in the phosphoproteome dataset. **Legend**: **S**: Serine, **T**: Threonine, and **Y**: Tyrosine. **C**. Limma-significant proteome hierarchical clustering using Euclidean distance (N=111). **D**. Top 300 (lowest Adj. p-value criteria) Limma-significant phosphosites hierarchical clustering using Euclidean distance (N=300). The table containing all the limma results for the phosphoproteome can be accessed in the Companion Paper Montoni, F. (2026c).

### Differential phosphorylation abundance suggests GEFs and GAPs rewiring under FGF2 and SOS inhibition + FGF2 perturbations in Y1 cell

As previously mentioned, the Y1 cell tumor phenotype is dependent on the combination of the KRAS gene amplification and GEF-mediated activation. Over the years, it was believed that the activation role was restricted to the GEF SOS. However, our research group recently showed that SOS is not the only GEF needed to sustain the high basal levels of KRAS in Y1 adrenocortical cells. The study revealed RASGRP4 as indispensable for this phenotype (**Montoni et al., 2026**). In addition, this study also shows that the interaction of GEFs and GAPs is more complex than actually perceived, and the results warrant deeper studies. Taking their importance into account, a survey of general GEFs and GAPs was performed to select the most relevant GEFs and GAPs found in the Y1 cell normal and phosphoproteome. In total, 28 GEFs and GAPs were identified in the proteome dataset (**Figure 3A**). These proteins were classified into their small GTPase families based on their annotated functions. Among GEFs, the RasGEF SOS1 was detected (UniProt ID: Q62245), two RalGEFs, including RGL2 (UniProt ID: Q61193), ten RhoGEFs, including TIAM1 (UniProt ID: Q60610), and one ArfGEF, IQEC1 (UniProt ID: Q8R0S2). Among GAPs, three RalGAPs were identified, including SIPA1 (UniProt ID: P46062), six RhoGAPs, including RHG05 (UniProt ID: P97393), and one ArfGAP, ASAP1 (UniProt ID: Q9QWY8).

**Figure 3:**
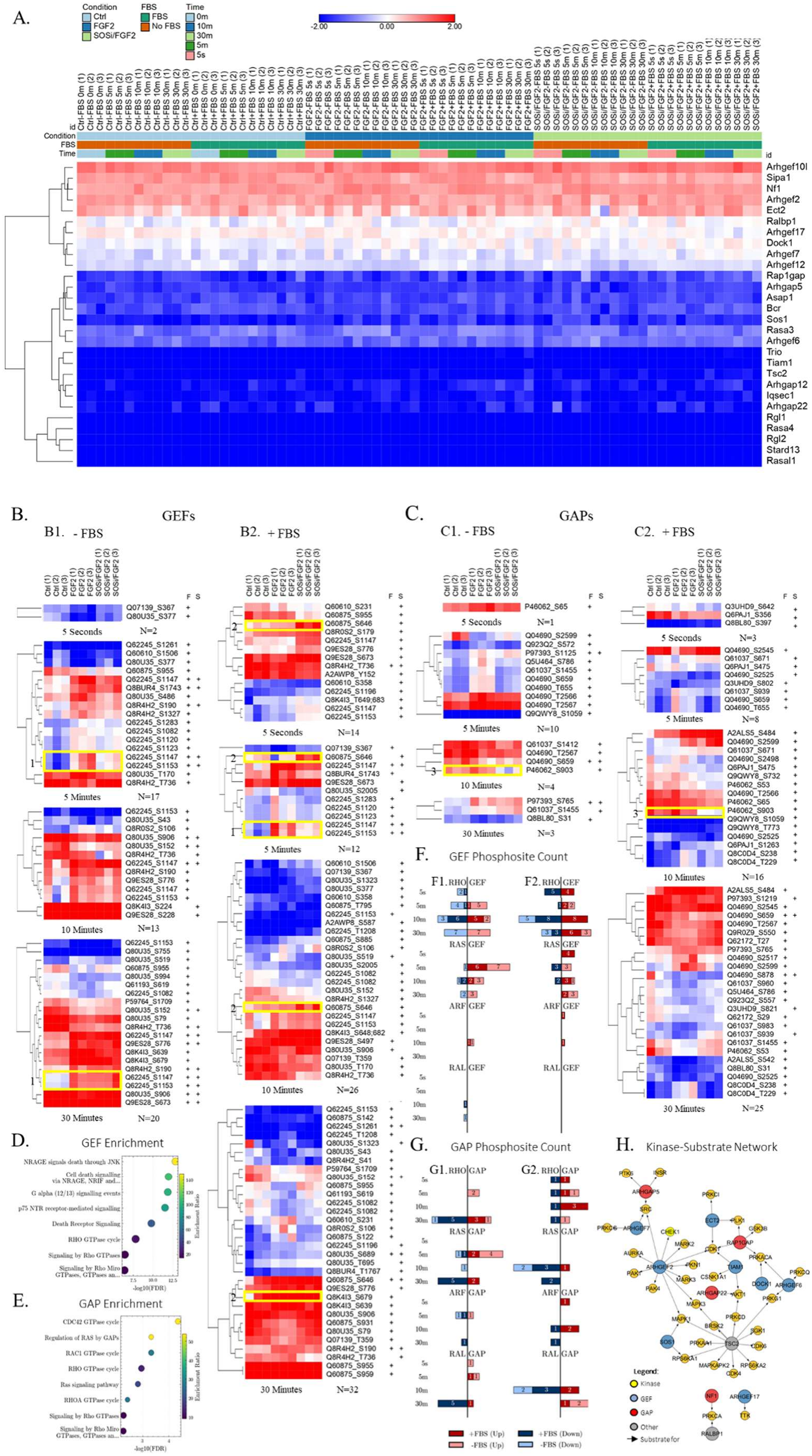
Differential phosphorylation abundance suggests GEFs and GAPs rewiring under FGF2 and SOS inhibition + FGF2 perturbations in Y1 cells. **A**. Hierarchical clustering of all GEFs and GAPs modules identified in the proteome analysis. **B**. Hierarchical clustering of all GEF modules identified in the phosphoproteome analysis. For each treatment and time point, only significant sites in at least one of the conditions (marked with “+”) were kept, following the p-value < 0.05 criterion from the Limma framework analysis. **B1:** GEF modules minus FBS from 5 seconds to 30 minutes. **B2:** GEF modules plus FBS from 5 seconds to 30 minutes. **C**. Hierarchical clustering of all GAP modules identified in the phosphoproteome analysis. For each treatment and time point, only significant sites in at least one of the conditions (marked with “+”) were kept, following the p-value < 0.05 criterion from the Limma framework analysis. **C1:** GAP modules minus FBS from 5 seconds to 30 minutes. **C2:** GAP modules plus FBS from 5 seconds to 30 minutes. **Legend**: **F:** Treated with FGF2. **S:** Treated with SOS inhibitor + FGF2. +: Limma-significant condition when compared to the control. **D**. Pathway enrichment of the GEFs identified using WebGestalt tools. Gene-centric Reactome pathway analysis was performed for all the GEFs identified in both proteome and phosphoproteome analyses. **E**. Pathway enrichment of the GAPs identified using WebGestalt tools. Gene-centric Reactome pathway analysis was performed for all the GAPs identified in both proteome and phosphoproteome analyses. **F**. GEFs phosphosite count. The GEFs module was subdivided into GEF sub-categories (RHO, RAS, ARF, and RAL). For each time point, the phosphosites were classified as “up” regulated and “down” regulated based on Limma results, counted and separated by time point. **F1**. GEF phosphosite count for FGF2 treatment. **F2**. GEF phosphosite count for SOS inhibitor + FGF2 treatment. **G**. GAPs phosphosite count. The GAPs module was subdivided into GAP sub-categories (RHO, RAS, ARF, and RAL). For each time point, the phosphosites were classified as “up” regulated and “down” regulated based on Limma results, counted and separated by time point. **G1**. GAP phosphosite count for FGF2 treatment. **G2**. GEF phosphosite count for SOS inhibitor + FGF2 treatment. **H**. Kinase-substrate network of the GEFs and GAPs identified using KiNet. This network reveals several kinases that the GEFs and GAPs identified in this dataset are substrates for, if there is at least one documented phosphosite-kinase interaction. The interaction graph list and the phosphosite identification intersection between the literature and this dataset can be seen in **Supplemental Table 1. Legend: Yellow Protein Node:** Kinase, **Blue Protein Node:** GEF, **Red Protein Node:** GAP, **Grey Protein Node: Other. Edge:** Indicates an interaction from a substrate to a specific kinase.

None of the proteome GEFs or GAPs were significantly regulated (log2(1.5) and an adjusted p-value below 0.05), showing stable abundance across all the conditions and time points. Differently, in the phosphoproteome data several sites were significantly regulated across the conditions (**Figure 3B and 3C**). In total, 206 sites (redundant) were modulated across the conditions, with most of the genes included in the proteome set, except the genes Asap3 (UniProt ID: Q5U464), Dlc1 (UniProt ID: Q9R0Z9), Agap2 (UniProt ID: Q3UHD9), Dock4 (UniProt ID: P59764) and Psd (UniProt ID: Q5DTT2).

**Table 1.**
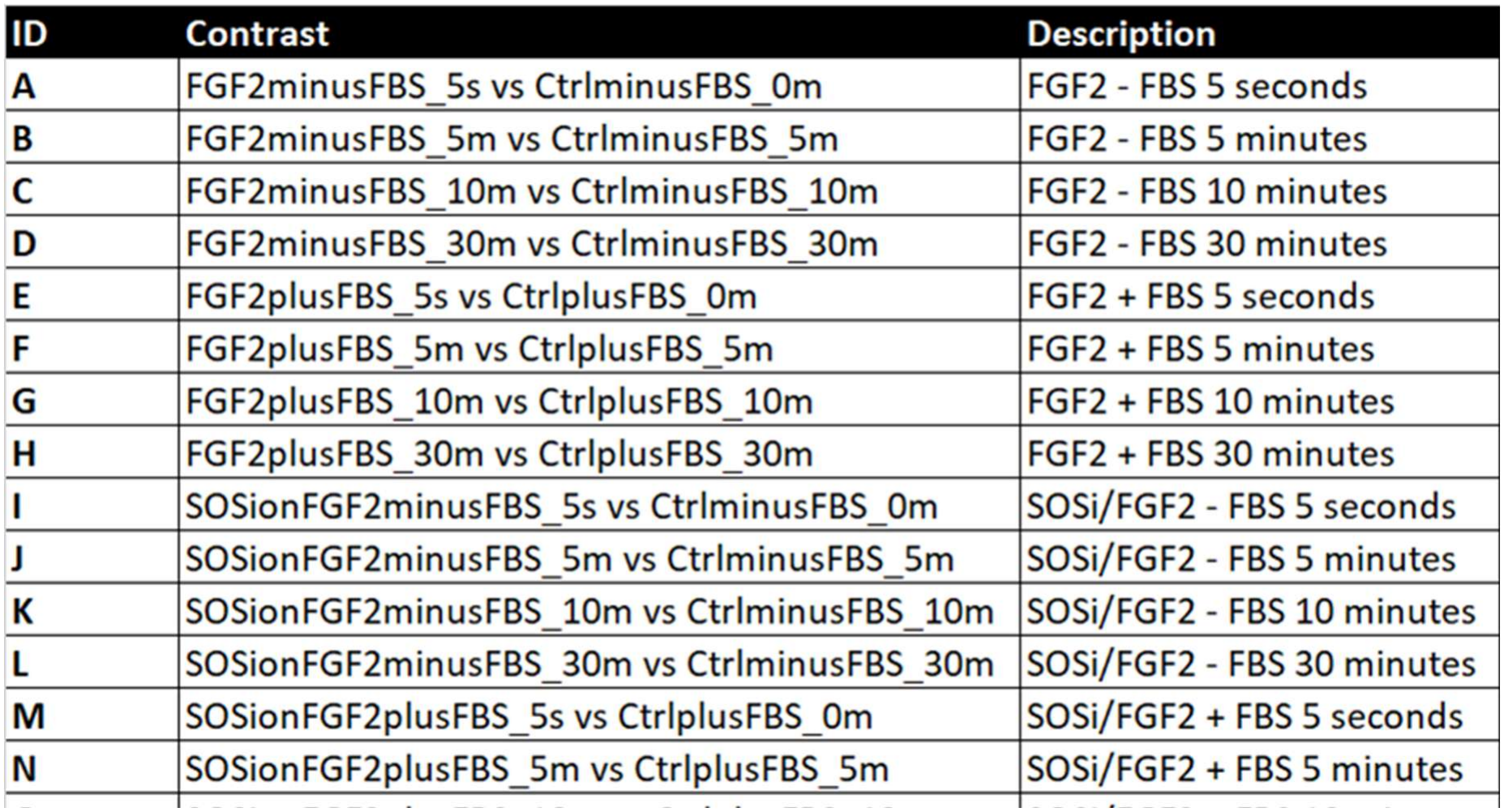
Limma contrasts assignment, for both proteome and phosphoproteome.

Due to the complex behavior of phosphosites throughout the time series and treatments, the sites were divided into two major modules, the GEF modules (**Figure 3B**) and the GAP modules (**Figure 3C**), which were further separated into the submodules B1 and B2 (**Figure 3B1 and 3B2**) for the GEFs and submodules 3C1 and 3C2 (**Figure 3C1 and 3C2**) for the GAPs. The first submodule corresponds to GEFs in the absence of fetal bovine serum (FBS), from 5 seconds to 30 minutes (**Figure 3B1**), containing 49 phosphosites. The second submodule corresponds to GEFs in the presence of fetal bovine serum (FBS), from 5 seconds to 30 minutes (**Figure 3B2**), containing 84 phosphosites. The third submodule corresponds to GAPs in the absence of fetal bovine serum (FBS), from 5 seconds to 30 minutes (**Figure 3C1**), containing 18 phosphosites. Lastly, the fourth submodule corresponds to GAPs in the absence of fetal bovine serum (FBS), from 5 seconds to 30 minutes (Figure 3C2), containing 52 phosphosites.

It is also interesting to note that the submodules treated without fetal bovine serum, for both GEFs and GAPs, showed a greater number of significant sites for FGF2 treatment (46/49 significant sites on submodule 3B1 (**Figure 3B1**) and 14/18 significant sites on submodule 3C1 (**Figure 3C1**). In contrast, treatment with fetal bovine serum showed a greater number of significant sites for the SOS inhibitor plus FGF2 condition, for both GEFs and GAPs sub-modules (42/52 significant sites on submodule B2 (**Figure 3B2**), from 5 seconds to 10 minutes, and 22/27 significant sites on submodule C2 (**Figure 3C2**), from 5 seconds to 10 minutes). The exception to this pattern occurs in both submodules B2 (**Figure 3B2**) and C2 (**Figure 3C2**) at the 30-minute time point, where FGF2 at 30 minutes presents more significant sites than the SOS inhibitor treatments (26/32 and 22/25, respectively). This result may suggest that the addition of fetal bovine serum, which contains several factors besides FGF2, in treatments with SOS1 inhibitor allows for greater signaling dynamics and a more exacer-bated response to inhibition. This may facilitate feedback effects and parallel pathway responses that compensate for the deleterious effect of the inhibitor, and this dynamic appears to occur up to 10 minutes of FGF2 stimulation.

Even though the complexity, the pattern of some sites are worth highlighting. For instance, some of the SOS1 (Uni-Prot ID: Q62245) phosphosites present a differential abundance when comparing both FGF2 and SOSi + FGF2 treatments (**Figure 3B, highlighted in yellow #1**). For both GEF submodules B1 (**Figure 3B1**) and B2 (**Figure 3B2**), the sites S1147 and S1153 are downregulated in the control conditions, upregulated upon FGF2 stimulation (B1: S1147: log_2_FC ≈ 1.93; S1153: log_2_FC ≈ 2.15 and B2: S1147: log_2_FC ≈1,92; S1153: log_2_FC ≈ 2,11), and notably attenuated in the SOS inhibition (B1: S1147: log_2_FC ≈ 1.13; S1153: log_2_FC ≈ 1.37 and B2: S1147: log_2_FC ≈ 0,99; S1153: log_2_FC ≈ 1,17) in the 5 minutes treatments, suggesting modulation by the inhibitor in these specific sites. However, this behavior seems not to be sustained at later time points, and both FGF2 and SOSi + FGF2 treatments display similar behavior, leading to the hypothesis that the attenuation of these sites has been compensated over the time course. For instance, the sites at the time point 30 minutes in the submodule B1 (**Figure 3B1**) for FGF2 (B1: S1147: log_2_FC ≈ 1,06; S1153: log_2_FC ≈ 1,47) are more similar to the SOSi + FGF2 treatment (B1: S1147: log_2_FC ≈ 1,28; S1153: log_2_FC ≈ 1,58). These sites have been noted on the PhosphoSitePlus platform, where Q62245_S1147 (**PhosphoSitePlus, 2026c**) and Q62245_S1153 (**PhosphoSitePlus, 2026c**) have been found to be upregulated in 3T3-L1 differentiated adipocytes after insulin stimulation, with further AKT-mTORC2 activity (**Humphrey et al., 2013**). Another example is (**Figure 3B, highlighted in yellow #2**), where the phosphosite of the Rho GEF Arhgef2 (UniProt ID: Q60875) Q60875_S646, which consistently upregulated in the SOSi + FBS conditions from 5 seconds to 30 minutes (e.g. B2: S646: log_2_FC ≈1,53 in the time-point of 5 seconds) in the submodule B2 (**Figure 3B2**). This site is well characterized in the PhosphositePlus database, where several publications has mentioned a number of direct and indirect regulators (**PhosphoSitePlus, 2026a**), including murine KRAS and PAK4 has been reported as indirect regulators for this specific site (**Gnad et al., 2013**). When it comes to GAPs (**Figure 3B, highlighted in yellow #3**), the SIPA1 (Uniprot ID: P46062) phosphosites have been found downregulated in SOS inhibitor + FGF2 10 minutes treatments without FBS in the submodule C1 (C1: S903: log_2_FC ≈-1.16) (**Figure 3C1**), and in SOS inhibitor + FGF2 10 minutes treatments with FBS in the module C2 (C2: S903: log_2_FC ≈-1,57) (**Figure 3C2**). In the PhosphoSitePlus database (**PhosphoSitePlus, 2026b**), this site is also mentioned as part of the AKT-mTORC2 activity, even though its major mechanism is still not fully disclosed (**Humphrey et al., 2013**).

In addition, the GEFs and GAPs have been enriched in Reactome pathways. Consistent with what was observed previously, the GEFs (**Figure 3D**) revealed a strong overrepresentation of signaling pathways related to cell death and Rho GTPase regulation. Among the most significantly enriched pathways were NRAGE signals death through JNK and cell death signaling via NRAGE, NRIF, and NADE, both displaying high enrichment ratios (>100) and extremely low FDR values (<10^−12^), supported by the consistent presence of eight genes across these categories (Arhgef10l, Ect2, Trio, Arhgef2, Sos1, Arhgef17, Arhgef6, Arhgef7). In parallel, pathways associated with G protein signaling, particularly G alpha (12/13) signaling events, and p75 NTR receptor-mediated signaling were also significantly enriched. Notably, Rho GTPase-related pathways, including RHO GTPase cycle, Signaling by Rho GTPases, and specific modules such as RHOA and RAC1 GTPase cycles, were consistently enriched, with multiple GEFs (e.g., Arhgef family members, Ect2, Trio, Dock1) contributing to this signal.

Pathway enrichment analysis of GAP-associated (**Figure 3E**) genes revealed a strong overrepresentation of signaling pathways related to Rho GTPase regulation. The most significantly enriched pathways included the CDC42 GTPase cycle and RAC1 GTPase cycle, both displaying high enrichment ratios (>25) and supported by multiple GAPs such as Arhgap5, Ralbp1, Bcr, Arhgap22, and Stard13. Consistently, broader pathways such as RHO GTPase cycle and Signaling by Rho GTPases were also enriched, reinforcing the central role of Rho GTPase regulatory mechanisms within this gene set. In addition, pathways related to RAS regulation were identified, including Regulation of RAS by GAPs and the Ras signaling pathway, driven by canonical RAS GAPs such as Nf1, Rasa3, Rasa4, and Rasal1.

Differently of what is stated in the literature for the Y1 cell, our data demonstrate that other GEFs besides the SOS RASGEFs and the recently validated essential RASGRP4, other categories of GEFs and also GAPs act dynamically under the various experimental conditions of this study. The next result (**Figure 3F**) summarizes the number of GEFs and GAPs that significantly changed across the treatments. Notably, RHO GEFs are the ones that suffer the most disturbances throughout the treatments, having 20 phosphosites upregulated, versus 24 phosphosites downregulated across the time points for the FGF2 treatment (**Figure 3F1**), followed by 25 upregulated and 25 downregulated for the SOS inhibitor + FGF2 treatment across the time points (**Figure 3F2**). Following, the RAS GEFs showed 21 phosphosites upregulated and six downregulated in the FGF2 treatment across the time points (**Figure 3F1**). In sequence, for the SOS inhibitor + FGF2 treatment, 16 sites were upregulated, and five were downregulated (**Figure 3F2**). Also, the presence of other GEF classes can be observed in both treatments (**Figure 3F1 and 3F2**). This result shows that both GEFs and GAPs dynamically change their networks, especially RHO GEFs, with the SOS inhibitor enhancing this effect.

In the case of GAPs, the RHO family remains practically absent, with only two phosphosites upregulated in 5 minutes, and only in 30 minutes does it show six downregulated sites and four upregulated sites in the FGF2 treatment (**Figure 3G1**). However, in the SOS inhibitor + FGF2 treatment, RHO GAPs are consistently present in all treatments, with a total of four downregulated sites versus seven upregulated sites (**Figure 3G2**). For the RAS GAPs, after five minutes of treatment, eight sites were upregulated and eight downregulated in the FGF2 treatment without fetal bovine serum (**Figure 3G1**), whereas after 10 minutes, seven sites appeared downregulated and only one upregulated in the SOS inhibitor + FGF2 treatment (Figure 3G2). ARF GAPs also show a presence across the treatments, with three sites downregulated and 1 upregulated in the FGF2 treatment (**Figure 3G1**). For the SOS inhibitor + FGF2 treatments, two downregulated and three upregulated phosphosites can be observed (**Figure 3G2**). Interestingly, RAL GAPs showed important changes across the treatments. Four sites were upregulated, while five sites were downregulated in the FGF2 treatments (**Figure 3G1**). In the SOS inhibitor + FGF2 treatments, after 10 minutes, five sites were upregulated, while five sites were downregulated. This scenario shows that the GAPs might also play an important role in sustaining the Y1 cell dynamics; also, the changes observed in other GAP classes show that the crosstalk of other pathways may be fundamental for maintaining the ERK levels in this cell line.

Finally, the Kinase-Substrate relationship reveals a complex interaction between GEFs, GAPs, and kinases (**Figure 3H)**. The graph network demonstrates that several GEFs and GAPs found in this study are substrates for various kinases. The members of the graph kinases and their classification can be observed in **Table 2**. This result reinforces the ability of this cell to alter its dynamics in response to disturbances and *stimuli* to compensate for the effects, such as ARHGEF2, which was already pointed out as a KRAS effector in pancreatic cancer, and its presence difficults the success of the ERK blockage (**Kent, Sandi and Rottapel, 2019**). Another hypothesis that can be raised is that a blockage of an important pathway can also be compensated by another activation axis, such as Tiam1, which is a GEF that can activate ERK/STAT3 (**Song et al., 2024**). One limitation of this analysis is that KiNet converts the gene and only shows sites confirmed for Homo sapiens. The overlap level of all sites found in the KiNet database and our study is approximately 7% (Gene_site) (**Supplemental Table 1**); however, the functional overlap may be even higher, since phosphosites are often conserved among mammals, but not necessarily phosphorylated at those sites (**Freschi, Osseni e Landry, 2014**).

**Table 2.** Classification of kinases and non-kinase proteins identified in the kinase–substrate network, grouped by kinase family. Corresponding gene symbols are provided for each protein.

| **Class** | **Corresponding Gene Symbols** |
|---|---|
| AGC | AKT1, PKN1, PRKACA, PRKCA, PRKCD, PRKCG, PRKCI, PRKCQ, PRKG1, RPS6KA1, RPS6KA2, SGK1 |
| CAMK | BRSK2, CHEK1, MAPKAPK2, MARK2, MARK3, PRKAA1 |
| CK1 | CSNK1A1 |
| CMGC | CDK1, CDK4, CDK6, GSK3B, MAPK1, MAPK3 |
| STE | PAK1, PAK4 |
| TK | INSR, PTK6, SRC |
| Other | AURKA, PLK1, TTK |
| Non-kinase | ARHGAP22, ARHGAP5, ARHGEF17, ARHGEF2, ARHGEF6, ARHGEF7, DOCK1, ECT2, NF1, RALBP1, RAP1GAP, SOS1, TIAM1, TSC2 |

Taken together, these results even though does not unveil the mechanistic path taken to respond to the multiple perturbations, it clearly suggests that the Y1 cells need to rewire its behavior through GEF and GAP. In the next result, a comprehensive kinase activity analysis was performed to visualize the effect of the changes observed in the phospho-proteome dynamics as a whole.

### SOS inhibition affects multiple sites, and pathway rewiring occurs through differential phosphorylation-mediated pathway propagation

All cellular types possess their own phosphorylation pattern, which governs which signaling pathways will be active, maintaining cell homeostasis. When this pattern is broken, (e.g. stimulus such as FGF2, fetal bovine serum, or through a disturbance such as an SOS inhibitor), the cell needs to adapt to the conditions so that the response can rebalance the original constitutive levels, keeping the cell homeostasis. In this section, we demonstrate how the Y1 cell differentially adapts to the *stimuli* and how this signal is propagated mechanistically, taking the MAPK pathway as the basis.

Given its central role in MAPK signaling and the fact that the inhibition occurs at the SOS1 level, the investigation was initiated by examining its dynamics. The statistically relevant sites were mapped onto a radial diagram (**Figure 4A**), which revealed condition- and time-specific patterns. The broader spectrum of phosphosite regulation was obtained with the FGF2 treatments, where the sites S1283, S1120, S1123, and S1261 were detected exclusively, independent of serum availability. The site S1082 was detected in all conditions except SOSi −FBS, whereas T1208 was restricted to serum-containing conditions. Interestingly, S1196 was exclusively detected under SOSi +FBS conditions, suggesting that SOS1 inhibition may induce distinct context-dependent phosphorylation patterns within the SOS1 regulatory region. Additionally, the sites S1147 and S1153 were kept in all conditions and almost all time points, even in the SOSi-FBS conditions, where the SOS1 phosphosites were mostly inhibited.

**Figure 4:**
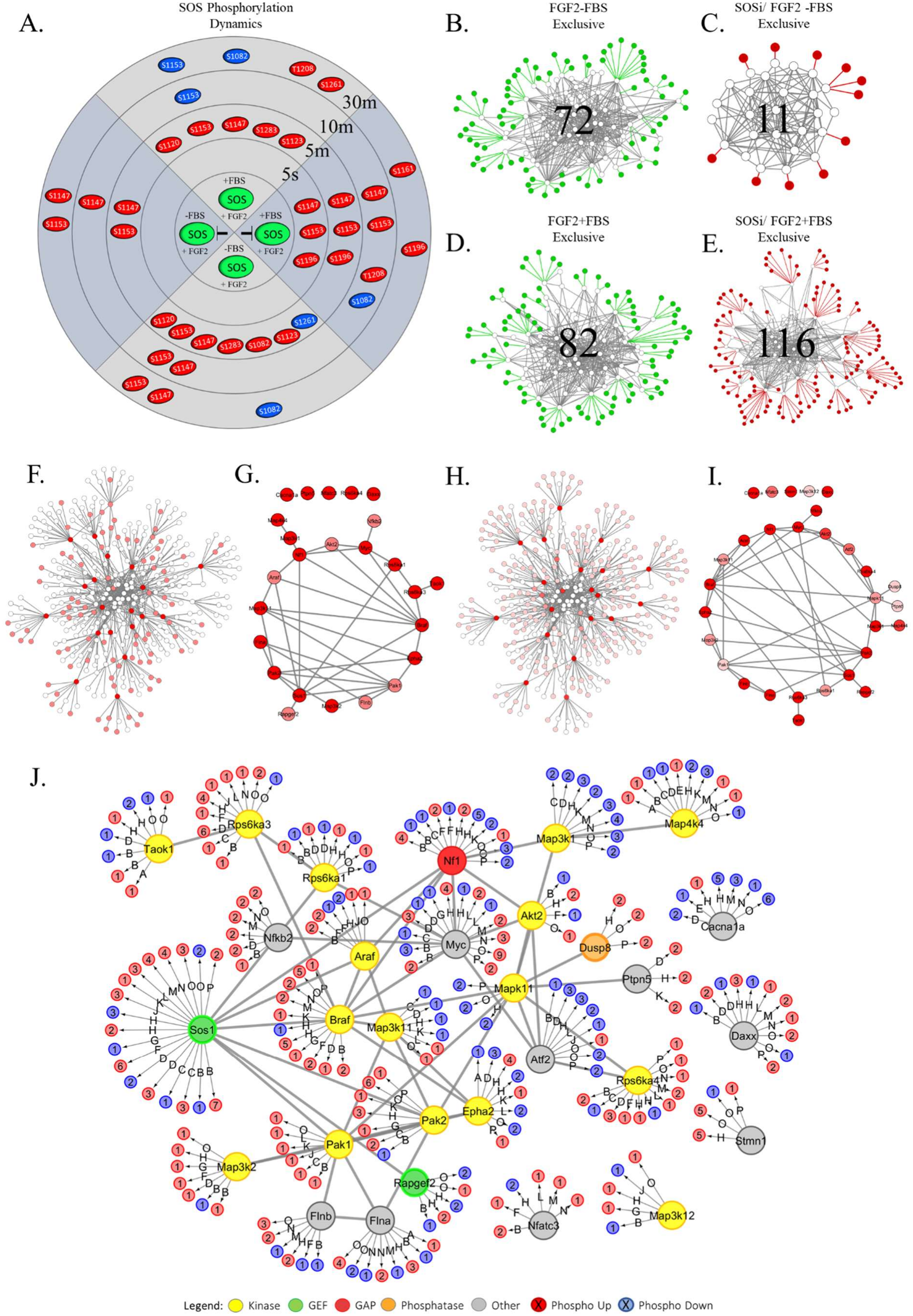
SOS inhibition affects multiple sites, and pathway rewiring occurs through differential phosphorylation-mediated pathway propagation. **A:** Radial graph of the propagation of SOS1 protein sites across treatments and time series. **Legend: Red Node:** Site upregulation. **Blue Node**: Site downregulation. **B:** Network of significant sites observed exclusively under the FGF2-FBS condition, compared to the significant sites under the SOSi/FGF2-FBS condition. **C:** Network of significant sites observed exclusively under the SOSi/FGF2-FBS condition, compared to the significant sites under the FGF2-FBS condition. **D:** Network of significant sites observed exclusively under the FGF2+FBS condition, compared to the significant sites under the SOSi/FGF2+FBS condition. **E:** Network of significant sites observed exclusively under the SOSi+FBS condition, compared to the significant sites under the FGF2+FBS condition. **F:** Sites with the most variations identified using the DyNet tool, which ranks sites that vary most between FGF2-FBS and SOSi/FGF2-FBS conditions. **G:** Network of top ranking proteins with the most variable phosphosite interactions between the FGF2-FBS and SOSi/FGF2-FBS treatments. **H:** Sites with the most variations identified using the DyNet tool, which ranks sites that vary most between FGF2+FBS and SOSi/FGF2+FBS conditions. **I:** Network of top ranking proteins with the most variable phosphosite interactions between the FGF2+FBS and SOSi/FGF2+FBS treatments. **J:** A protein interaction network was generated from the relevant phosphosites identified by LIMMA analysis and subsequently uploaded to the ProteomicsDB PTM Navigator. Phosphosites enriched for the MAPK pathway (mmu) were selected (pValue = 0.05 <) to reconstruct the interaction network using a combination of STRING and Cytoscape tools. **Legend**: **Yellow Protein Node:** Kinase, **Orange Protein Node:** Phosphatase. **Green Protein Node:** GEF, Red Protein Node: GAP, **Grey Protein Node:** Other Proteins. **Red node:** Upregulated phosphosite. **Blue node:** Downregulated phosphosite. **Edges:** Treatment where the phosphosite was found up/downregulated; **Contrast A:** FGF2 - FBS 5 seconds; **Contrast B:** FGF2 - FBS 5 minutes; **Contrast C:** FGF2 - FBS 10 minutes; **Contrast D:** FGF2 - FBS 30 minutes; **Contrast E:** FGF2 + FBS 5 seconds; **Contrast F:** FGF2 + FBS 5 minutes; **Contrast G:** FGF2 + FBS 10 minutes; **Contrast H:** FGF2 + FBS 30 minutes; **Contrast I:** SOSi/FGF2 - FBS 5 seconds; **Contrast J:** SOSi/FGF2 - FBS 5 minutes; **Contrast K:** SOSi/FGF2 - FBS 10 minutes; **Contrast L:** SOSi/FGF2 - FBS 30 minutes; **Contrast M:** SOSi/FGF2 + FBS 5 seconds; **Contrast N:** SOSi/FGF2 + FBS 5 minutes; **Contrast O:** SOSi/FGF2 + FBS 10 minutes; **Contrast P:** SOSi/FGF2 + FBS 30 minutes.

All these sites are located in the proline-rich region of the SOS1 protein (a.a. 1141–1300) (**McDonald et al., 2013**), or in its proximity, such as the site S1082. This region is an intrinsically disordered region (IDR) (**Tateno et al., 2024**), known for being a hub for GRB2 binding (**Terasawa et al., 1994**), ERK feedback effects, and pathway rewiring (**Kamioka et al., 2010**).

These results suggest that SOS inhibition resulted in inhibition of multiple phosphorylation sites, and the extracellular context (e.g., FGF2, fetal bovine serum) allows SOS1 to adapt to the inhibition. This idea is congruent with other author’s work, such as the work of Kamioka et al. 2010, which shows that the role of multiple SOS1 phosphorylations is more complex than initially perceived, and more precise consequences still need further investigation (**Kamioka et al., 2010**).

Regarding the whole phosphorylation profile, in total, 466 phosphosites from 65 proteins from the MAPK pathway were differentially abundant (KEGG mmu04010, enriched in PTM Navigator) (**Supplemental Figure S7**). Due to the large dataset size, the analysis was restricted to the top 30 proteins and their significant phosphorylations. In addition to differential abundance, exclusively abundant sites in each of the conditions were also observed. For the minus FBS conditions, 72 sites were exclusive to the FGF2-FBS treatment (**Figure 4B**, full-length figure on **Supplemental Figure S8**), while 11 sites were exclusive to the SOSi/FGF2-FBS treatment (**Figure 4C**, full-length figure on **Supplemental Figure S9**). As for the plus FBS conditions, 82 sites were exclusive to the FGF2+FBS treatment (**Figure 4D**, full-length figure on **Supplemental Figure S10**), while 116 sites were exclusive to the SOSi/FGF2+FBS treatment (**Figure 4E**, full-length figure on **Supplemental Figure S11**). These results demonstrate that cells differentially rewire signaling networks through condition-specific mechanisms, whereas the presence of fetal bovine serum broadens the spectrum of adaptive responses relative to treatments performed in its absence. Additionally, distinct proteins exhibited the greatest changes in phosphorylation-site profiles across conditions. DyNet analysis highlighted the nodes with the greatest variation between the FGF2-FBS versus SOSi/FGF2-FBS (**Figure 4F**, full-length figure on **Supplemental Figure S12**), with 25 proteins with significant variation levels among the treatments(raw DN-Score > 0.5) (**Figure 4G**), whereas FGF2+FBS versus SOSi/FGF2+FBS comparisons (**Figure 4H**, full-length figure on **Supplemental Figure S13**), with 30 proteins with significant variation level among the treatments (**Figure 4I**). In both cases, the major differential phosphorylation hubs were Myc (UNIPROT ID: P01108), Sos1 (UNIPROT ID: Q62245), Braf (UNIPROT ID: P28028), Cacna1a (UNIPROT ID: P97445), Flna (UNIPROT ID: Q8BTM8), Nf1 (UNIPROT ID: Q04690), and Map3k1 (UNIPROT ID: P53349), which represented the most variable phosphorylation hubs (DN-Score > 5). Furthermore, analysis of the degree-corrected score identified Flnb (UNIPROT: Q80X90) and Rapgef2, while the previously mentioned Cacna1a remained among the most variable proteins across both analytical approaches. Cacna1a is a Voltage-Gated Calcium Channel (VGCC) frequently associated with cancer. Its abundance may be either upregulated or downregulated depending on the tumor type (**Phan et al., 2017**). The full results can be observed in **Table 3**. Finally, Figure 4J highlights the 30 genes with the highest number of significant phosphorylation events and their differential profiles across treatments.

**Table 3.** Proteins with the most variable phosphosite interactions between the FGF2 and SOSi/FGF2 treatments, identified using DyNet. The corresponding gene symbols are reported for each protein. The table includes raw DyNet rewiring scores, which quantify changes in phosphosite interactions, as well as degree-corrected rewiring scores.

| FGF2-FBS vs. SOSi/FGF2-FBS |  |  | FGF2+FBS vs. SOSi/FGF2+FBS |  |  |
|---|---|---|---|---|---|
| Gene | Dynet Rewiring (DN) | DN-Score (Degree Corrected) | Gene | Dynet Rewiring (DN) | DN-Score (Degree Corrected) |
| Cacna1a | 6,40 | 0,34 | Cacna1a | 6,49 | 0,34 |
| Flnb | 4,50 | 0,34 | Flnb | 4,50 | 0,34 |
| Rapgef2 | 3,50 | 0,31 | Rapgef2 | 3,50 | 0,31 |
| Flna | 6,40 | 0,30 | Flna | 6,49 | 0,30 |
| Daxx | 4,00 | 0,25 | Stmn1 | 2,99 | 0,29 |
| Taok1 | 2,50 | 0,22 | Daxx | 4,00 | 0,25 |
| Myc | 9,50 | 0,21 | Taok1 | 2,50 | 0,22 |
| Rps6ka4 | 2,90 | 0,19 | Myc | 9,50 | 0,21 |
| Sos1 | 8,00 | 0,18 | Rps6ka4 | 2,99 | 0,19 |
| Braf | 7,00 | 0,18 | Sos1 | 8,00 | 0,18 |
| Map3k1 | 5,50 | 0,18 | Braf | 7,00 | 0,18 |
| Rps6ka3 | 3,50 | 0,18 | Map3k1 | 5,50 | 0,18 |
| Epha2 | 3,50 | 0,18 | Epha2 | 3,50 | 0,18 |
| Nfkb2 | 2,90 | 0,18 | Rps6ka3 | 3,50 | 0,18 |
| Map4k4 | 2,00 | 0,18 | Nfkb2 | 2,99 | 0,18 |
| Nf1 | 5,90 | 0,17 | Map4k4 | 2,00 | 0,18 |
| Pak2 | 3,50 | 0,14 | Nf1 | 5,99 | 0,17 |
| Nfatc3 | 1,40 | 0,14 | Pak2 | 3,50 | 0,14 |
| Araf | 2,90 | 0,13 | Nfatc3 | 1,49 | 0,14 |
| Ptpn5 | 0,50 | 0,12 | Araf | 2,99 | 0,13 |
| Akt2 | 2,00 | 0,10 | Ptpn5 | 0,50 | 0,12 |
| Map3k2 | 1,00 | 0,09 | Akt2 | 2,00 | 0,10 |
| Map3k11 | 1,00 | 0,07 | Dusp8 | 0,50 | 0,10 |
| Rps6ka1 | 1,00 | 0,06 | Map3k12 | 0,50 | 0,10 |
| Pak1 | 0,50 | 0,02 | Map3k2 | 1,00 | 0,09 |
|  |  |  | Atf2 | 1,49 | 0,07 |
|  |  |  | Map3k11 | 1,00 | 0,07 |
|  |  |  | Rps6ka1 | 1,00 | 0,06 |
|  |  |  | Mapk11 | 1,00 | 0,04 |
|  |  |  | Pak1 | 0,50 | 0,02 |

Taken together, these results provide sufficient mechanistic evidence to support that Y1 cells rewire their signaling networks in a complex manner in response to treatments and perturbations, while stimuli such as fetal bovine serum can enhance the availability of paths, allowing different responses.

The top 30 KEGG/String MAPK (mmu04010) network can be accessed in **Supplemental Table 2**. The full-length KEGG/String MAPK (mmu04010) network can be accessed in **Supplemental Table 3**. All exclusive phosphorylations can be accessed in **Supplemental Figure S14** for FGF2-FBS, **Supplemental Figure S15** for SOSi/FGF2-FBS phosphorylations, **Supplemental Figure S16** for FGF2+FBS, and **Supplemental Figure S17** for SOSi/FGF2+FBS.

### Differential phosphorylation and kinase enrichment analysis

In the previous section, the main differential phosphorylations between treatments and time series were highlighted, providing a mechanistic clue of the differential signal propagation of the phosphosite signal. Finally, this section will demonstrate the results of differential phosphorylations by measuring the kinase activity. For this, the chosen tool was Kinase Library tools (**Johnson et al., 2023; Yaron-Barir et al., 2024**). This enrichment analysis uses the user-imputed limma log-transformed data and converts it to frequency factors (logFF), which quantify the over- or under-representation of kinase-specific motifs among differentially regulated phosphosite signatures. A positive logFF value indicates enrichment, whereas a negative value indicates depletion of the kinase activity. For this analysis, 311 Ser/Thr kinases and 13 Tyr kinases were differentially abundant (0.05% FDR significance threshold) in total. The enrichment score for each kinase in all conditions is displayed in the **Supplemental Figure S18** and the **Supplemental Table S4** and **Supplemental Table S5**. The most relevant kinase activities are presented in modules below.

The RAS → RAF→ MEK→ ERK axis module (**Figure 5A**) revealed that SOS1 inhibition could not prevent ERK activation, but raised its levels. For instance, the level of kinase activation for both ERK1 and ERK2 in the condition SOSi/FGF2-FBS 30 minutes is Log2FF 0,67x and 0,65x, respectively, in contrast to the Log2FF 0,33x and 0,34x for the condition FGF2-FBS 30 minutes. This suggests that the SOS1 inhibition, combined with the FGF2 stimulation, relieves the feedback effect caused by BI-3406. Next, the most relevant kinases outside the canonical MAPK axis are discussed, in order to unveil the mechanism behind the perturbations performed in this study.

**Figure 5:**
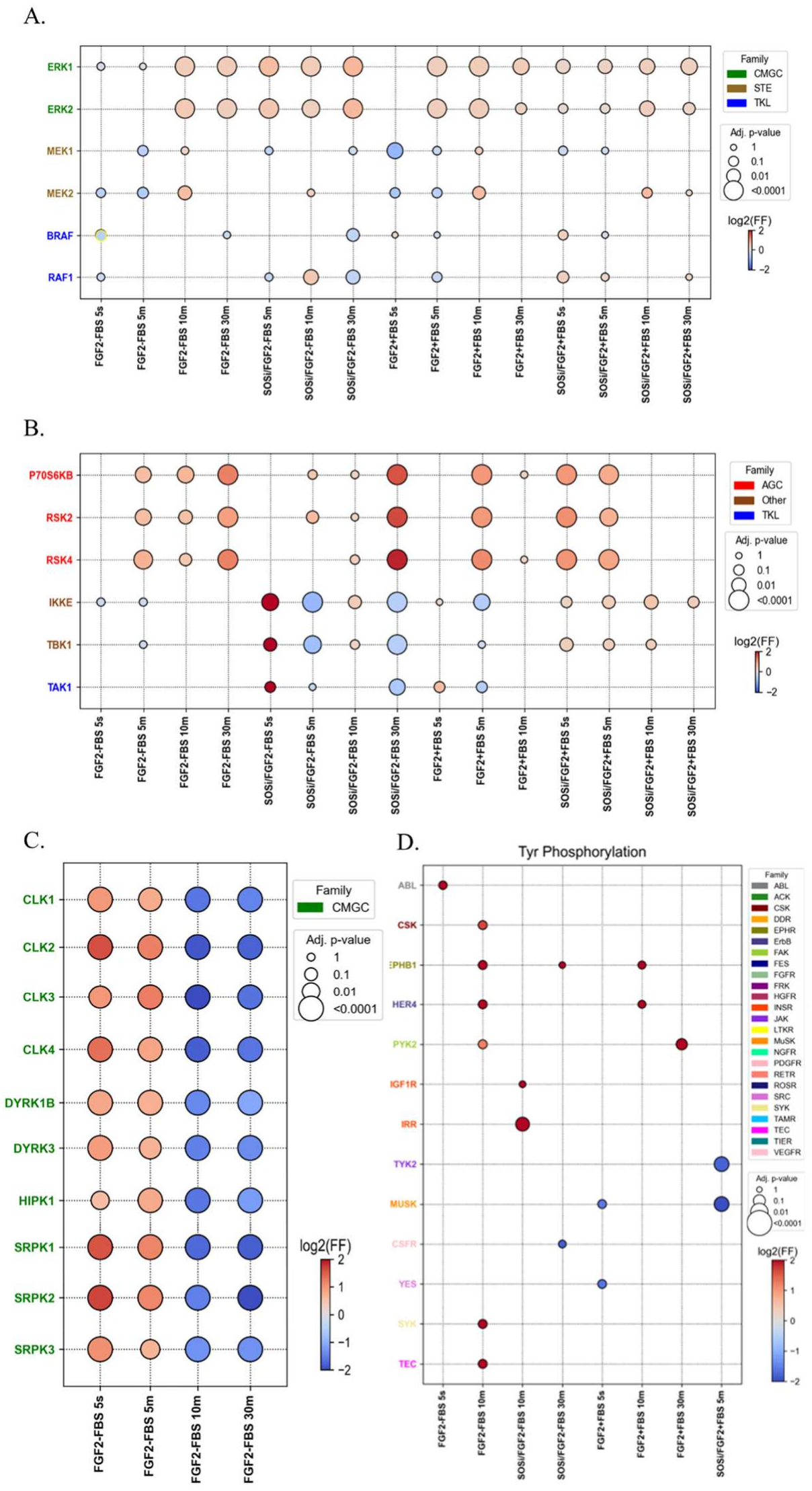
Differential phosphorylation and kinase enrichment analysis. Differentially phosphorylated tyrosine sites for each Limma contrast were identified using the DiffPhosData class from the kinase_library Python package (Johnson et al., 2023; Yaron-Barir et al., 2024). Sites with adjusted p-value < 0.05 were analyzed using log fold-change values and phosphosite sequence windows. Kinase enrichment was inferred using percentile-rank scoring (threshold = 15 for Ser/Thr and 8 for Tyr). When the treatment does not appear in a specific time-point in a module, there is no statistical significance, therefore not shown. **A:** Kinase Activity of the RAS → RAF→ MEK→ ERK axis members module. **B:** Kinase Activity of Mtor and Ras-Ralb members module. **C:** Kinase Activity of exclusively activated members for the FGF2-FBS 5s - 5m. **D:** Tyr kinase activity. Legend: Red: Upregulated phosphorylation events. Blue: blue indicates downregulated events. The individual graphics and the source table for this result can be seen on **Supplemental Figures S19-26** and **Supplemental Table S4** for Ser/The residues and **Supplemental Figures S27-34** for and **Supplemental Table S5** Tyr residues.

The module containing the contrasting kinase activity between the SOSi/FGF2 and FGF2-only treatments (**Figure 5B**) pointed to intermittent stress signaling, followed by MAPK parallel activation that sustained ERK levels. The SOSi/FGF2+FBS 5 seconds condition showed upregulation of IKKε (log2FF ~2.2x), TAK1 (log2FF ~2.4x), and TBK1 (log2FF ~2.0x) kinases activity, which correlated with inflammatory processes. TBK1 has been described as a critical effector of the Ras–RalB pathway that sustains cell survival in Ras-driven contexts (Chien et al., 2006; Barbie et al., 2009), while TAK1 and IKKε are regulated by ERK and their coabundance and upregulation are related to KRAS oncogenic phenotypes (Graves et al., 2025). Interestingly, the effect of FGF2 for 5 minutes was sufficient to reverse this effect, restoring the canonical Ras–ERK activation pattern and concomitantly reducing the phosphorylation of IKKε (log2FF ~-0,14x), TAK1 (log2FF ~-0,37x), and TBK1 (log2FF ~-0,77x). This reciprocal behavior highlights the dynamic interplay between the SOS–Ras axis and stress-activated kinases in regulating the balance between mitogenic and inflammatory signaling.

In addition, increased phosphorylation of p70S6K, RSK2, and RSK4 was observed in the SOSi/FGF2-FBS 30-minute condition. p70S6K (log2FF ~1.6x) is a kinase activated downstream of the mTOR signaling cascade, which regulates cell growth and protein synthesis (Magnuson, Ekim, and Fingar, 2012). The RSK family members RSK2 (log2FF ~1.7x) and RSK4 (log2FF ~1.3x) are downstream effectors of the RAS–RAF–MEK–ERK pathway, which mediates the mitogen-induced transcriptional and translational responses (Frödin and Gammeltoft, 1999; Romeo, Zhang, and Roux, 2012). In addition, the FGF2-related module (**Figure 5C**) revealed kinases that were exclusively activated under FGF2-FBS conditions at 5 seconds and 5 minutes of FGF2 stimulation. These conditions revealed CLK1-4, DRYK1B, DYRK3, HIPK1 and SRPK1-3 kinases upregulation. From those mentioned, the only one previously reported and confirmed with experimental confidence was the SRPK1 kinase, which was found to be activated by FGF2-driven angiogenic signaling (Jia et al., 2021). All the aforementioned kinases, to the best of our knowledge, are novel in the FGF2 signaling context and warrant further validation.

Lastly, in the Tyr kinase activation module (**Figure 5D**), a limited number of kinases with significant scores could be assessed. Once none of the kinase activities were identified in more than three conditions, a comparative analysis could not be performed. Nevertheless, it is worth mentioning that IRR (insulin receptor-related receptor) was significantly increased in the SOSi/FGF2-FBS 10 minutes condition (log2FF ~3.8). The IRR pathway was already pointed out as a canonical MAPK pathway activator, such as proliferation in hepatocellular carcinoma through IRR-mediated MAPK activation shown by Wang et al. (**Wang et al., 2013**). Taken together, these results hint at the compensatory activation of the MAPK cascade through alternative paths in response to SOS1 inhibition, where the ERK levels are sustained by Ras-Ralb and mTOR pathways, supported by stress-mediated kinases such as IKKε, TAK1, and TBK1.

## Discussion

Phosphorylation is an indispensable cell signaling regulation mechanism, which is involved in many cellular processes, ranging from signal transduction and cell growth to pathway activation and deactivation (**Hornberg et al., 2005; Ardito et al., 2017**). The study of phosphorylation can reveal critical regulatory mechanisms in cancer (**Blume-Jensen and Hunter, 2001**) that are undetectable at the global proteome level (**Drake et al., 2016**), uncovering an additional layer of signaling complexity.

One of the most well-studied phosphorylation cascades is the MAPK cascade, whose main axis comprises RAS → RAF → MEK → ERK signaling (**Seger & Krebs, 1995**). In the absence of RAS mutations, this pathway is initiated by SOS-mediated RAS activation, which recruits adaptor proteins that promote sequential phosphorylation of RAF, MEK, and ERK to enable a plethora of cellular events (**Seger and Krebs, 1995**). Despite years of characterization, our most recent study revealed that additional GEF control is necessary to sustain the GEF-mediated activation of non-mutated KRAS that confers the tumor phenotype in Y1 cells (**Montoni et al., 2026**).

Since GEFs and GAPs are molecular switches capable of activating or deactivating G-proteins such as Ras (**Bos, Rehmann e Wittinghofer, 2007**), consequently regulating multiple phosphorylation events in the cell, this supports the premise that additional unknown GEF/GAP regulatory mechanisms may underlie several unexplored signaling responses. Based on that, this paper explored the proteomic and phosphoproteomic molecular dynamics of the Y1 cell in front of FGF2, a sensitizing factor for this cell (**Costa et al., 2008; Salotti et al., 2013; Dias et al., 2019**), and FGF2 plus SOS inhibition, to expose compensatory/adaptive signaling without one of its main activators (**Chardin et al., 1993; Mita et al., 2009**), with or without FBS presence in a 0 to 30-minutes time-span.

The analysis revealed a stable proteome in the short experimental time span, where only ≈ 1.26% (111/8,786) of the protein abundance significantly changed. Among them, MAPK cascade-related proteins were found, such as JUNB, which has a context-dependent effect on tumorigenesis (**Li et al., 2023**), and SERPINE1/PAI-1, a JUNB downstream signaling member (**Shaikh et al., 2024**). Both were found upregulated under SOSi 30-minute treatments (**Figure 2C**). In contrast, ≈ 72.2% (10,416/14,512) of the phosphosites were differentially abundant across the conditions, revealing broader phosphoregulatory dynamics compared to the proteome (**Figure 2D**).

Next, the GEF and GAP phosphoproteomic abundance assessment revealed that all the GEFs and GAPs identified had stable proteomic levels (**Figure 3A**). In contrast, several classes of GEFs and GAPs were altered across the experimental conditions and time series at the phosphoproteomic level (**Figure 3B and 3C**). RHO GEFs displayed the highest number of regulated phosphosites across the treatments, particularly under SOSi + FGF2 conditions, suggesting adaptive rewiring beyond canonical RAS signaling. In parallel, RAL GAPs also exhibited dynamic phosphoregulation across the treatments, reinforcing the participation of alternative small GTPase signaling branches in the Y1 adaptive response.

In sequence, the SOS1 dynamics exhibited time- and context-dependent phosphorylations (**Figure 4A**). The FGF2 treatment displayed a broader spectrum of phosphorylation, with four exclusively significant phosphosites. It was also revealed that SOS inhibition without FBS stimulation suppressed almost all the phosphorylations, with the exception of the sites S1147 and S1153, which were kept in all conditions and almost all time points, possibly being a hub for feedback effect relief. The serum availability allowed the phosphorylation sites to be recovered in SOS inhibition plus FBS stimulation and even enabled the exclusive site S1196. This is expected, once the FBS contains several growth factors (**Lee et al., 2022**) and their presence is known to play a role in counteracting the effect of inhibitors (**Koch et al., 2016**). All the sites are located in the proline-rich region of the SOS1 protein (a.a. 1141–1300), which is an IDR, known for ERK feedback effects and pathway rewiring (**Kamioka et al., 2010**). The mechanistic evaluation revealed that a large proportion of the MAPK phosphoproteome rewired its pattern across the conditions (**Figure 4A-J**). Notably, MYC, SOS1, BRAF, NF1, MAP3K1, and CACNA1a showed highly adaptive signaling that extends beyond the canonical ERK propagation.

Finally, kinase activity analysis revealed the functional consequences of the mechanistic differences observed across treatments and time points (**Figure 5)**. Despite the phosphoproteomic change observed at the identification and mechanistic level of the RAS → RAF→ MEK→ ERK axis, the kinase activity of the ERK module did not display inhibition, but upregulation, including the SOS-inhibited modules (**Figure 5A**). When opening the scope, it can be observed that in the SOSi/FGF2+FBS treatment, stress-related kinases from the Ras-RalB pathway, TBK1, TAK1, and IKKε were particularly upregulated (**Figure 5B**), supporting the previous premise that parallel regulation of phosphorylation sustains ERK levels. Also, the SOSi/FGF2-FBS 30-minute condition showed increased mTOR-associated p70S6K and ERK downstream kinases RSK2 and RSK4 activation. Additionally, the FGF2-related module (**Figure 5C**) revealed exclusive activation of kinases such as SRPK1 and DYRK3 under FGF2-FBS conditions, which are, to the best of our knowledge, unreported in FGF2 signaling. Together, these results suggest that SOS inhibition promotes adaptive signaling rewiring through alternatives, such as Ras-RalB, stress-response, and mTOR-associated pathways to sustain ERK signaling. Together, these findings provide a systems-level view of dynamic signaling adaptation in Y1 cells and establish a foundation for understanding sensitization mechanisms in RAS-dependent tumors. Our current working hypothesis, integrating the kinase activity results discussed above, is summarized in **Figure 6**.

**Figure 6:**
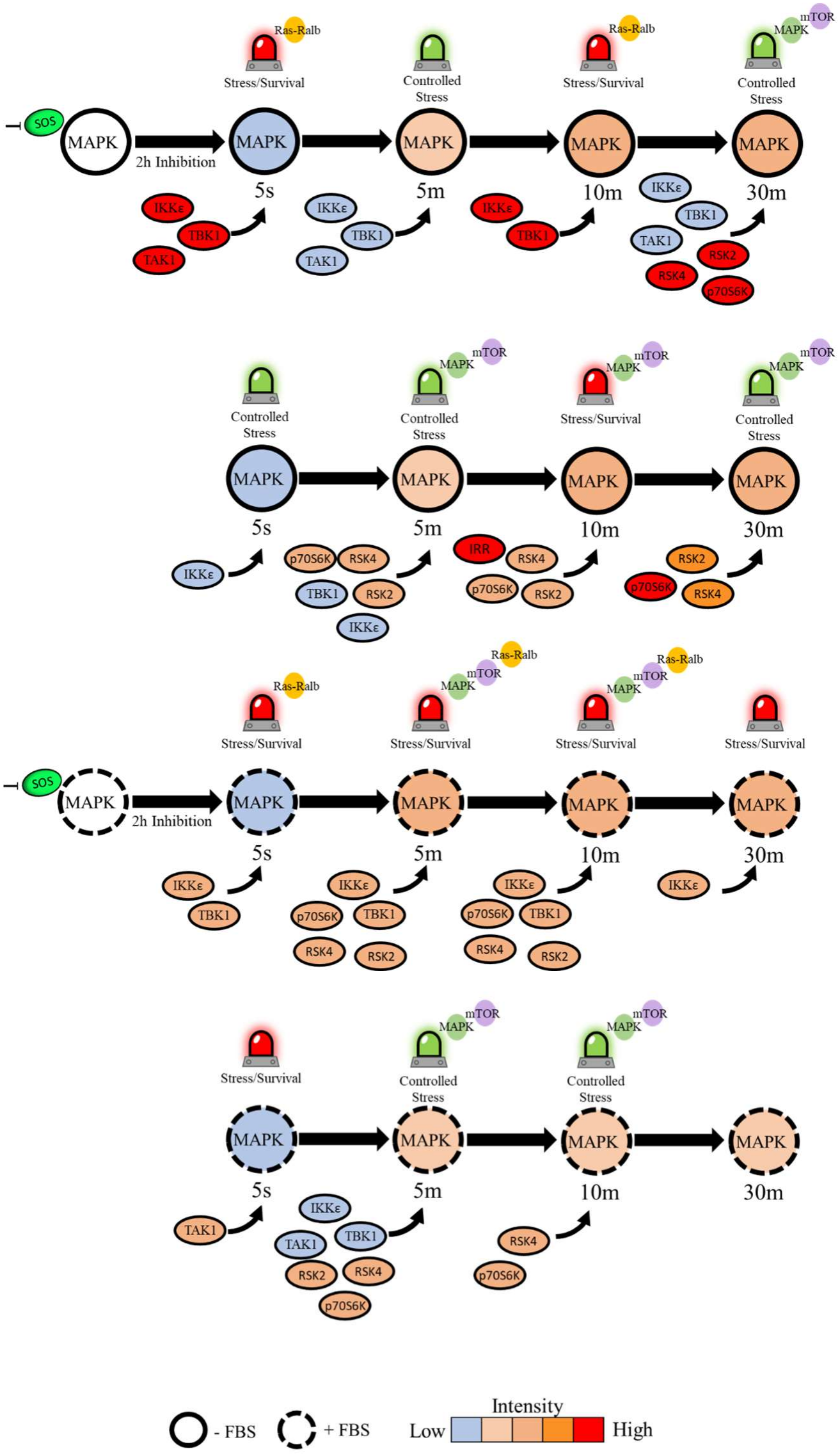
Integrated Overview of Kinase Activity. This schematic summarizes the differential kinase activity observed across FGF2 treatment conditions and time points, highlighting the central roles of MAPK signaling and SOS inhibition in the proposed regulation of the balance between cell survival and cell death in Y1 cells. Color intensity, ranging from blue (reduced activity) to red (increased activity), follows the same patterns of activity represented in **Figure 5**.

## Acknowledgements

We thank Hugo A. Armelin for insightful discussions and valuable intellectual contributions throughout the development of this work. We also gratefully acknowledge financial support from the National Institutes of Biomedical Innovation, Health and Nutrition (NIBN), the São Paulo Research Foundation (FAPESP; Grants #13/07467-1, #20/10329-3, #19/21619-5, #20/08555-5, #19/24580-2, and #21/04355-4), the Coordination for theImprovement of Higher Education Personnel (CAPES; Finance Code 001), and the Butantan Institute. This paper was typeset with the bioRxiv word template by @Chrelli: www.github.com/chrelli/bioRxiv-word-template.

## Author contributions

**F.M**.: Conceptualization, Methodology, Investigation, Formal Analysis, Data Curation, Visualization, Writing – Original Draft, Writing – Review & Editing; **T.H**.: Methodology, Investigation; **D.K**.: Investigation, Formal Analysis; **R.A.M.W**.: Conceptualization, Methodology, Investigation, Validation; **K.A.M.T**.: Methodology, Validation; **T.E.P.T**.: Methodology, Validation; **J.A**.: Resources, Supervision, Writing – Review & Editing, Funding Acquisition; **M.S.R**.: Conceptualization, Supervision, Writing – Review & Editing; **Y.N.K**.: Supervision, Writing – Review & Editing.

The authors **J.A., M.S.R**. and **Y.N.K**. jointly supervised this work.

## Materials and Methods

### Experimental Design

For each sample, 1×10^6^ Y1 cells were plated. The next day, synchronization in the G0/G1 phase was performed by serum starvation using FBS-free DMEM for 48 h. The experimental conditions were: (1) Control group, without stimulation; (2) 10 ng/mL Fibroblast Growth Factor 2 (FGF2) (in-house preparation, according to Costa et al. (**Costa et al., 2008**) and (3) 0.5 nM SOS1 inhibitor BI-3406 (SOSi) combined with 10 ng/mL FGF2. Each condition was evaluated in the presence or absence of FBS. Samples were collected at 0, 5, 10, and 30 min for the control group, and at 5 s, 5, 10, and 30 min for the FGF2 and SOSi + FGF2 groups, in triplicates.

### Cell Culture

The Y1 murine adrenocortical carcinoma cell line was obtained from ATCC and cultured under standard conditions in Dulbecco’s Modified Eagle Medium supplemented with 10% fetal bovine serum and 1% penicillin/streptomycin at 37 °C under a humidified 5% CO_2_ atmosphere. Cell counting was performed using the Countess™ system according to the manufacturer’s instructions.

### Protein Extraction and Sample Preparation

Following stimulation, cells were harvested, and proteins were extracted using a PTS-based lysis workflow containing phosphatase and protease inhibitors. Protein concentration was determined using the DC Protein Assay™ Kit II according to the manufacturer’s instructions. Protein digestion was performed using the automated SP3 workflow as previously described (**Muraoka et al., 2022**). After desalting, peptide samples were separated for global proteome and phosphoproteome analyses. Phosphopeptides were enriched using the Fe-IMAC StageTip workflow (**Adachi et al., 2022; Gunji et al., 2023**), followed by Phospho-TMT labeling, cleanup, and high-pH reversed-phase fractionation according to previously described methodologies (**Adachi et al., 2016; Gunji et al., 2024; Shoji et al., 2024**).

### Mass Spectrometry Analysis

Global proteome samples were analyzed by DIA LC-MS/MS using a Q Exactive™ mass spectrometer coupled to an UltiMate™ 3000 NanoLC system. GPF-DIA acquisition was performed for spectral library generation. Phosphoproteome samples were analyzed by DDA LC-MS/MS using an Orbitrap Fusion™ Lumos™ Tribrid™ mass spectrometer coupled to an UltiMate™ 3000 NanoLC system.

### Proteomic and Phosphoproteomic Data Processing

Phospho-TMT samples were analyzed using MaxQuant (v2.6.2.0) (**Tyanova, Temu e Cox, 2016**) against the Mus musculus SwissProt data-base. Phosphorylation of serine, threonine, and tyrosine residues was considered for phosphosite identification, with identification confidence controlled by false discovery rate filtering as described previously. DIA proteome samples were processed using DIA-NN (v1.8.2) (**Demichev et al., 2020**) using the GPF-derived spectral library and the same reference database.

### Data Preprocessing and Differential Analysis

Raw data tables were filtered to remove decoy sequences, potential contaminants, and features with more than 30% missing or zero values across samples. Remaining values were log_2_-transformed, and missing values were imputed using Bayesian PCA with automatic relevance determination (BPCA-ARD) (**Oba et al., 2003; Kitamoto et al., 2023**). Batch correction was performed using the ComBat algorithm implemented in pycombat (**Johnson, Li, and Rabinovic, 2007; Behdenna et al., 2023**), followed by mean centering using StandardScaler (**Pedregosa et al., 2011**). PCA and UMAP analyses were performed using the scikit-learn (**Pedregosa et al., 2011**) and umap-learn (**McInnes, Healy, and Melville, 2018**) implementations, respectively.

Differential abundance analysis was performed using the limma package in R (**Ritchie et al., 2015**). Treated and control groups were compared at each time point using predefined contrasts, as described in **Table 1**. Features with an absolute log_2_ fold change greater than log_2_(1.5) and an adjusted p-value below 0.05 were considered significantly regulated.

For detailed experimental procedures, data normalization, and technical validation, please refer to the companion paper preprint Montoni, F. (2026c) and the doctorate thesis Montoni, F. (2026b).

### Differential phosphorylation and kinase enrichment

Differentially regulated phosphosites were identified for each limma contrast (adjusted p < 0.05) and subjected to kinase enrichment analysis using the Kinase Library framework (**Johnson et al., 2023; Yaron-Barir et al., 2024**). Enrichment was computed from log fold change values and phosphosite sequence windows, applying percentile-rank thresholds of 15 for Ser/Thr and 8 for Tyr sites. Enrichment results were visualized as bubble plots.

### Kinase Network and GEFs and GAPs Enrichment

To construct the interaction network between kinases, GAPs, and GEFs, a gene-centric analysis was performed using KiNet (**Sekar et al., 2024**), which generated networks based on the input genes. GAP and GEF genes were associated with kinases for which they have been reported as potential substrates in the literature. In parallel, over-representation analysis (ORA) of GAP and GEF gene sets was conducted using WebGestalt (**Elizarraras et al., 2024**) with all mapped Entrez Gene IDs from the selected genome protein-coding reference list as background. Analysis parameters included a minimum category size of 5 genes, maximum category size of 2000 genes, Benjamini–Hochberg (BH) false discovery rate correction, and significance level set to the top 10 enriched categories. Network visualization was performed in Cytoscape (**Shannon et al., 2003**).

### Phosphosite Network Analysis

Significant phosphosites identified by limma for each treatment were analyzed using the PTM Navigator tool (**Müller et al., 2025**) available in ProteomicsDB (https://www.proteomicsdb.org/) (**Schmidt et al., 2018**). Relevant phosphosites associated with the KEGG (**Kanehisa et al., 2017**) MAPK signaling pathway (mmu04010) were extracted and annotated. Protein–protein interaction networks were subsequently generated in STRING (**Szklarczyk et al., 2024**) using the top 30 proteins containing significant phosphosites (P < 0.05) and visualized in Cytoscape. The differential network analysis was performed using the Cytoscape plugin DyNet (**Salamon, Goenawan, and Lynn, 2018**).

### Data visualization

Plots and graphical summaries were generated using matplotlib (**Hunter, 2007**) and seaborn (**Waskom, 2021**), with the scienceplots style (**Garrett, 2021**) applied. Heatmaps were generated using the Morpheus web tool (https://software.broadinstitute.org/morpheus/).

### Data availability

For this preprint version only the main text and supplementary figures are available. If you have interest in seeing the supplementary tables, raw data, processed tables and other information, please wait for the publication.

