## Supplementary Information for "Beyond canonical: Hidden phosphoregulatory control in RAS signaling reveals context-dependent rewiring in Y1 tumor cells"

    - [1] Center of Toxins, Immune-Response and Cell Signaling (CeTICS), Butantan Institute, São Paulo, 05503-900, Brazil.
    - [2] Cell Cycle Laboratory, Butantan Institute, São Paulo, 05503-900, Brazil.
    - [3] Bioinformatics Graduate Program, University of São Paulo, Brazil
    - [4] Laboratory of Artificial Intelligence and Inference in Complex Data (Recod.ai), Institute of Computing, University of Campinas, Campinas, 13083-852, Brazil.
    - [5] AI Center for Health and Biomedical Research (ArCHER), National Institutes of Biomedical Innovation, Health and Nutrition, Osaka, 566-0002, Japan.
    - [6] Laboratory of Proteomics for Drug Discovery, Center for Drug Design Research (CDDR), National Institutes of Biomedical Innovation, Health and Nutrition, Ibaraki, 567-0085, Japan.
    - [‡] Current affiliation: Laboratory of Computational Biomedicine, Graduate School of Science, Technology and Innovation, Kobe University, Kobe, 650-0017, Japan.
    - [§] Current affiliation: Department of Mechanical and System Engineering, Graduate School of Engineering, University of Fukui, Fukui, 910-8507, Japan.
    - * Correspondence Author
    - † These authors jointly supervised this work

- **Keywords**: Phosphoproteomics, Y1 Adrenocortical Cell, Cancer Research, Cell Signaling, Mass Spectrometry

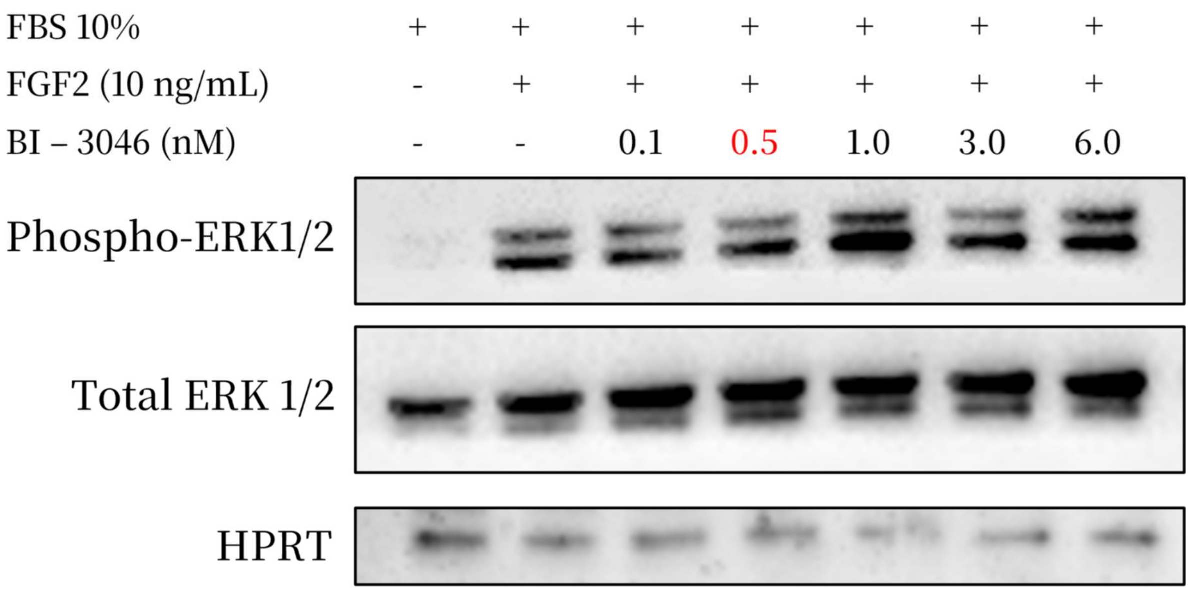


**Figure S1:** Western blot analysis of SOSi concentrations. The phosphorylated ERK 1/2 was observed after 2 hours of BI-3406 stimulation. The concentration used for the downstream applications was 0.5 nM, which was the smaller concentration that exhibited higher inhibition of the pERK. Some of the higher concentrations such as 1.0 nM and 6.0 nM, showed higher pERK activity, being an indication of feedback effect from the SOS inhibition.

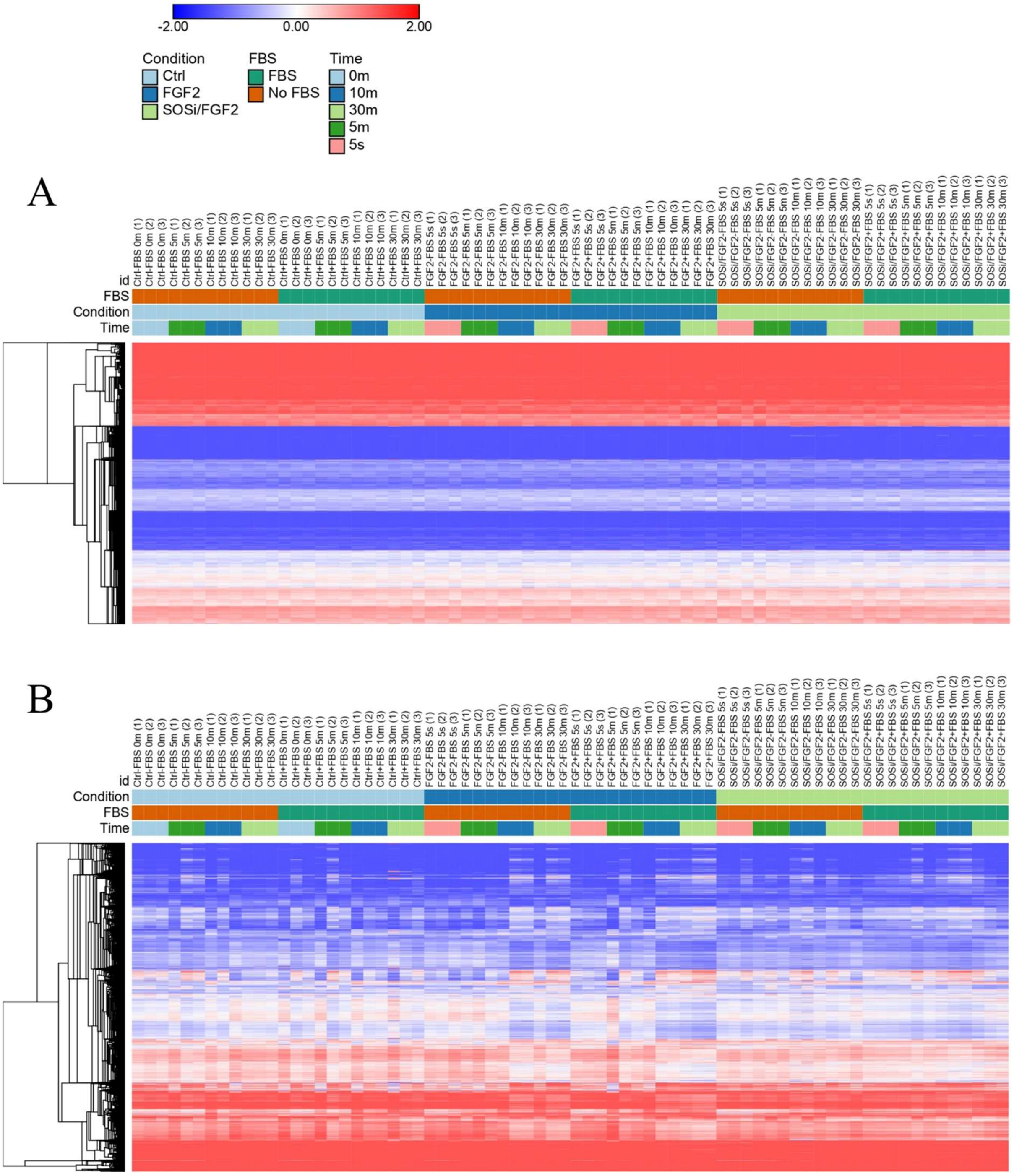


**Figure S2**: Hierarchical clustering analysis using Euclidean distance prior to differential abundance analysis. **A.** Normal Proteome (N=8,786). **B.** Phosphoproteome (N=14,512).

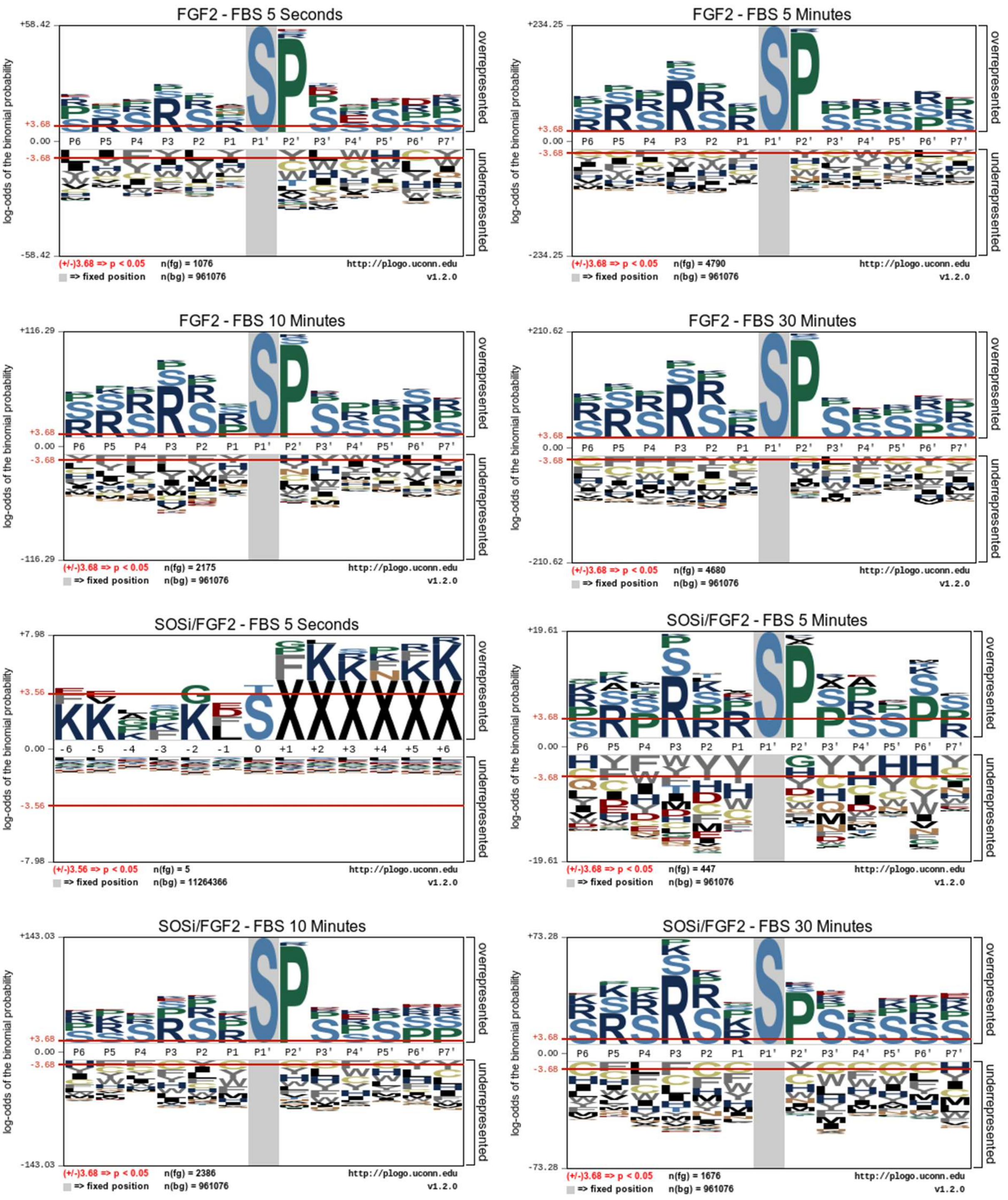


**Figure S3**: IceLogo analysis of phosphosites significantly regulated in minus FBS conditions. Amino acid frequencies were calculated within a ±6 residue window centered on the phosphorylated S/T site. The logo displays residues significantly over- or under-represented compared to the background distribution limma significant sites.

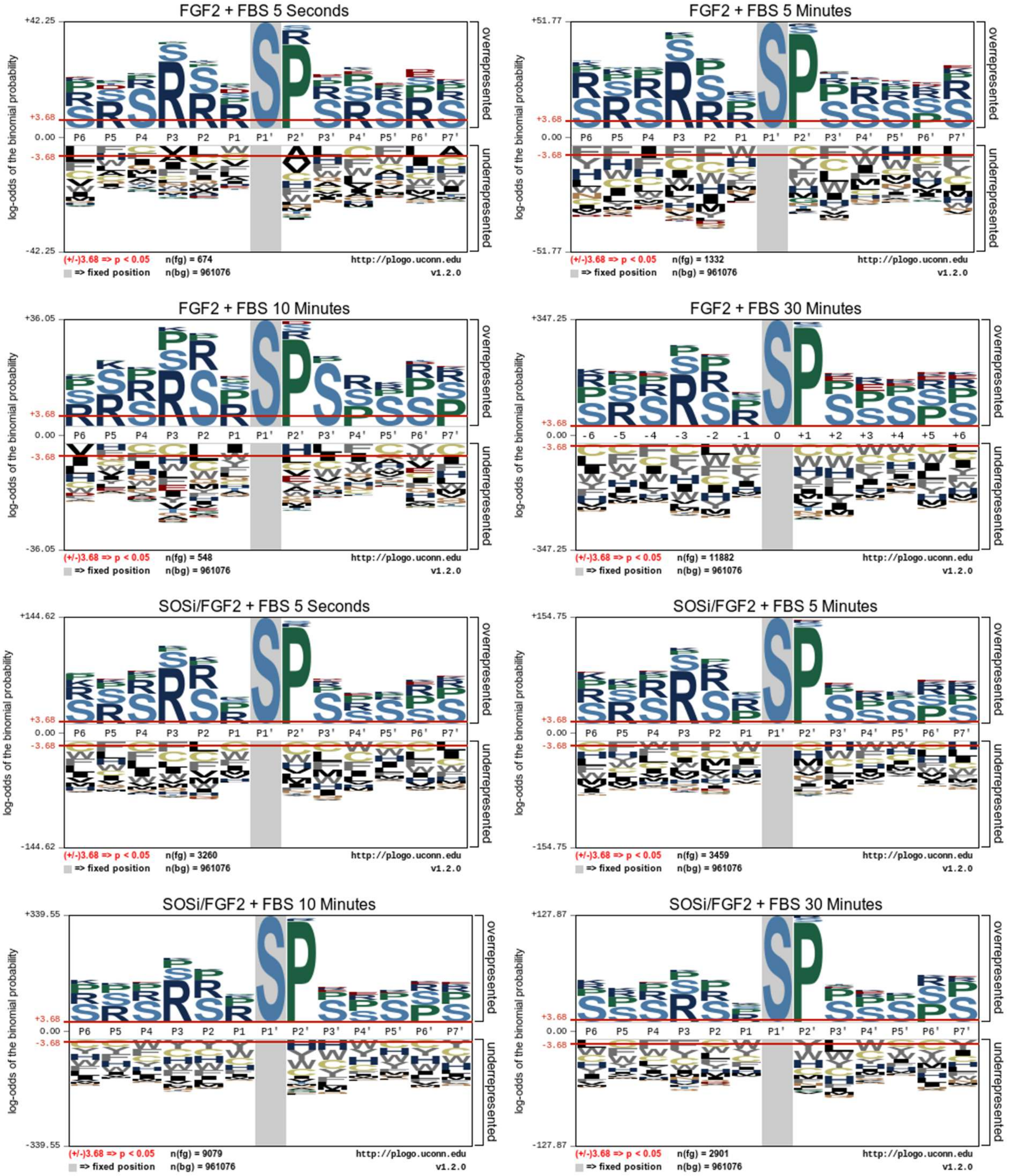


**Figure S4**: IceLogo analysis of phosphosites significantly regulated in plus FBS conditions. Amino acid frequencies were calculated within a ±6 residue window centered on the phosphorylated S/T site. The logo displays residues significantly over- or under-represented compared to the background distribution limma significant sites.

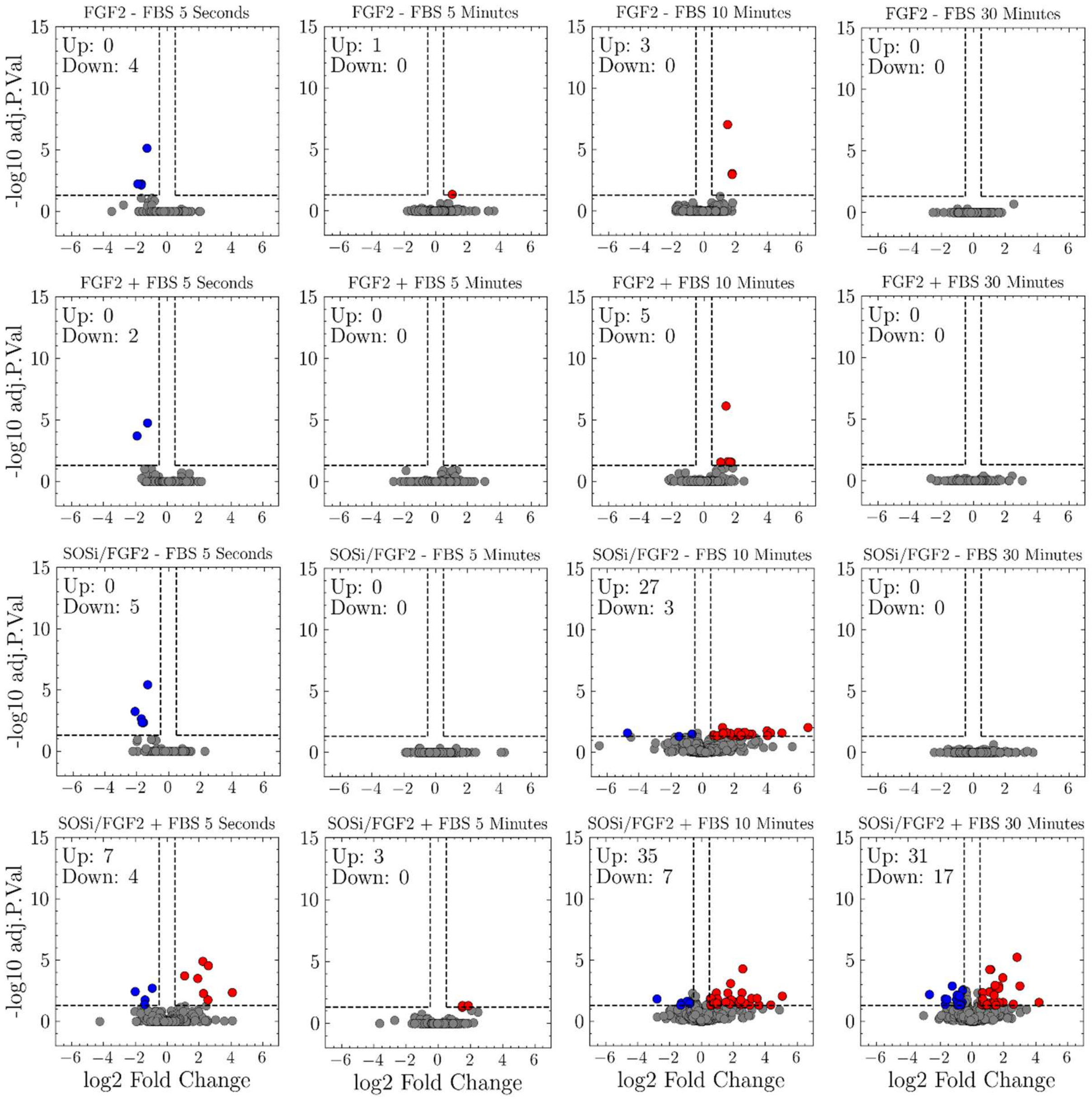


**Figure S5**: Volcano plot of protein abundance changes. The x-axis represents log2 fold change, and the y-axis shows − log10 (Adjusted p-value) limma analysis. Proteins with an absolute log2 fold change ≥ 0.585 (corresponding to a 1.5-fold change) and an adjusted p-value ≤ 0.05 ( − log10 ≈ 1.3) are considered significantly differentially abundant. Proteins highlighted in blue are downregulated, while proteins in red are upregulated.

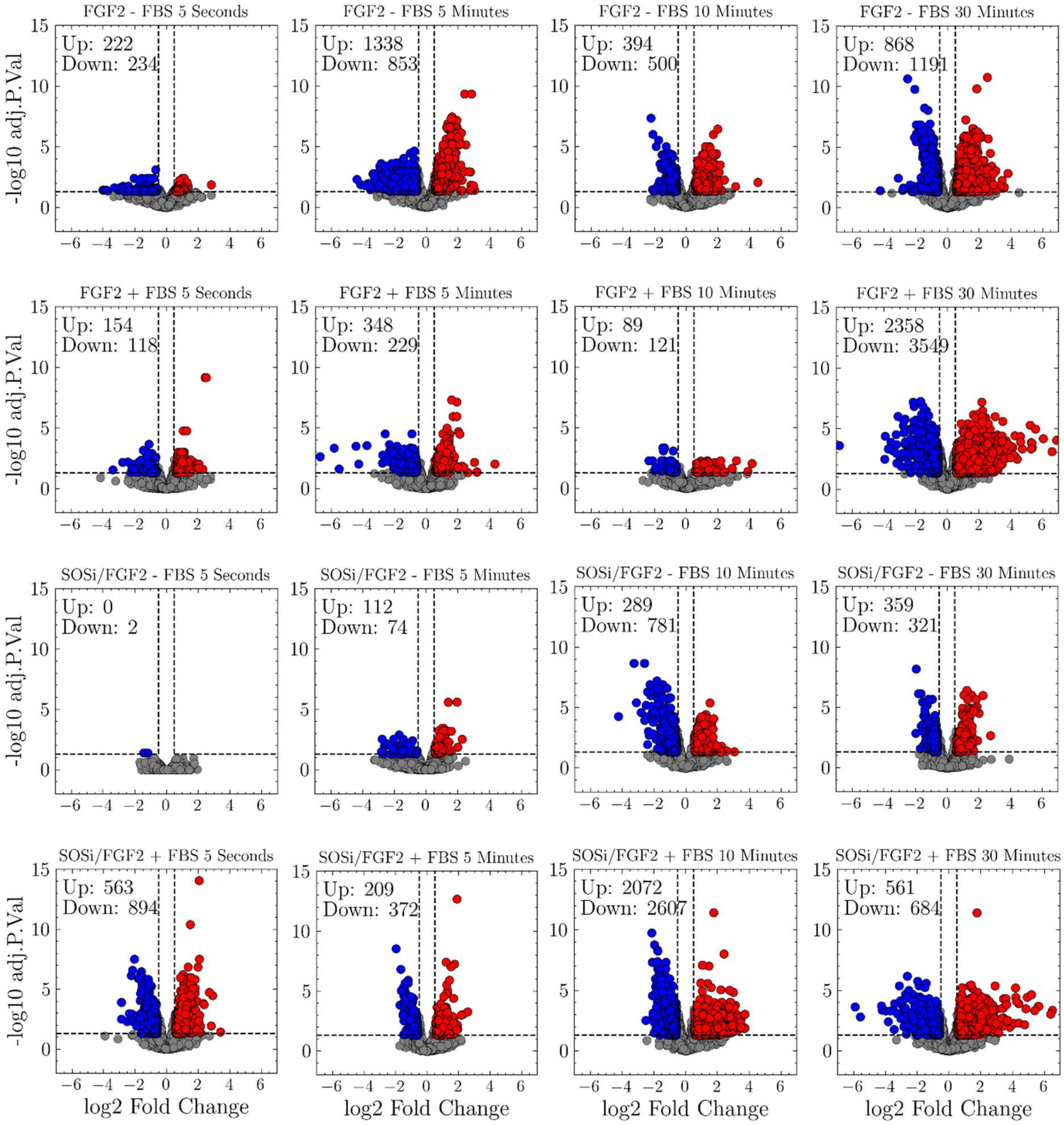


**Figure S6**: Volcano plot of phosphosite abundance changes. The x-axis represents log2 fold change, and the y-axis shows − log10 (Adjusted p-value) limma analysis. Proteins with an absolute log2 fold change ≥ 0.585 (corresponding to a 1.5-fold change) and an adjusted p-value ≤ 0.05 ( − log10 ≈ 1.3) are considered significantly differentially abundant. Proteins highlighted in blue are downregulated, while proteins in red are upregulated

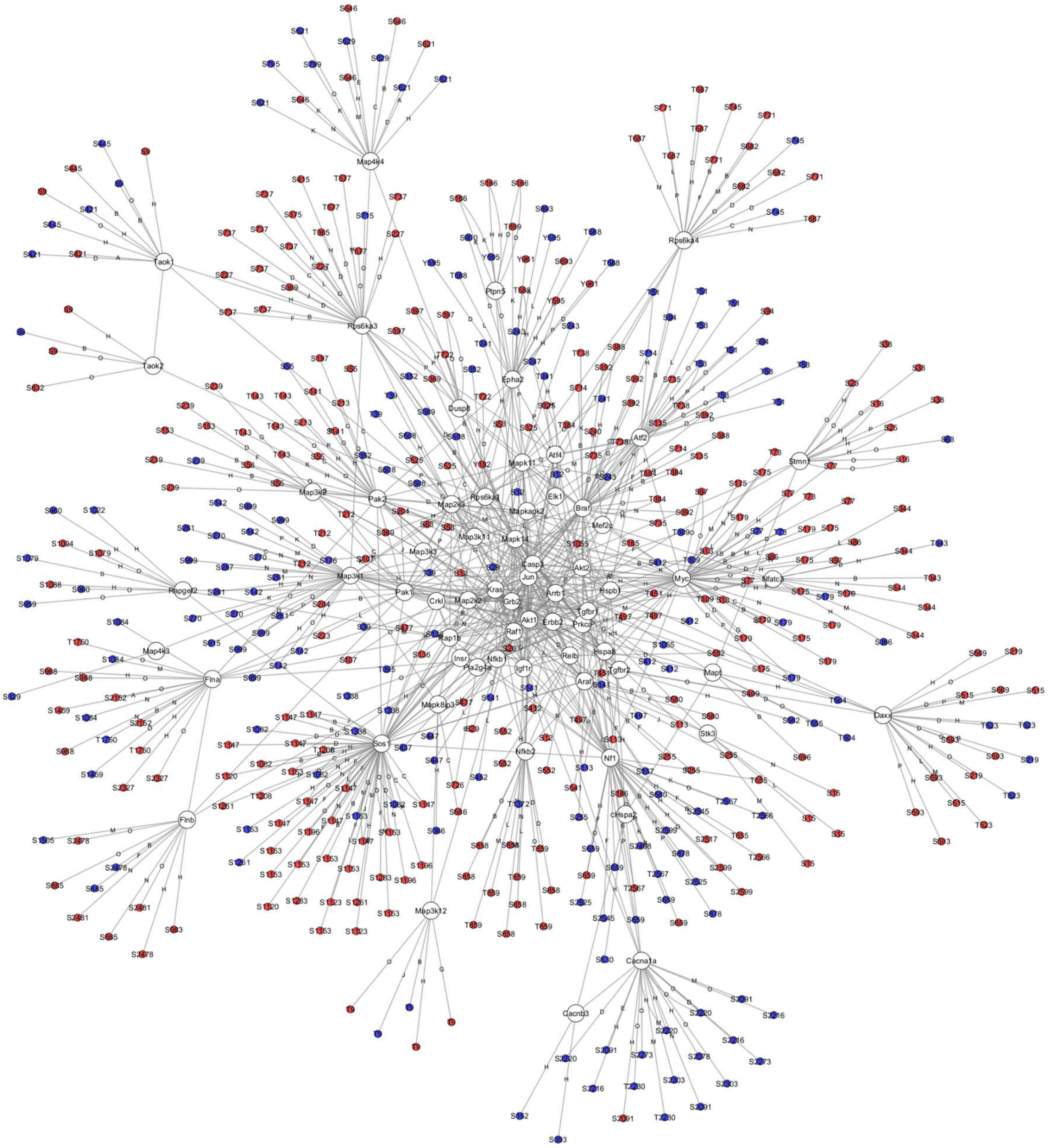

**Figure S7**: Protein interaction network generated from the relevant phosphosites identified by LIMMA analysis and subsequently uploaded to the ProteomicsDB PTM Navigator. Phosphosites were enriched for the MAPK pathway (mmu04010) and then selected (pValue = 0.05 <) to reconstruct the interaction network using a combination of STRING and Cytoscape tools. **Legend**: **White Node**: Proteins. **Red node**: Upregulated phosphosite. **Blue node:** Downregulated phosphosite. **Edges** **A**: FGF2 - FBS 5 seconds; **B**: FGF2 - FBS 5 minutes; **C**: FGF2 - FBS 10 minutes; **D**: FGF2 - FBS 30 minutes; **E**: FGF2 + FBS 5 seconds; **F**: FGF2 + FBS 5 minutes; **G**: FGF2 + FBS 10 minutes; **H**: FGF2 + FBS 30 minutes; **I**: SOSi/FGF2 - FBS 5 seconds; **J**: SOSi/FGF2 - FBS 5 minutes; **K**: SOSi/FGF2 - FBS 10 minutes; **L**: SOSi/FGF2 - FBS 30 minutes; **M**: SOSi/FGF2 + FBS 5 seconds; **N**: SOSi/FGF2 + FBS 5 minutes; **O**: SOSi/FGF2 + FBS 10 minutes; **P**: SOSi/FGF2 + FBS 30 minutes.

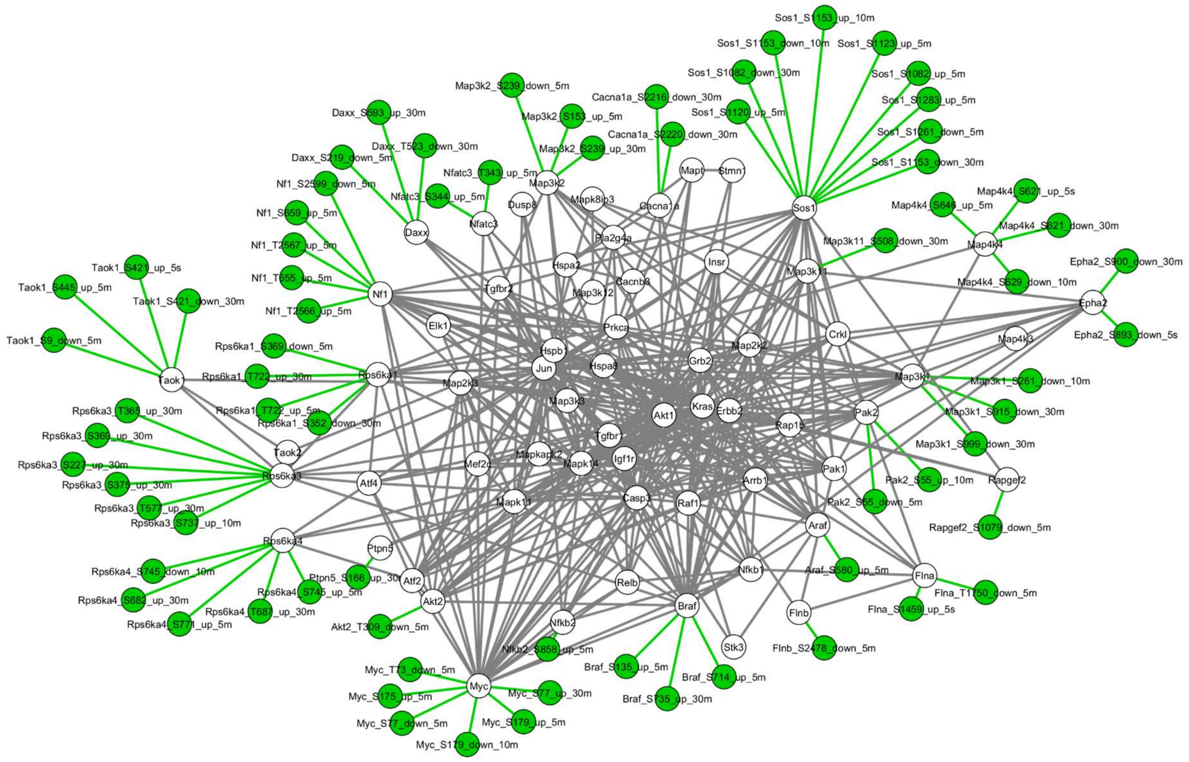


**Figure S8:** Top 30 proteins with the highest number of phosphorylation sites and $p < 0.05$ (Limma analysis) network was compared using DyNet for the FGF2-FBS versus SOSi/FGF2-FBS treatment. Phosphorylation sites exclusive to the FGF2-FBS treatment are highlighted in green. In total, 72 sites were exclusively identified across treatments and time points.

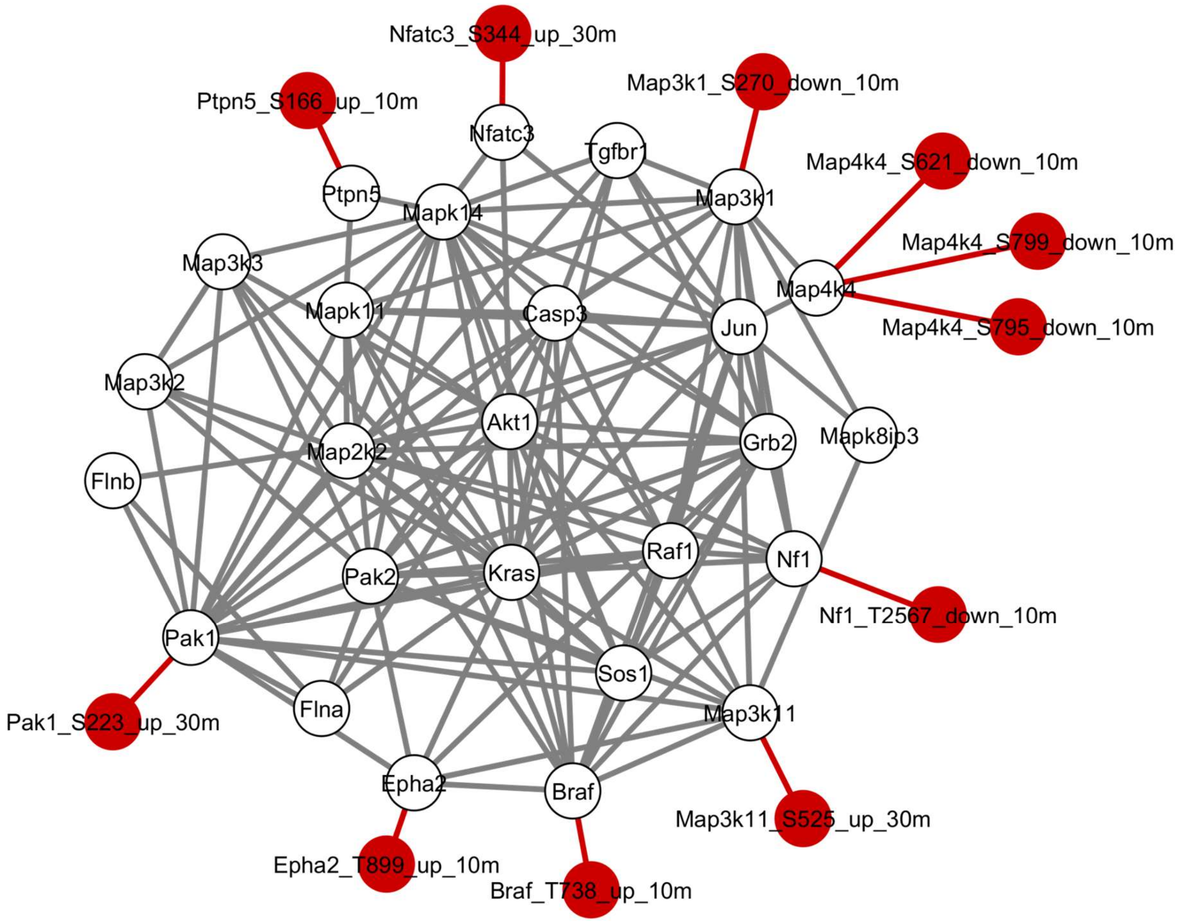


**Figure S9:** Top 30 proteins with the highest number of phosphorylation sites and $p < 0.05$ (Limma analysis) network was compared using DyNet for the SOSi/FGF2-FBS versus FGF2-FBS treatment. Phosphorylation sites exclusive to the SOSi/FGF2-FBS treatment are highlighted in red. In total, 11 sites were exclusively identified across treatments and time points.

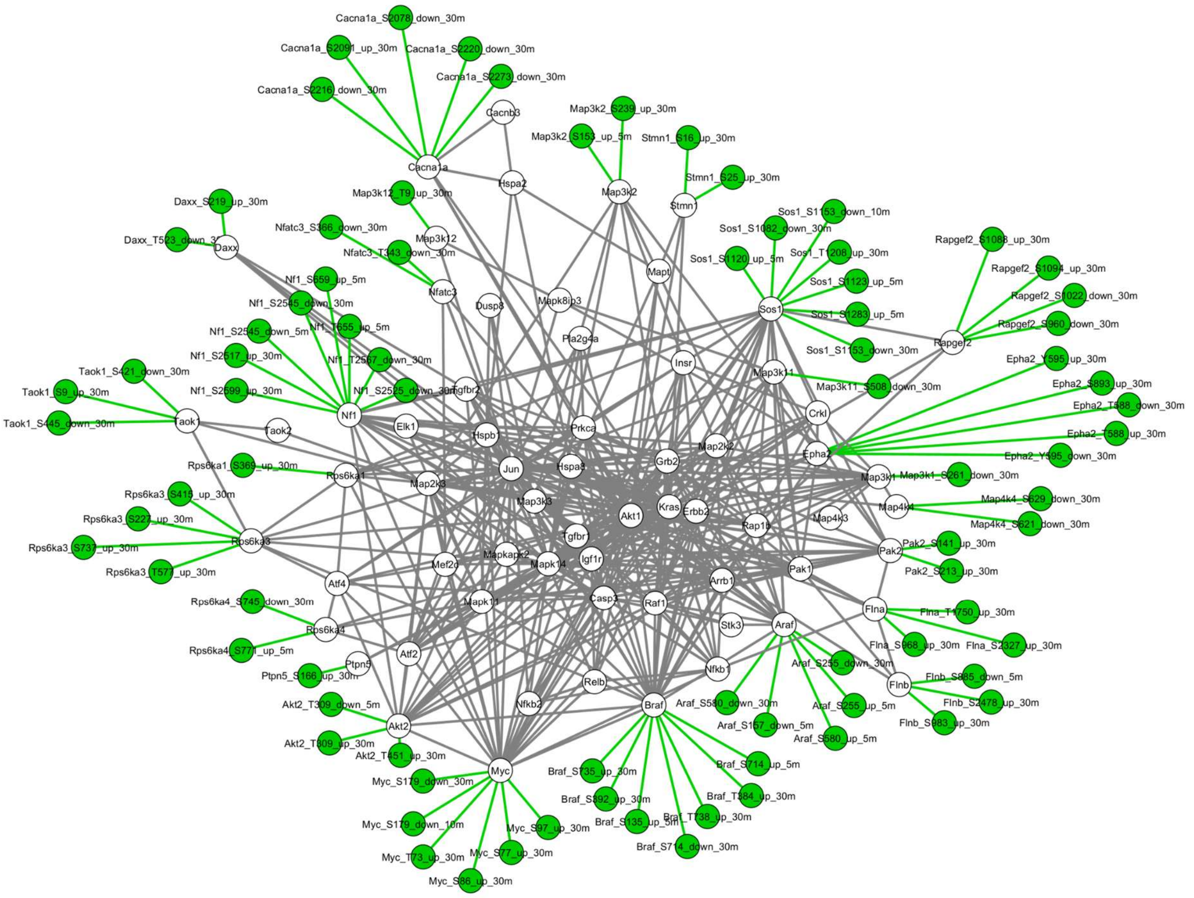


**Figure S10:** Top 30 proteins with the highest number of phosphorylation sites and p < 0.05 (Limma analysis) network was compared using DyNet for the FGF2+FBS versus SOSi/FGF2+FBS treatment. Phosphorylation sites exclusive to the FGF2+FBS treatment are highlighted in green. In total, 82 sites were differentially identified across treatments and time points.

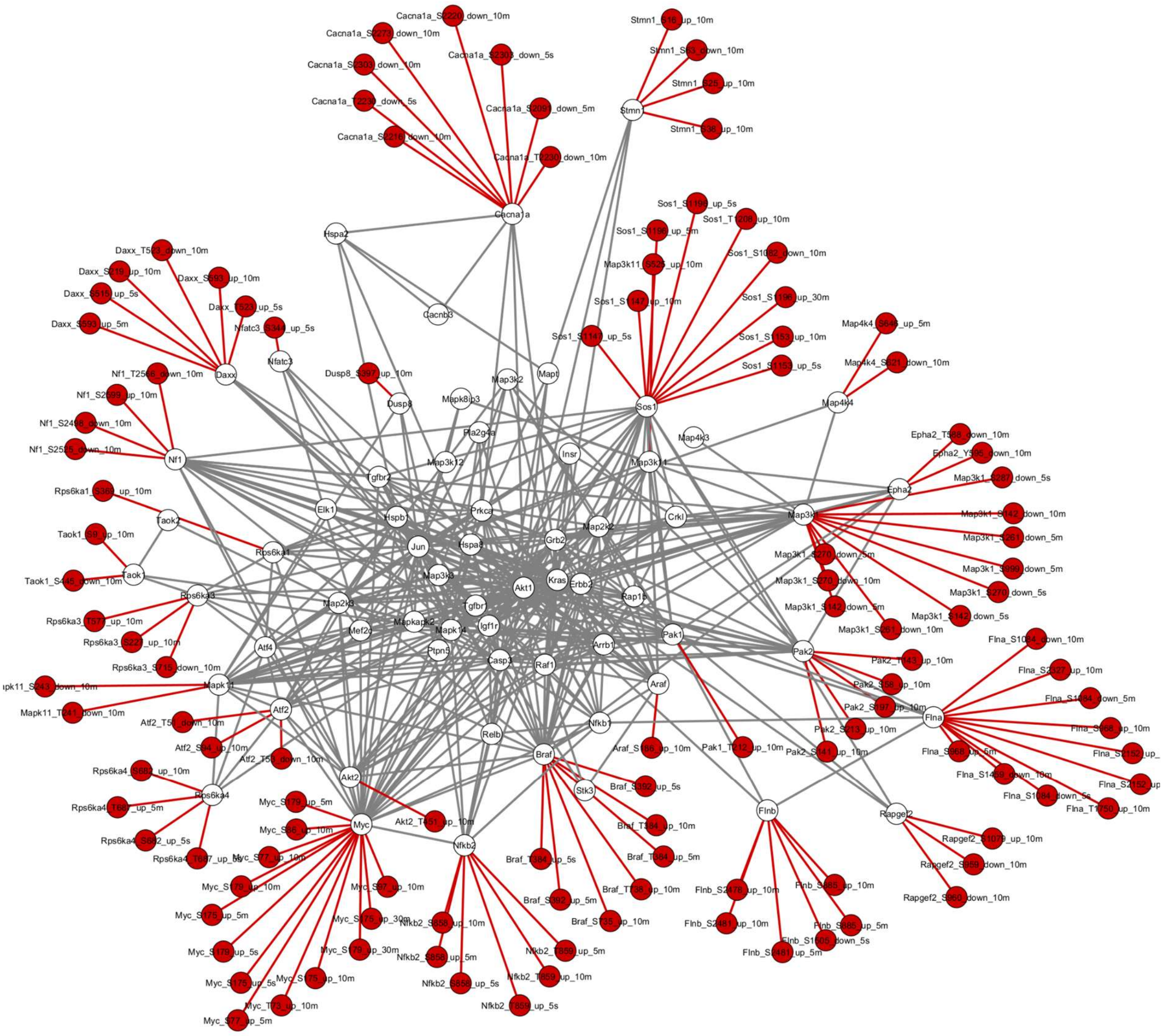


**Figure S11:** Top 30 proteins with the highest number of phosphorylation sites and $p < 0.05$ (Limma analysis) network was compared using DyNet for the SOSi/FGF2+FBS versus FGF2+FBS treatment. Phosphorylation sites exclusive to the SOSi/FGF2+FBS treatment are highlighted in red. In total, 116 sites were exclusively identified across treatments and time points.

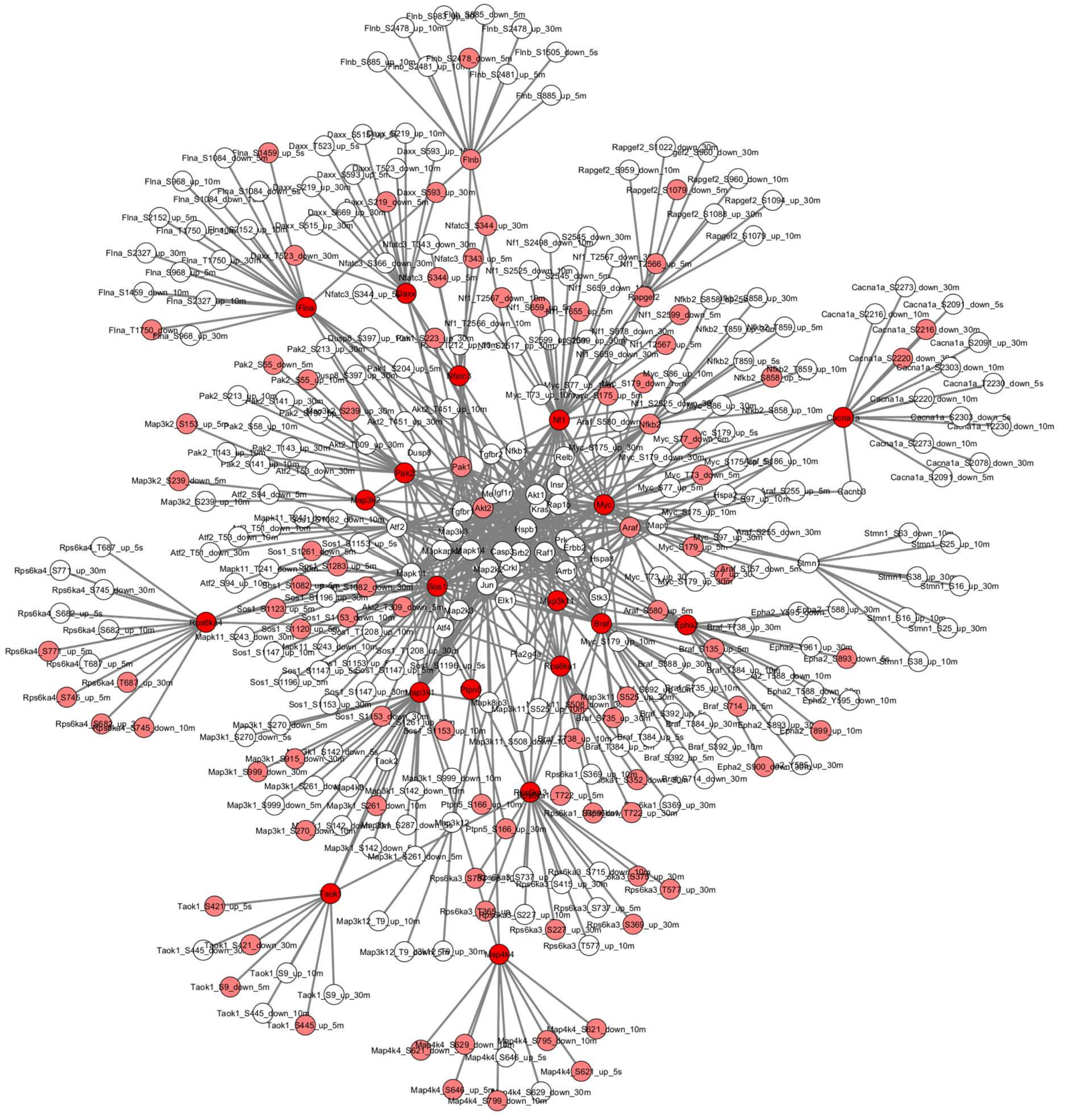

**Figure S12:** Sites with the most variations identified using the DyNet tool, which ranks sites that vary most between FGF2-FBS and SOSi/ FGF2-FBS conditions. The nodes in dark red represent the DyNet variation score higher than 0.5.

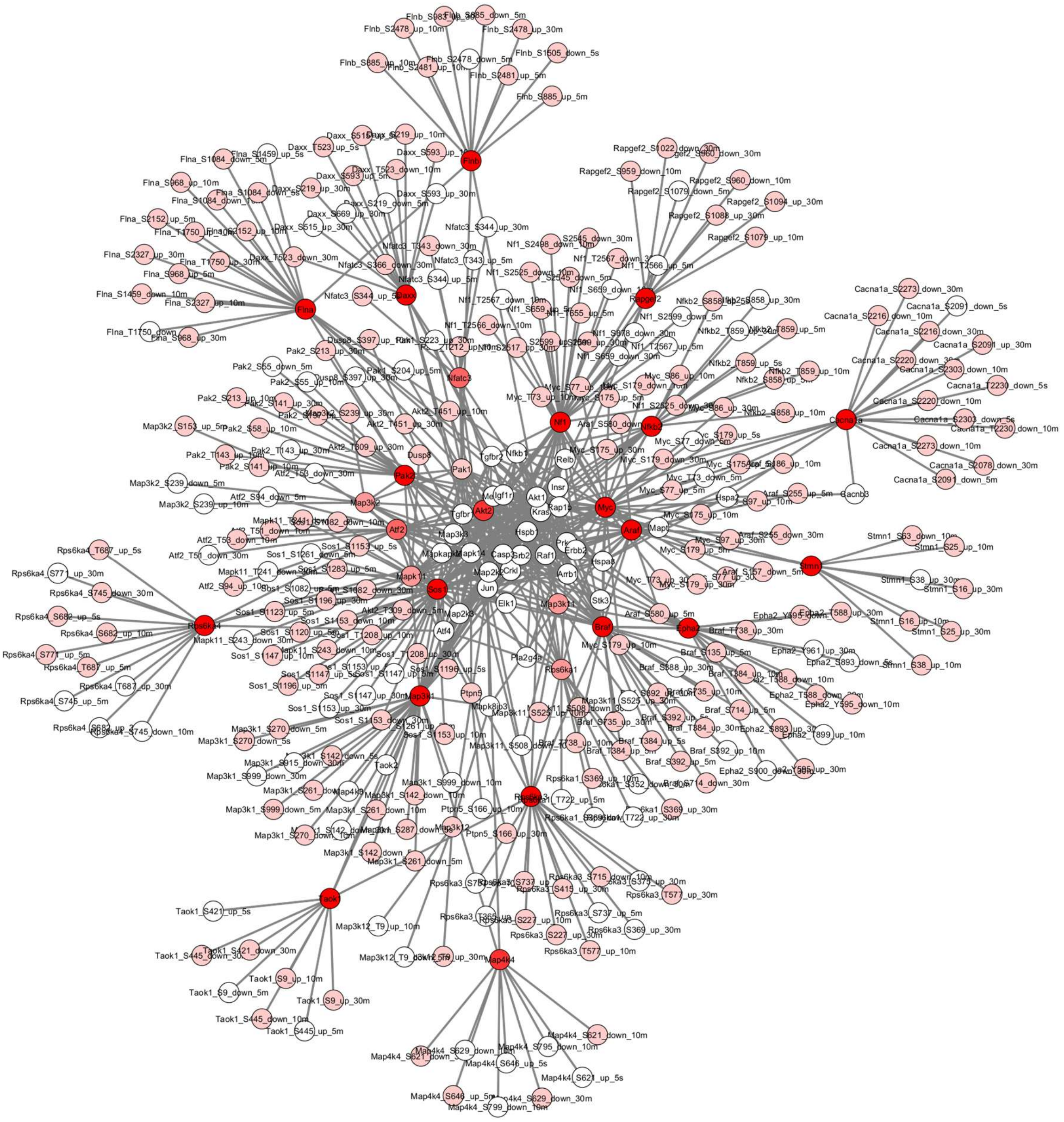


**Figure S13:** Sites with the most variations identified using the DyNet tool, which ranks sites that vary most between FGF2+FBS and SOSi/ FGF2+FBS conditions. The nodes in dark red represent the DyNet variation score higher than 0.5.

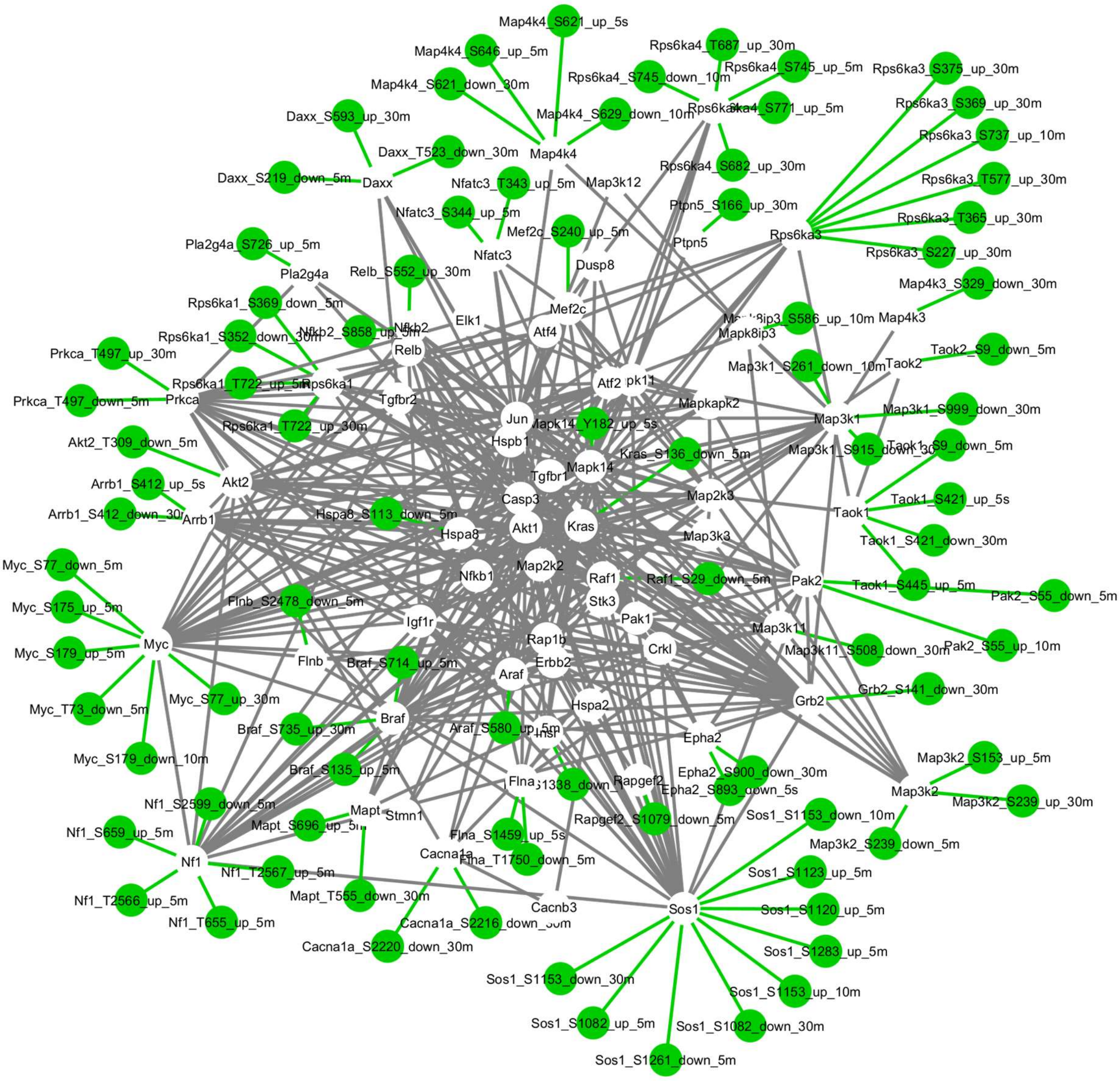


**Figure S14:** Among all genes from the KEGG MAPK pathway (mmu04010) with phosphorylation sites showing $p < 0.05$ (Limma analysis), the network was compared using DyNet for the FGF2-FBS versus SOSi/FGF2-FBS treatment. Phosphorylation sites exclusive to the FGF2-FBS treatment are highlighted in green. In total, 51 sites were exclusively identified across treatments and time points.

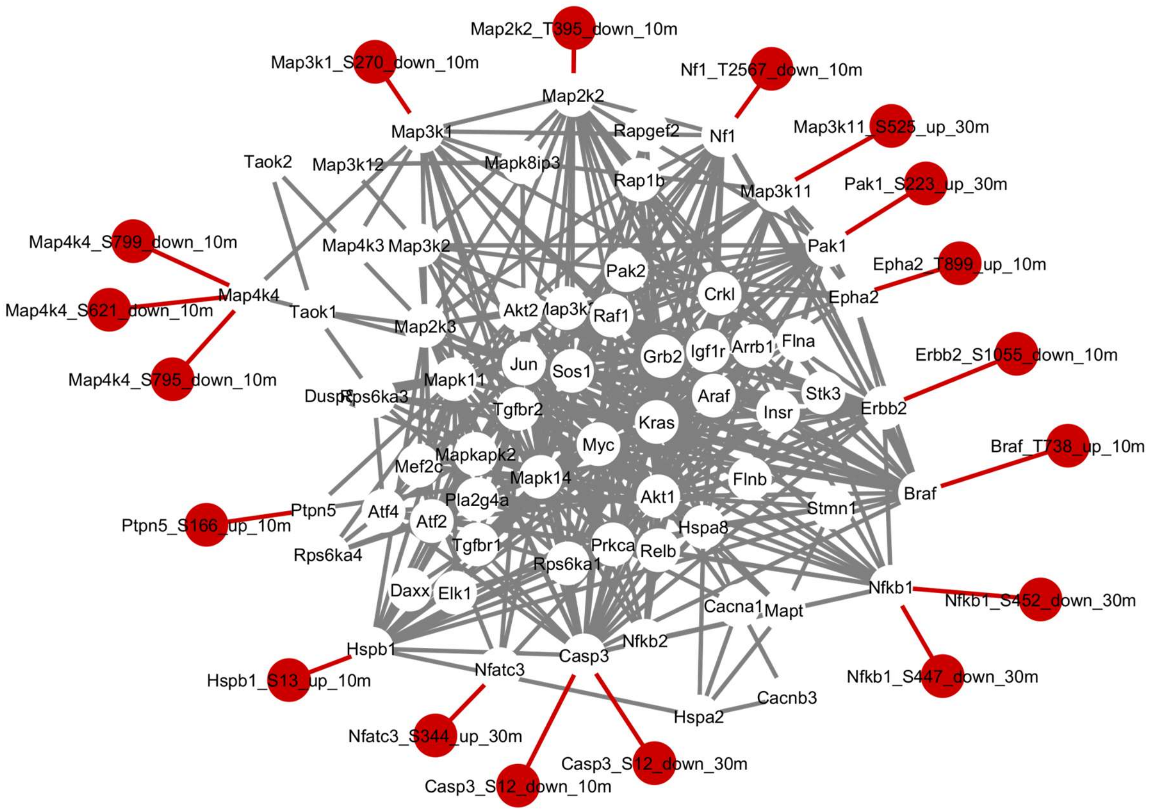


**Figure S15:** Among all genes from the KEGG MAPK pathway (mmu04010) with phosphorylation sites showing p < 0.05 (Limma analysis), the network was compared using DyNet for the SOSi/FGF2-FBS versus FGF2-FBS treatment. Phosphorylation sites exclusive to the SOSi/FGF2-FBS treatment are highlighted in red. In total, 18 sites were exclusively identified across treatments and time points.

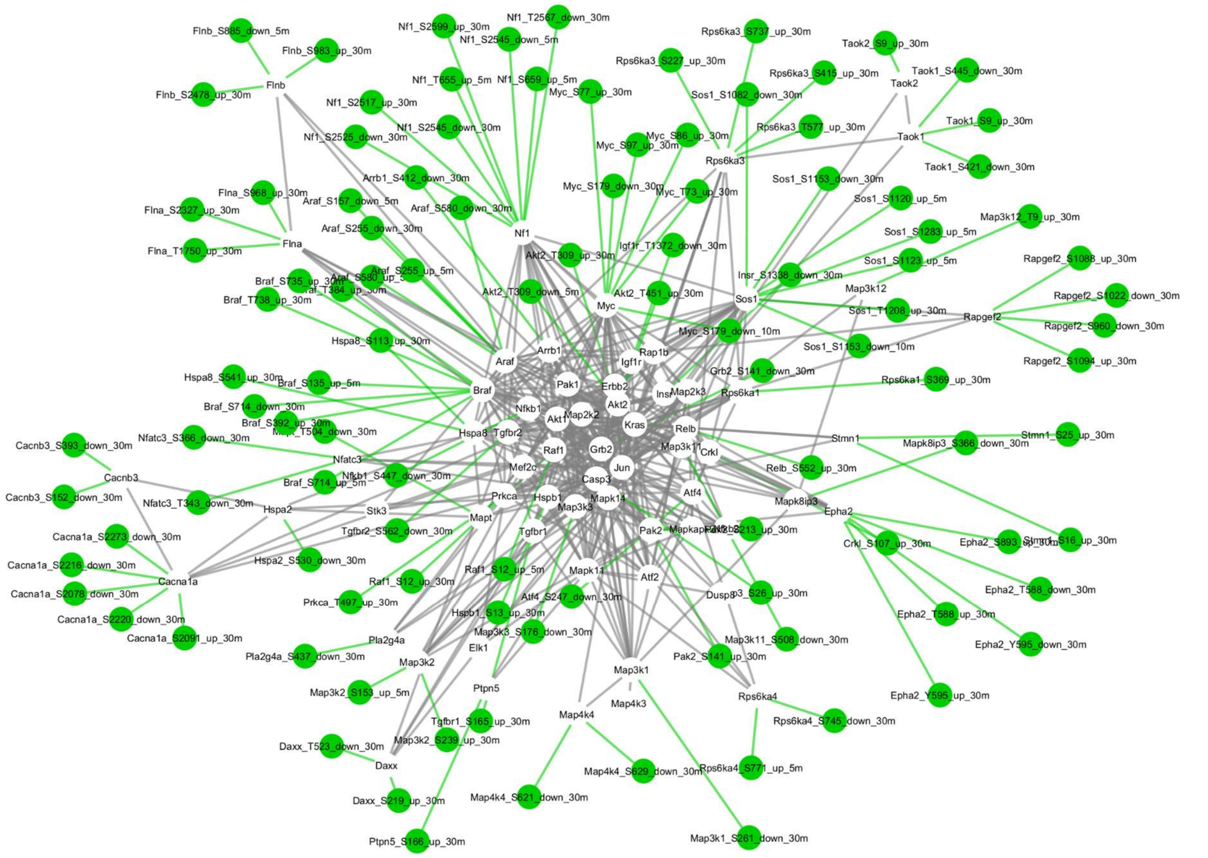


**Figure S16:** Among all genes from the KEGG MAPK pathway (mmu04010) with phosphorylation sites showing p < 0.05 (Limma analysis), the network was compared using DyNet for the FGF2+FBS versus SOSi/FGF2+FBS treatment. Phosphorylation sites exclusive to the FGF2+FBS treatment are highlighted in green. In total, 107 sites were exclusively identified across treatments and time points.

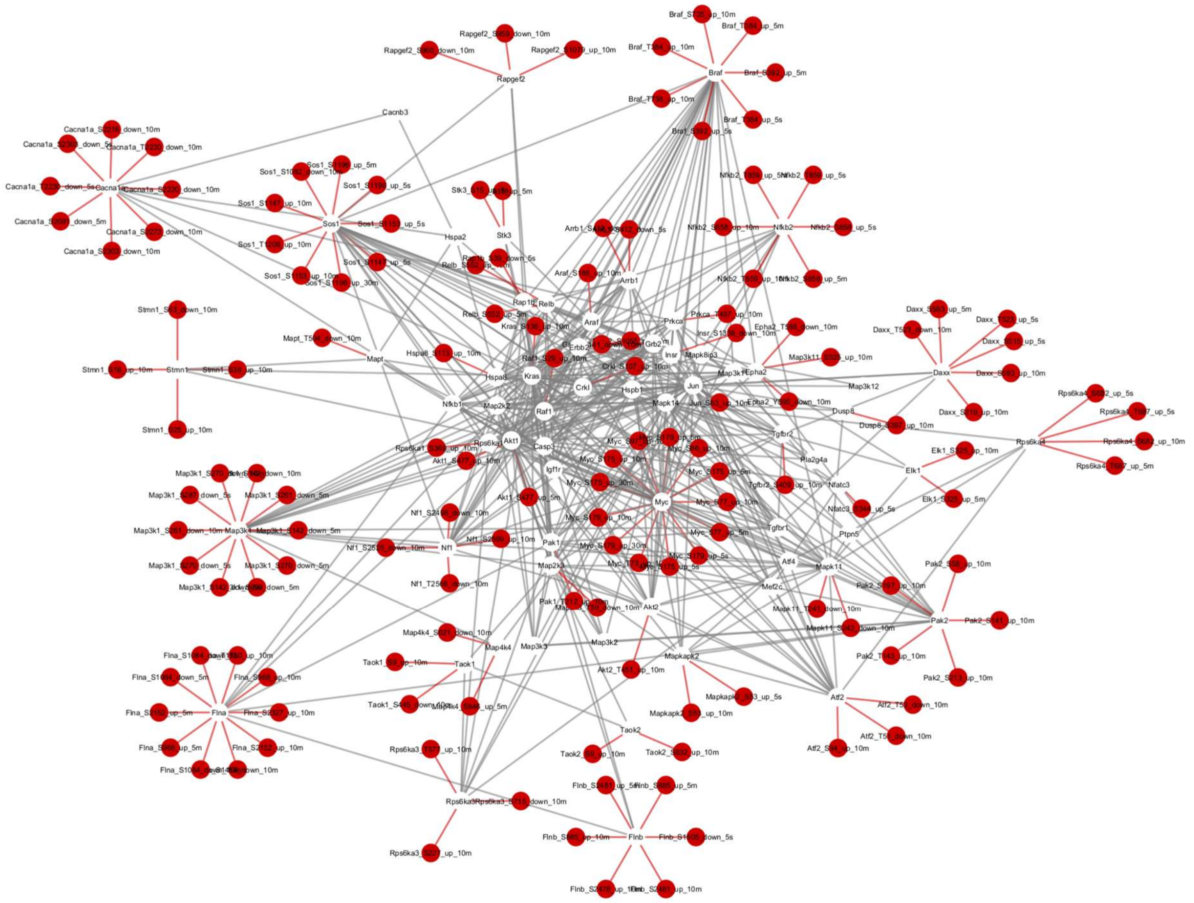


**Figure S17:** Among all genes from the KEGG MAPK pathway (mmu04010) with phosphorylation sites showing p < 0.05 (Limma analysis), the network was compared using DyNet for the SOSi/FGF2+FBS versus FGF2+FBS treatment. Phosphorylation sites exclusive to the SOSi/FGF2+FBS treatment are highlighted in red. In total, 143 sites were exclusively identified across treatments and time points.

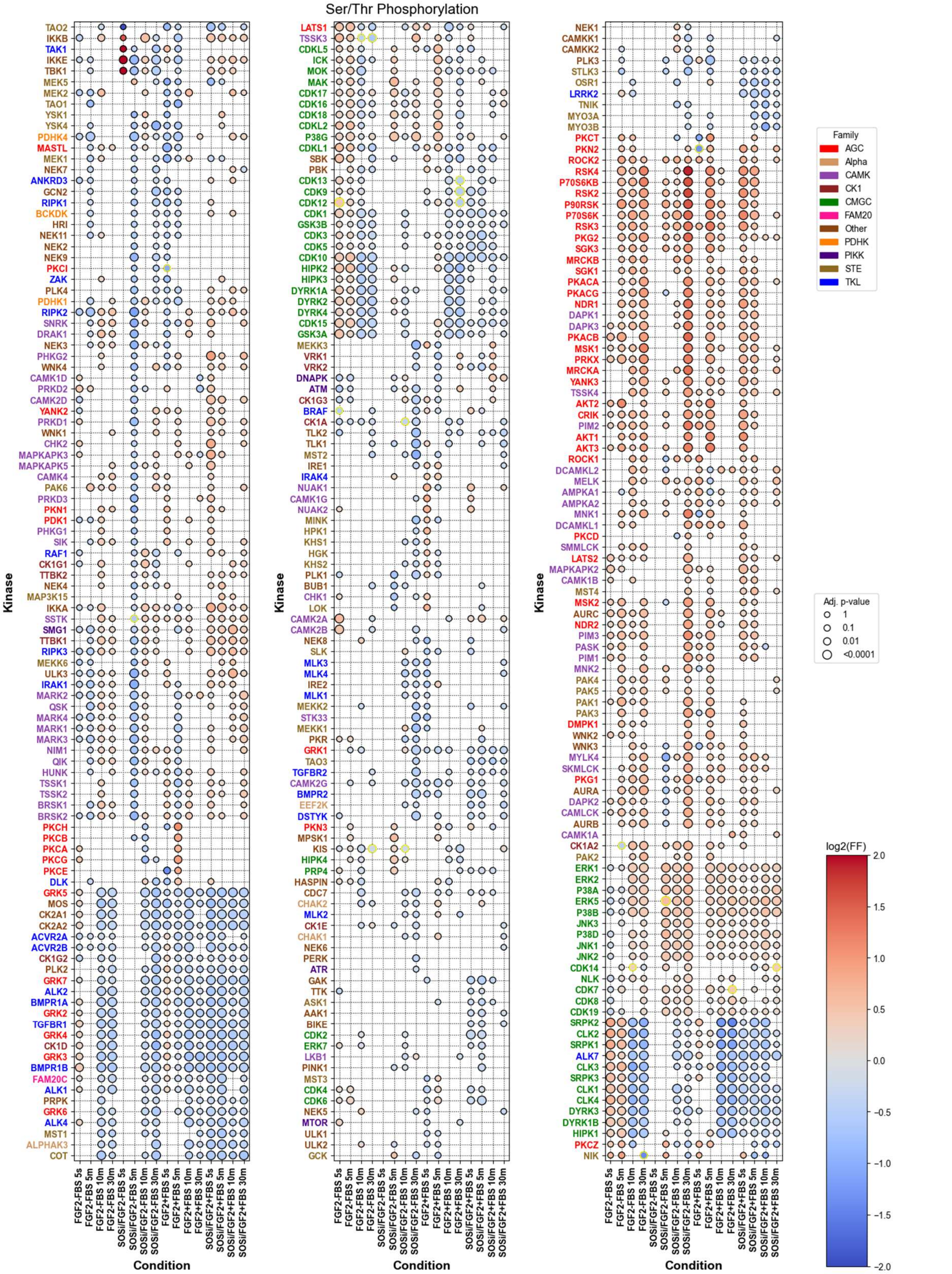


**Figure S18**: Differentially phosphorylated serine/threonine sites for each limma contrast were identified using the DiffPhosData class from the kinase_library Python package. Sites with adjusted p-value < 0.05 were analyzed using log fold-change values and phosphosite sequence windows. Kinase enrichment was inferred using percentile-rank scoring (threshold = 15). **Legend: Red:** Upregulated phosphorylation events. **Blue:** blue indicates downregulated events. The individual graphics and the source table for this result can be seen on **Figures S19-26** and **Supplemental Table S6**.

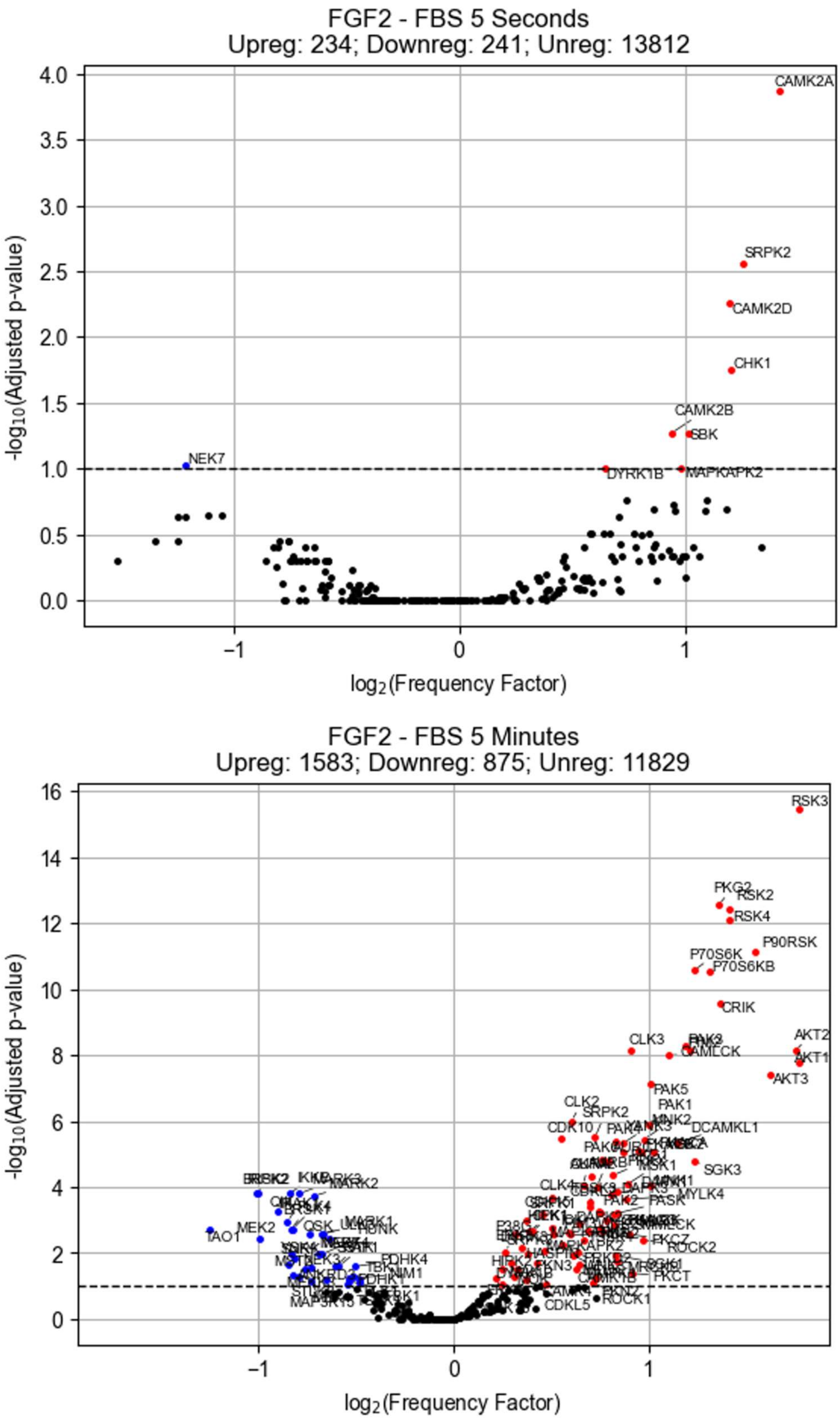


**Figure S19**: Serine/threonine kinase activity (FGF2 minus FBS 5 seconds and 5 minutes). Kinase library, FDR 0.05%, kin_type = ’ser_thr’, method = ’percentile_rank’, thresh = 15.

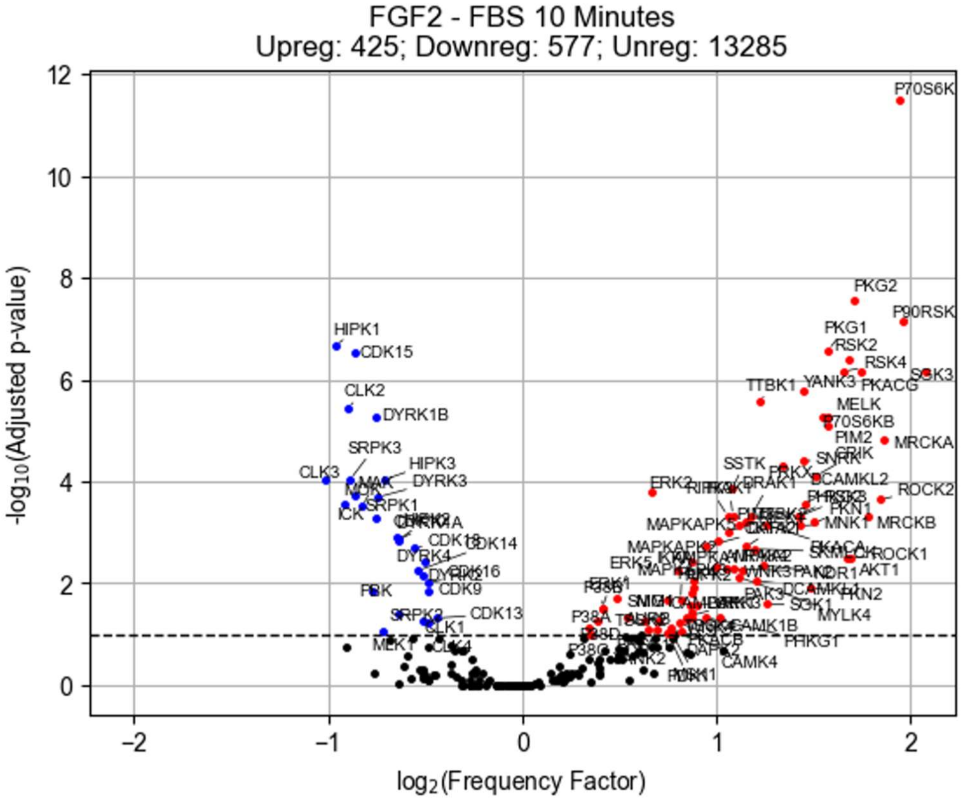


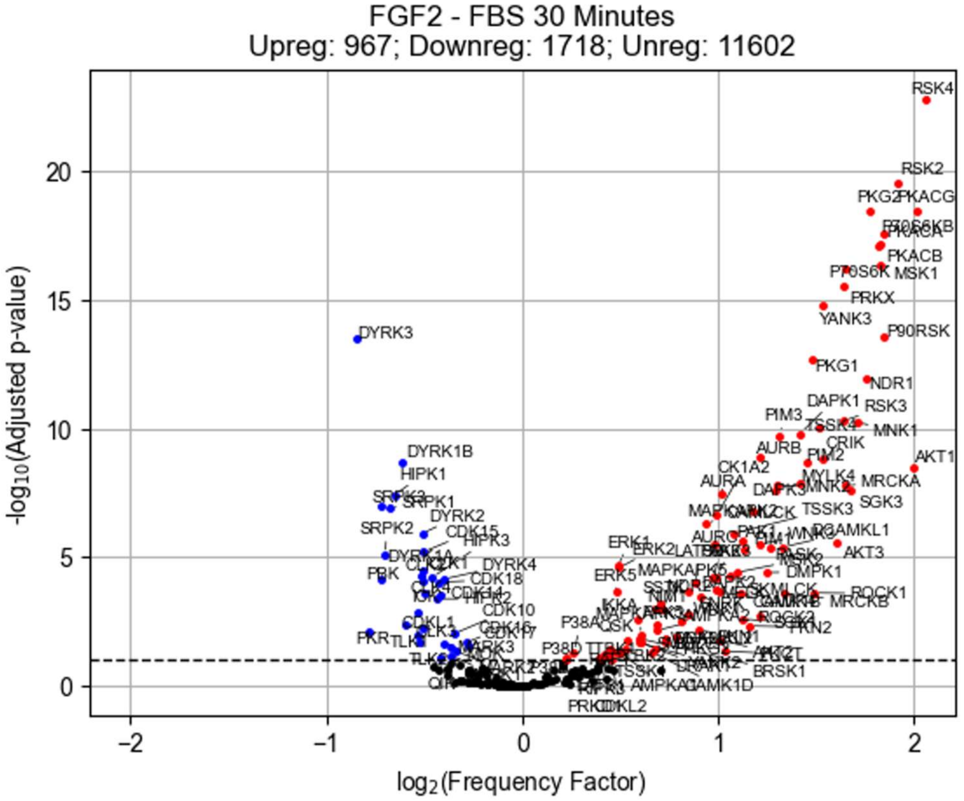


**Figure S20**: Serine/threonine kinase activity (FGF2 minus FBS 10 minutes and 30 minutes). Kinase library, FDR 0.05%, kin_type = ’ser_thr’, method = ’percentile_rank’, thresh = 15

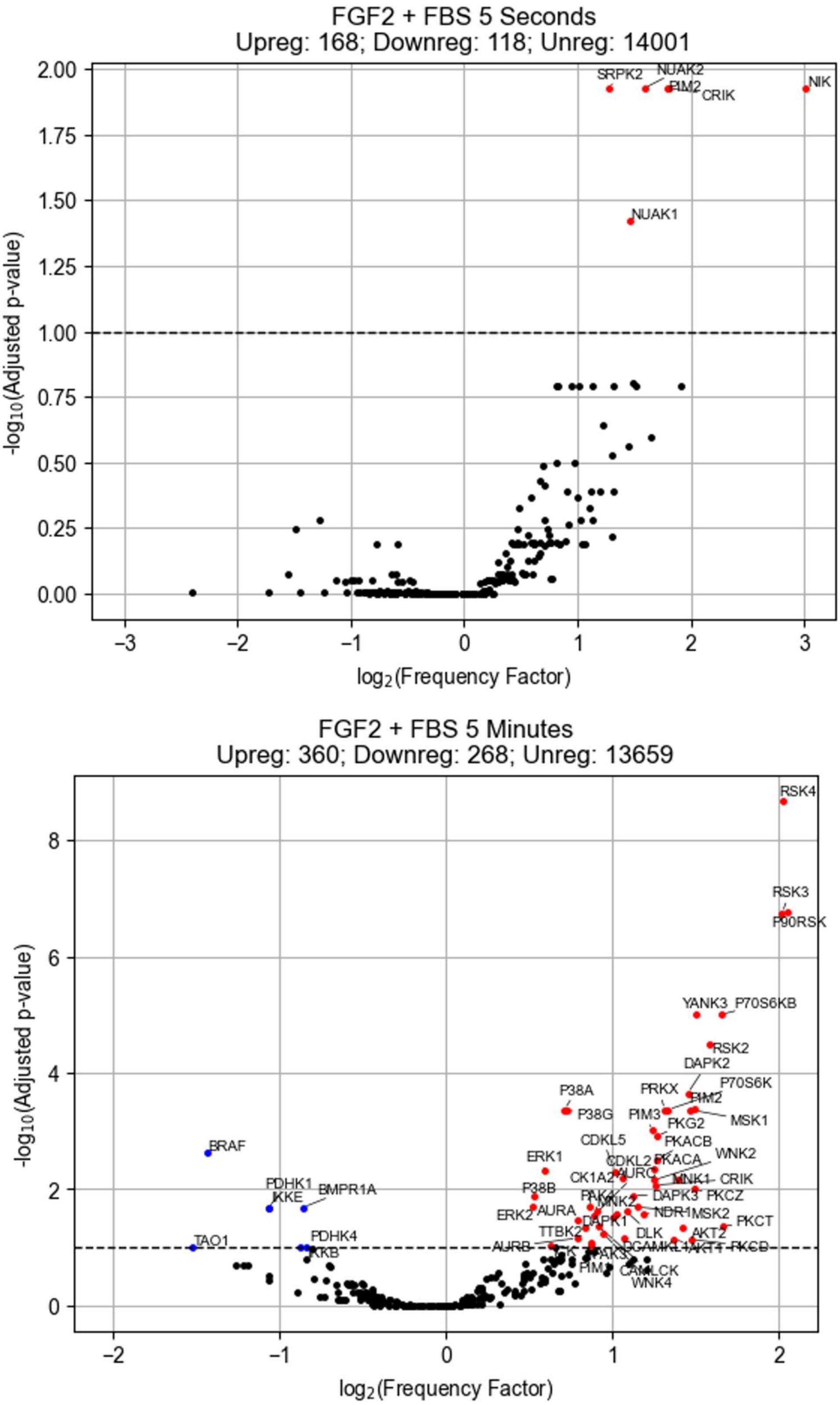


**Figure S21:** Serine/threonine kinase activity (FGF2 plus FBS 5 seconds and 5 minutes). Kinase library, FDR 0.05%, kin_type = 'ser_thr', method = 'percentile_rank', thresh = 15.

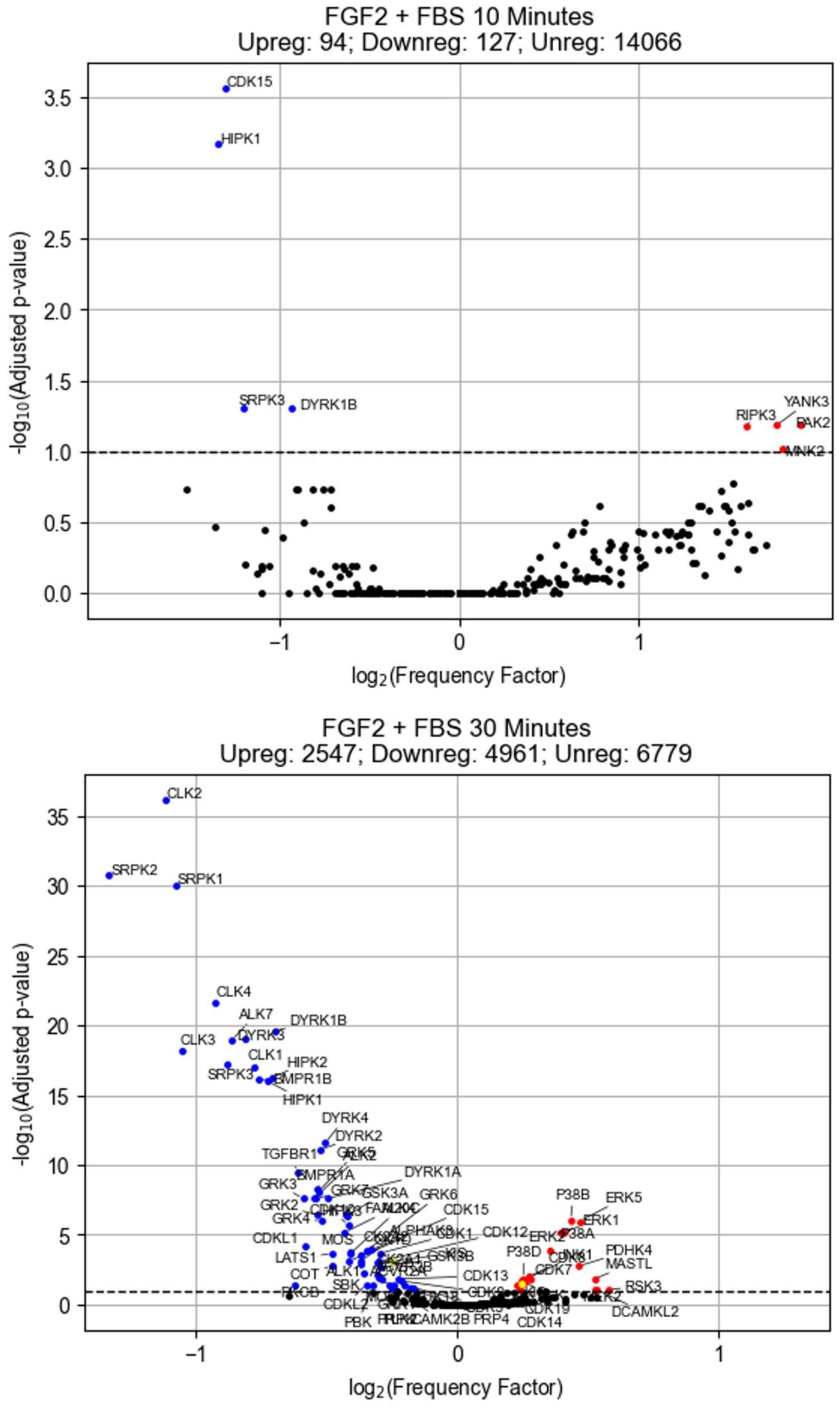


**Figure S22:** Serine/threonine kinase activity (FGF2 plus FBS 10 minutes and 30 minutes). Kinase library, FDR 0.05%, kin_type = 'ser_thr', method = 'percentile_rank', thresh = 15.

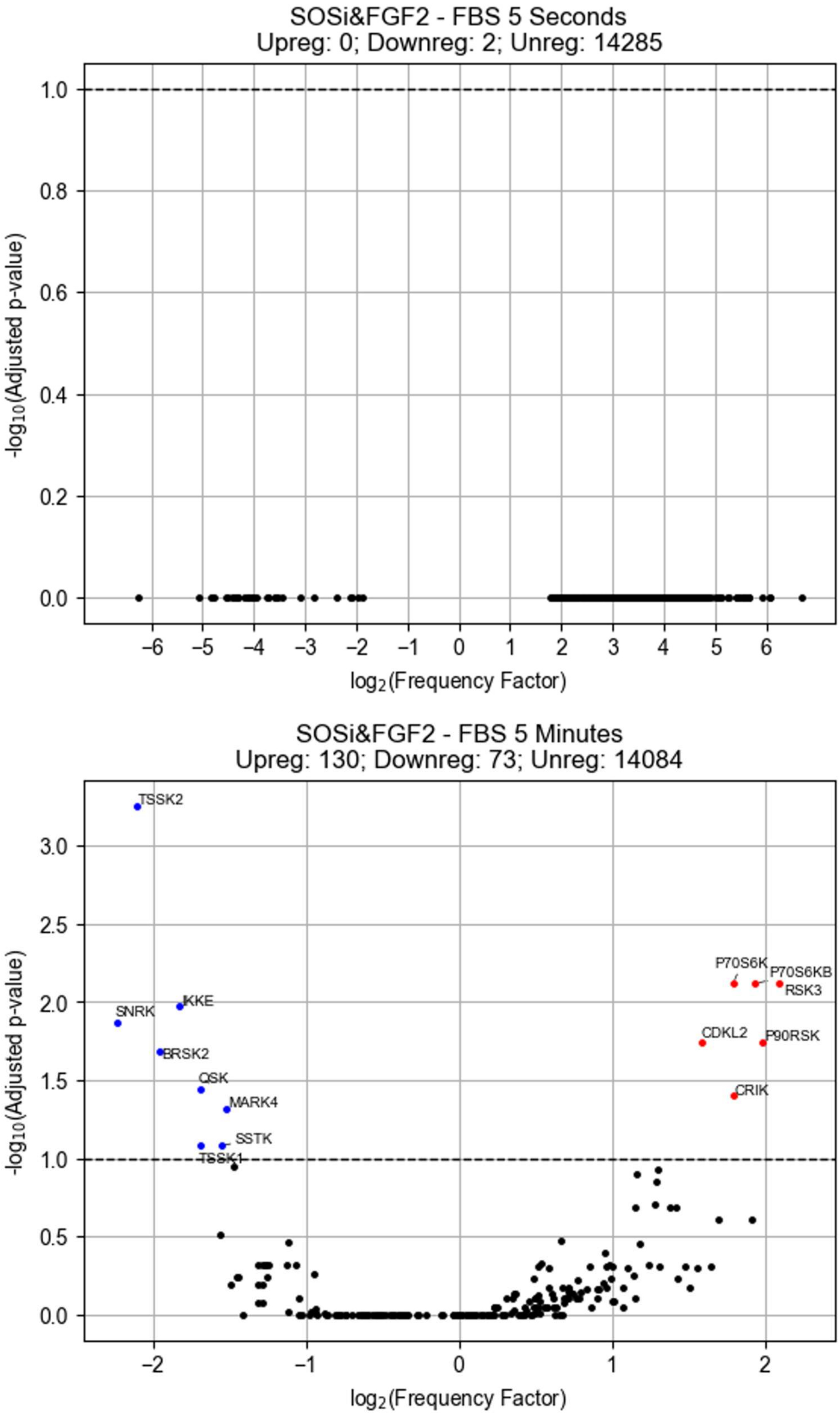


**Figure S23:** Serine/threonine kinase activity (SOSi/FGF2 5 minus FBS seconds and 5 minutes). Kinase library, FDR 0.05%, kin_type = 'ser_thr', method = 'percentile_rank', thresh = 15.

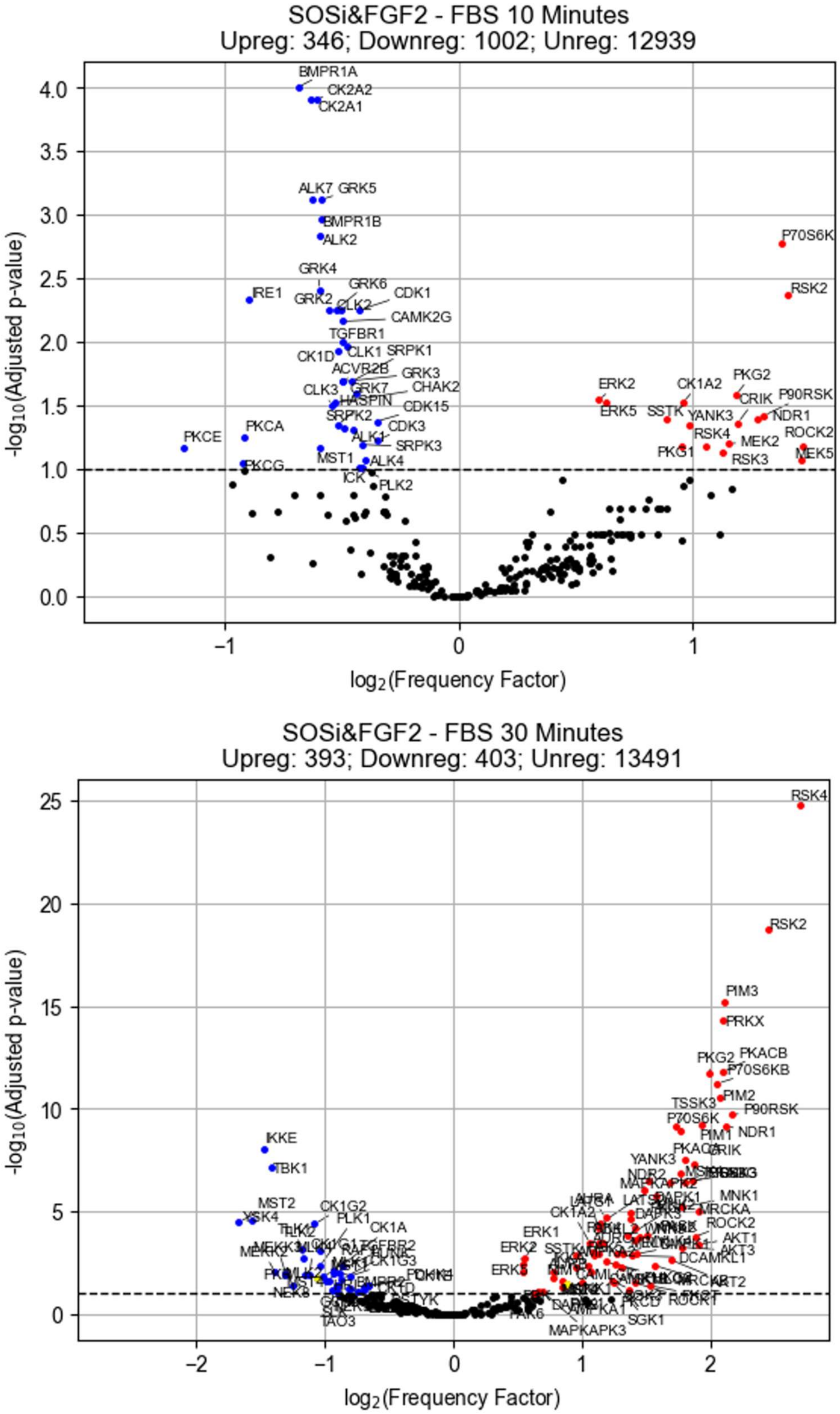


**Figure S24:** Serine/threonine kinase activity (SOSi/FGF2 minus FBS 10 minutes and 30 minutes). Kinase library, FDR 0.05%, kin_type = ’ser_thr’, method = ’percentile_rank’, thresh = 15.

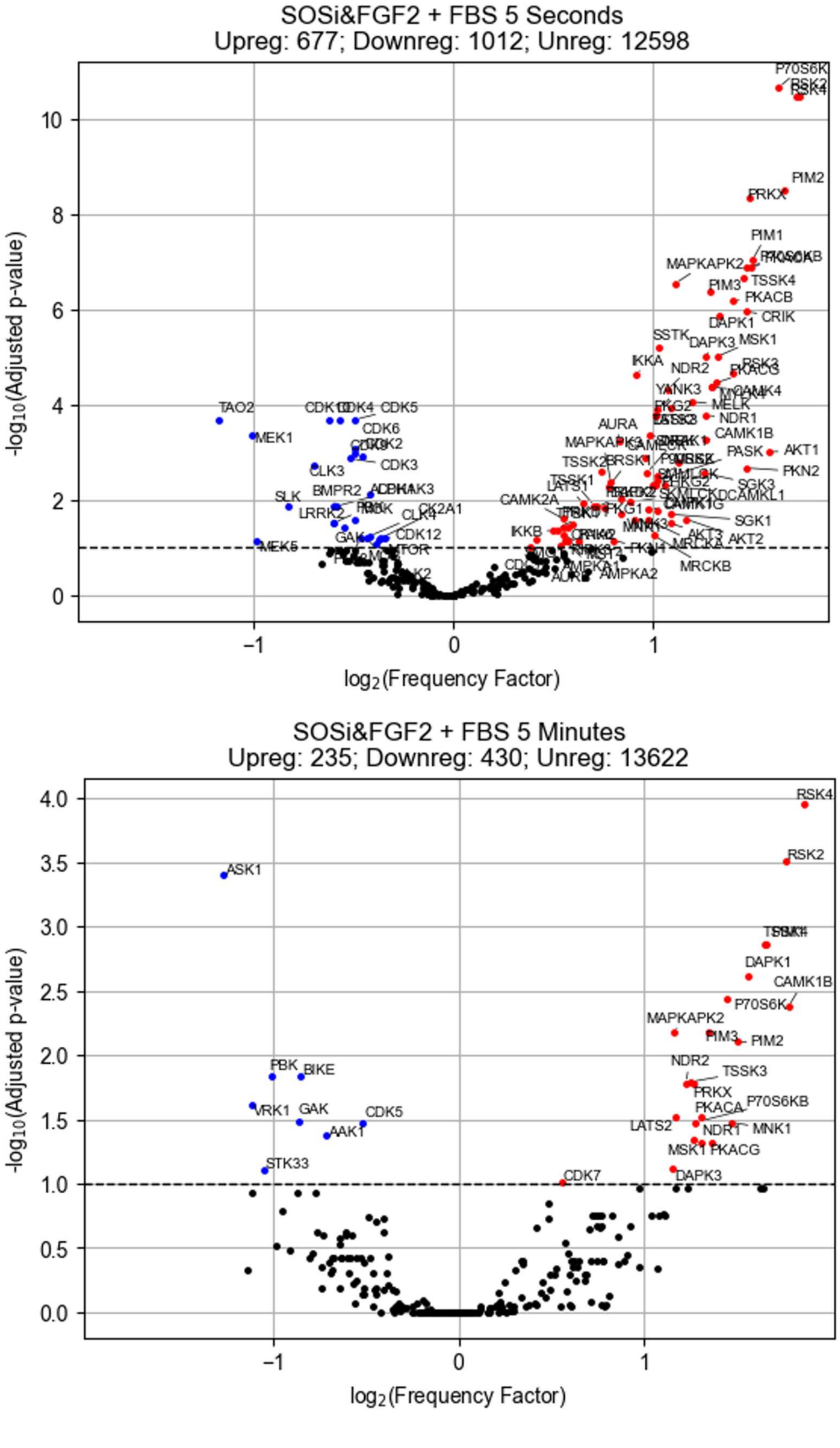


**Figure S25:** Serine/threonine kinase activity (SOSi/FGF2 plus FBS 5 seconds and 5 minutes). Kinase library, FDR 0.05%, kin_type = ’ser_thr’, method = ’percentile_rank’, thresh = 15.

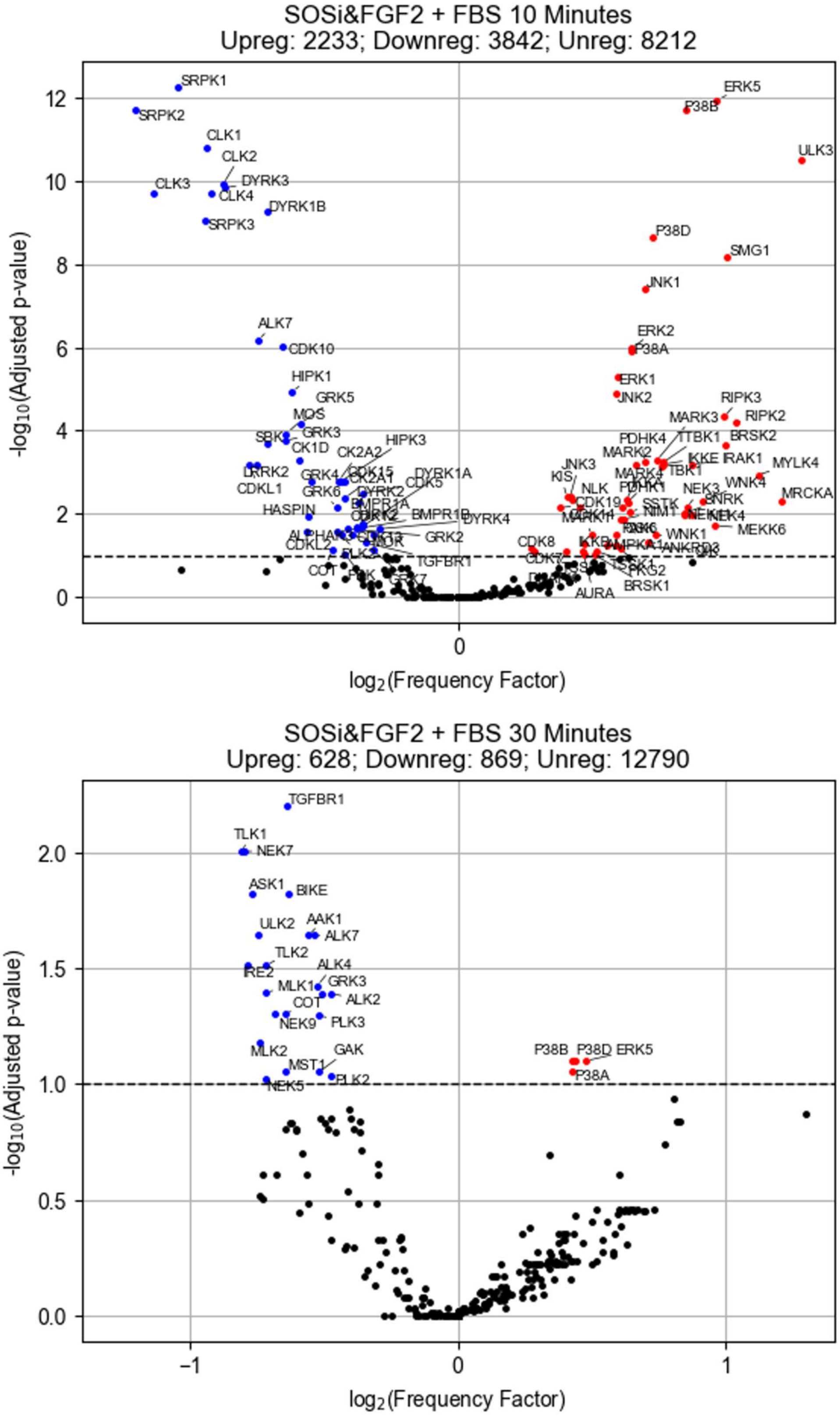


**Figure S26:** Serine/threonine kinase activity (SOSi/FGF2 plus FBS 10 minutes and 30 minutes). Kinase library, FDR 0.05%, kin_type = ’ser_thr’, method = ’percentile_rank’, thresh = 15.

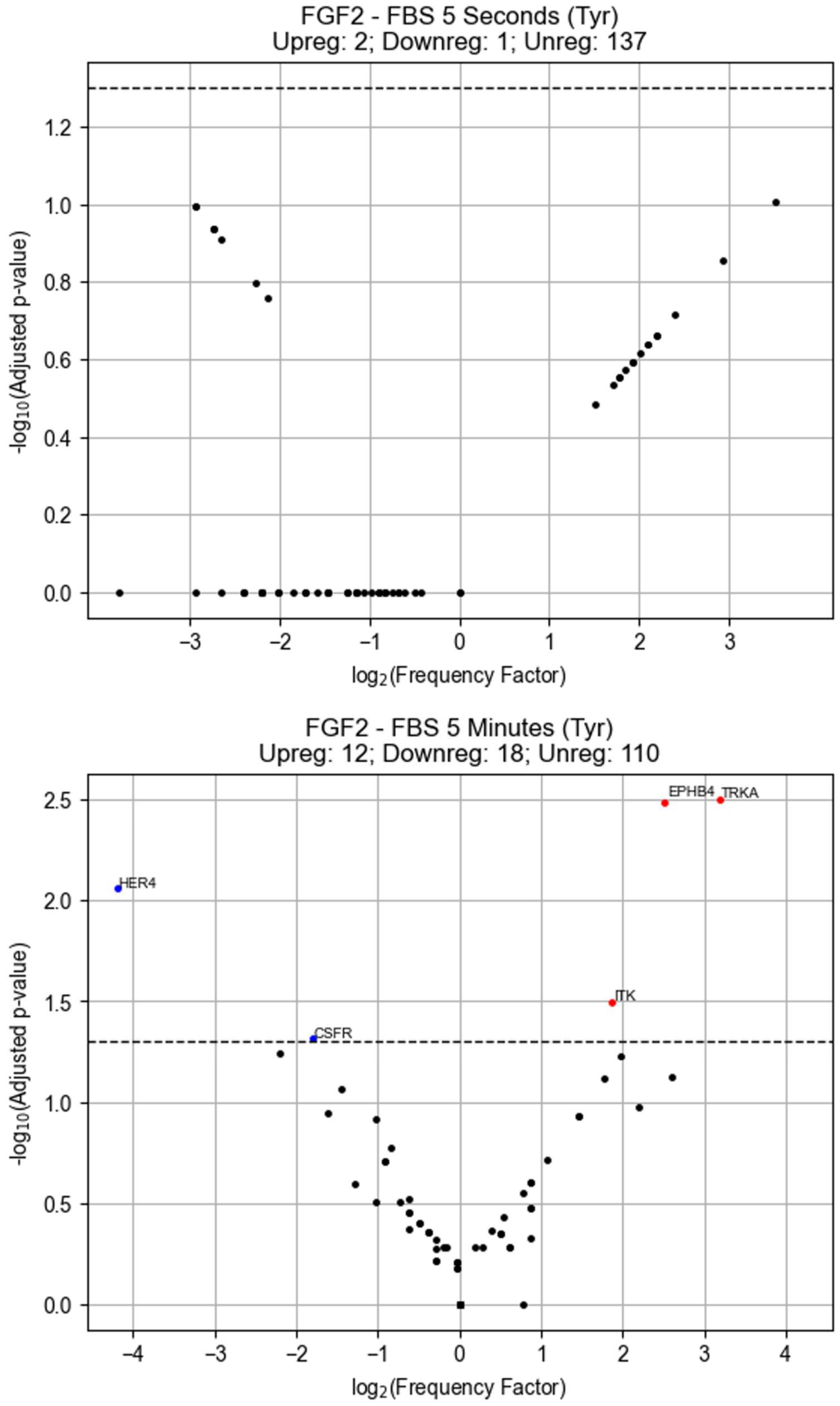


**Figure S27:** Tyrosine kinase activity (FGF2 minus FBS 5 seconds and 5 minutes). Kinase library, FDR 0.05%, kin_type = ’tyr’, method = ’percentile_rank’, thresh = 8.

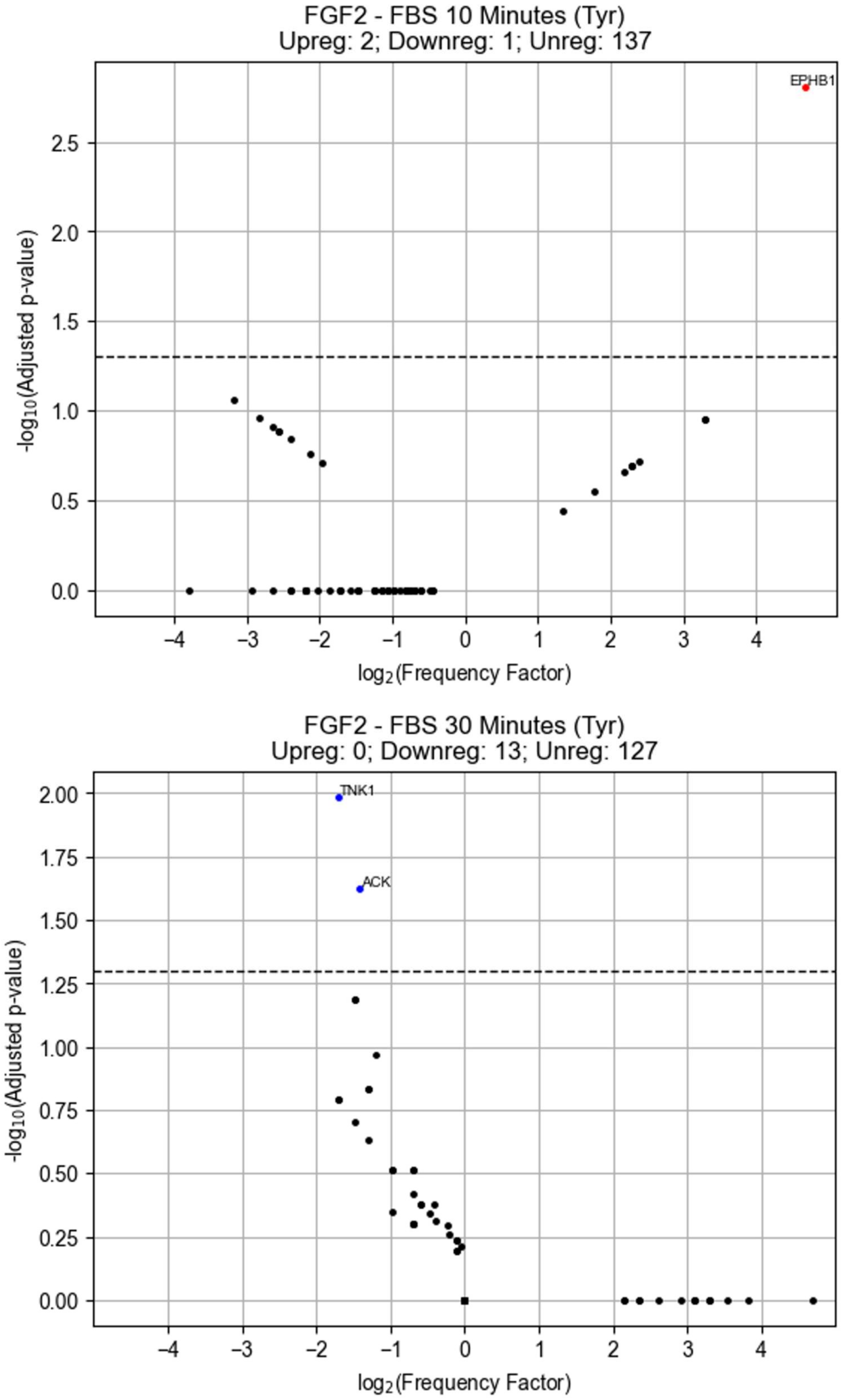


**Figure S28:** Tyrosine kinase activity (FGF2 minus FBS 10 minutes and 30 minutes). Kinase library, FDR 0.05%, kin_type = ’tyr’, method = ’percentile_rank’, thresh =8.

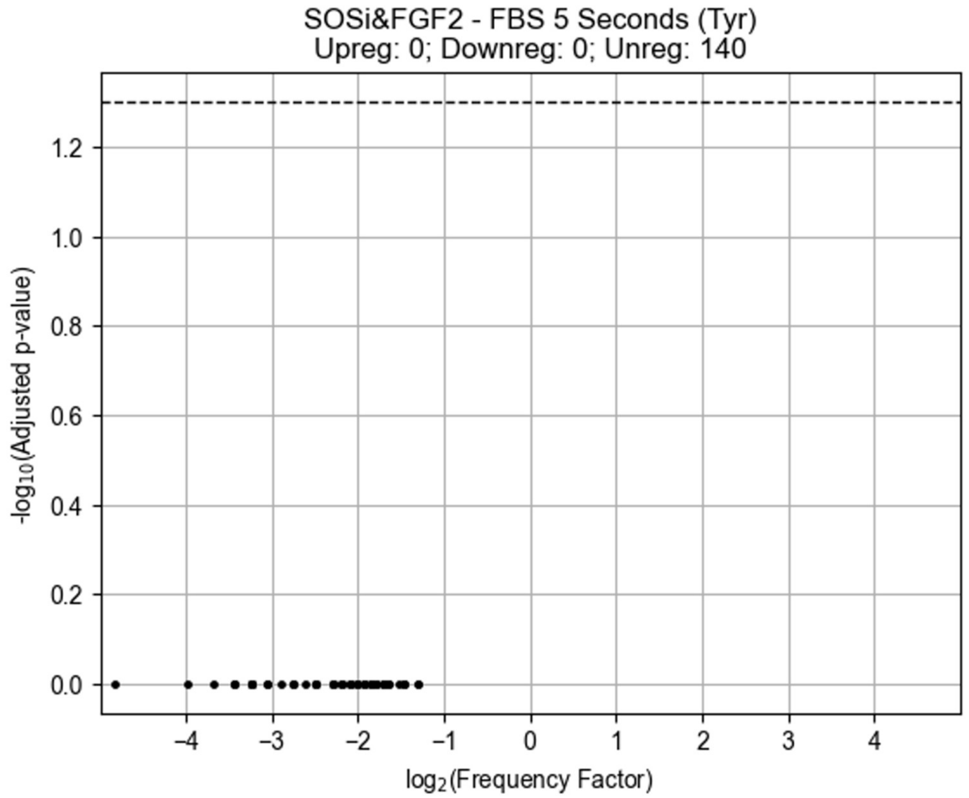


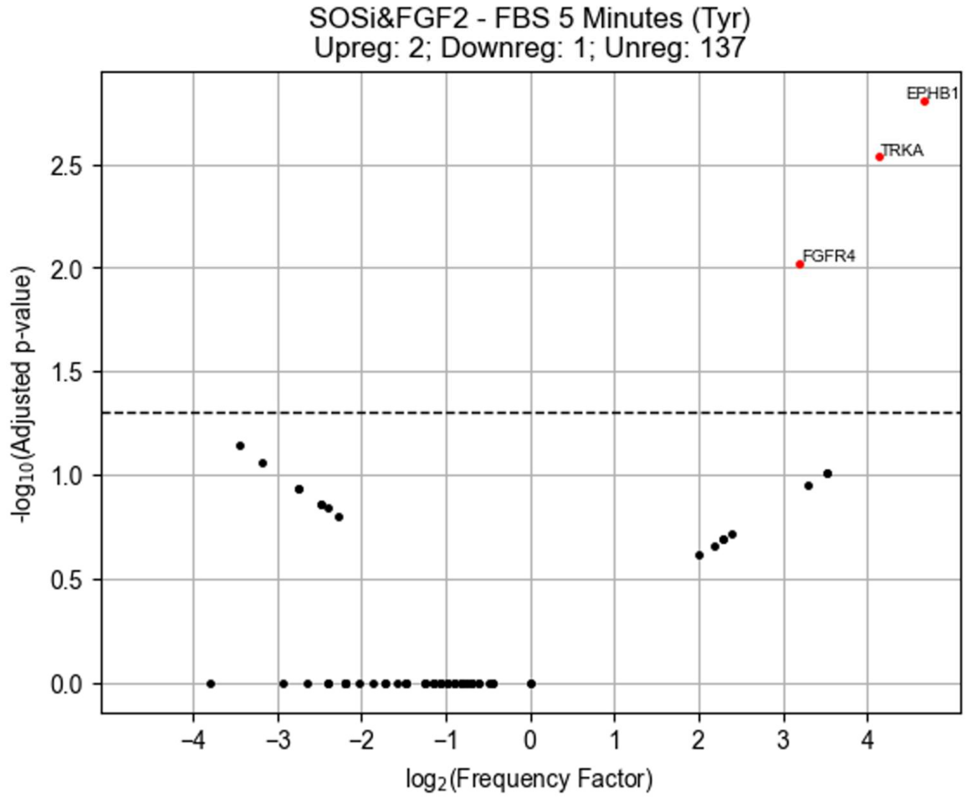


**Figure S29:** Tyrosine kinase activity (SOSi/FGF2 minus FBS 5 seconds and 5 minutes). Kinase library, FDR 0.05%, kin_type = ’tyr’, method = ’percentile_rank’, thresh = 8.

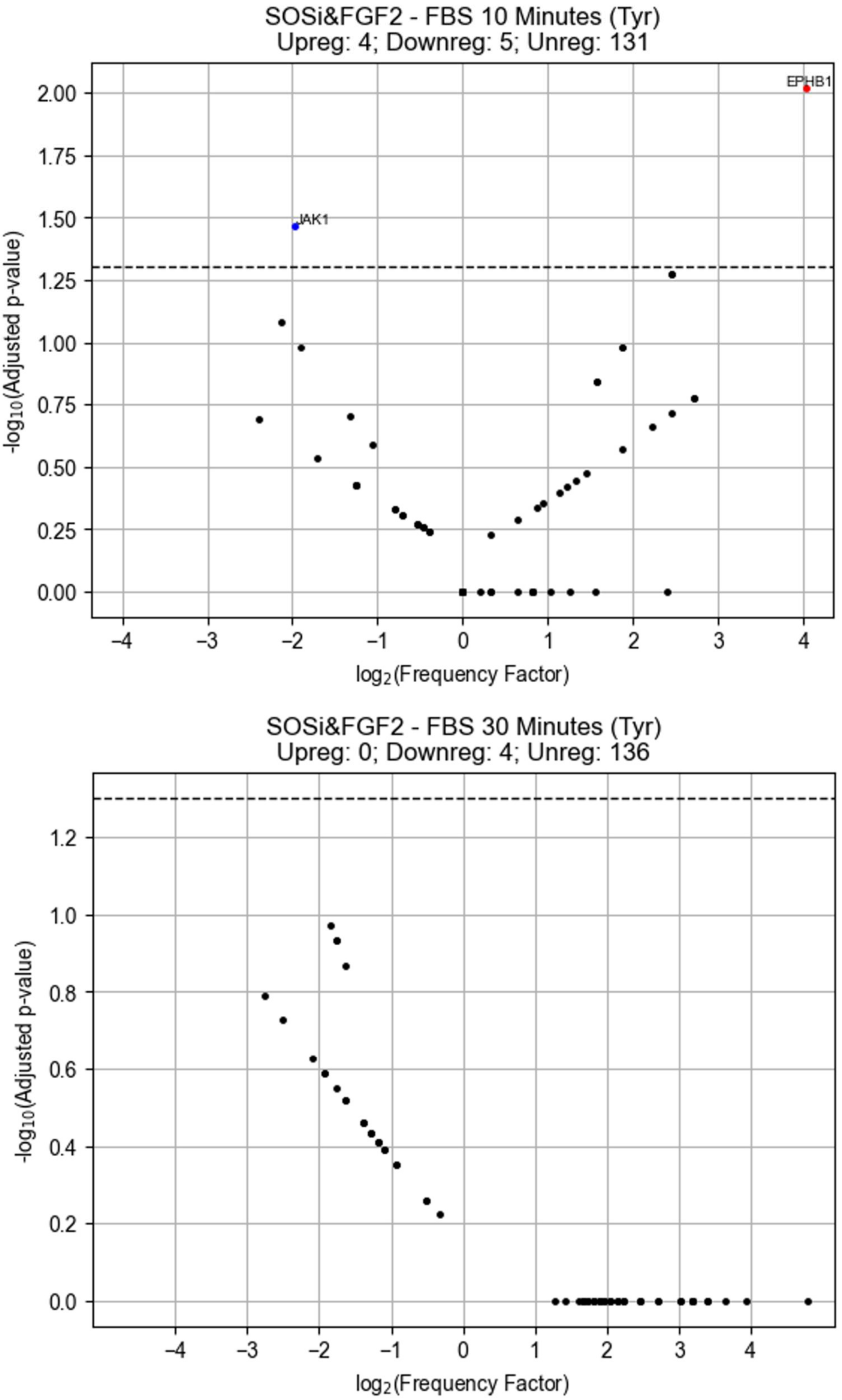


**Figure S30:** Tyrosine kinase activity (SOSi/FGF2 minus FBS 10 minutes and 30 minutes). Kinase library, FDR 0.05%, kin_type = 'tyr', method = 'percentile_rank', thresh = 8.

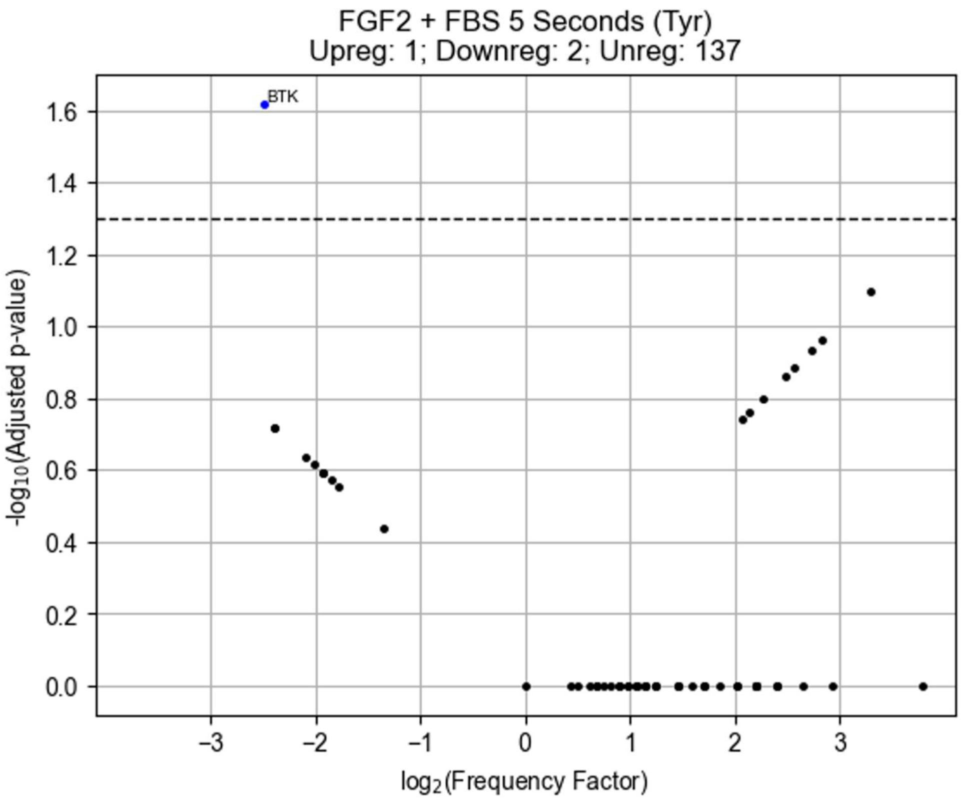


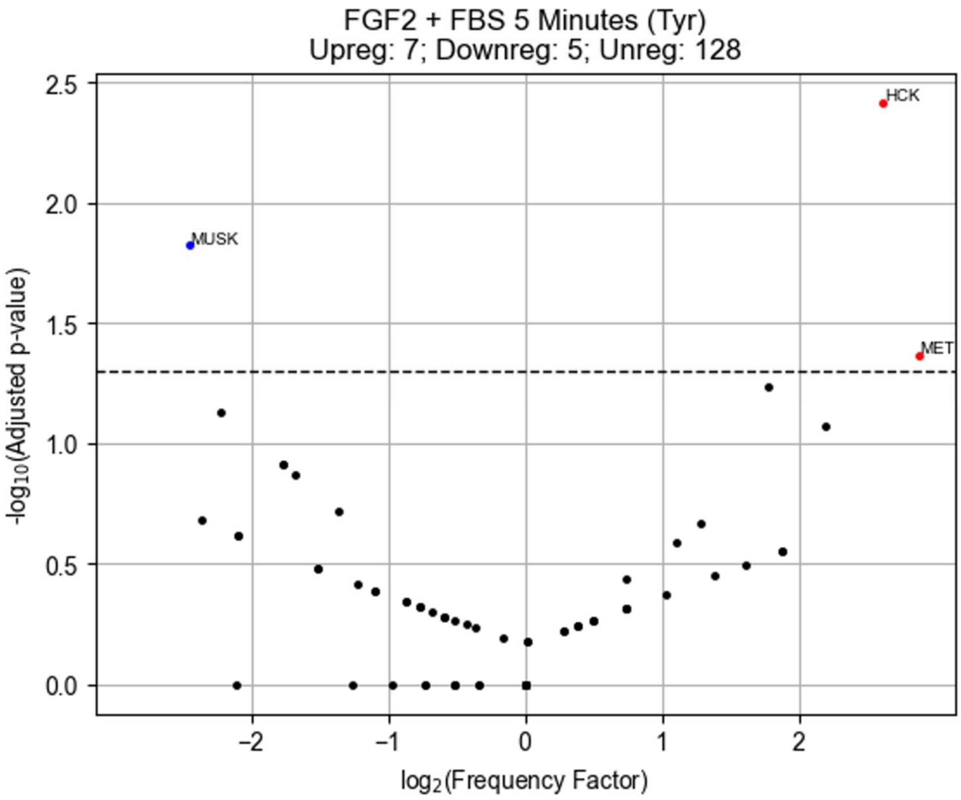


**Figure S31:** Tyrosine kinase activity (FGF2 plus FBS 5 seconds and 5 minutes). Kinase library, FDR 0.05%, kin_type = 'tyr', method = 'percentile_rank', thresh = 8.

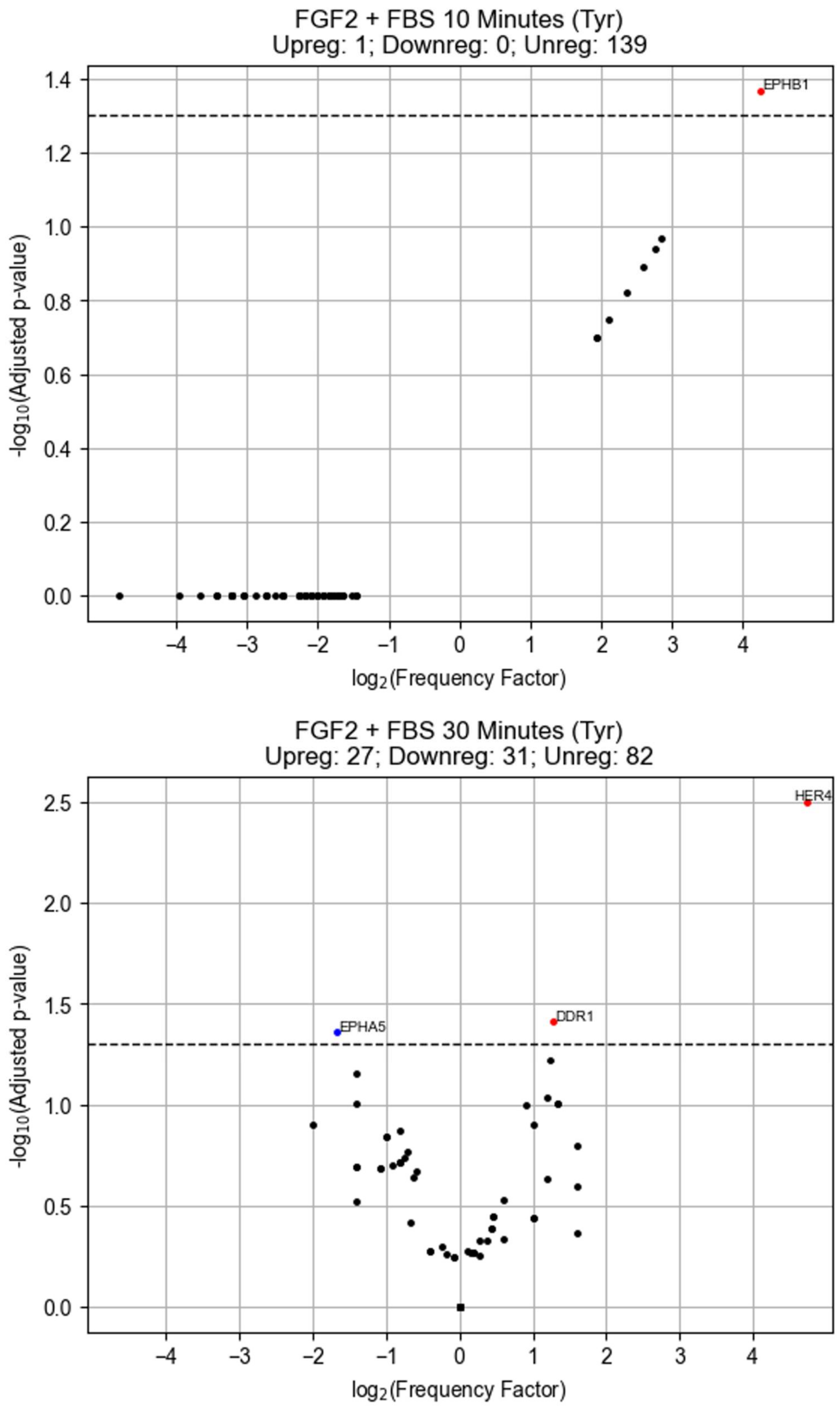


**Figure S32:** Tyrosine kinase activity (FGF2 plus FBS 10 minutes and 30 minutes). Kinase library, FDR 0.05%, kin_type = 'tyr', method = 'percentile_rank', thresh = 8.

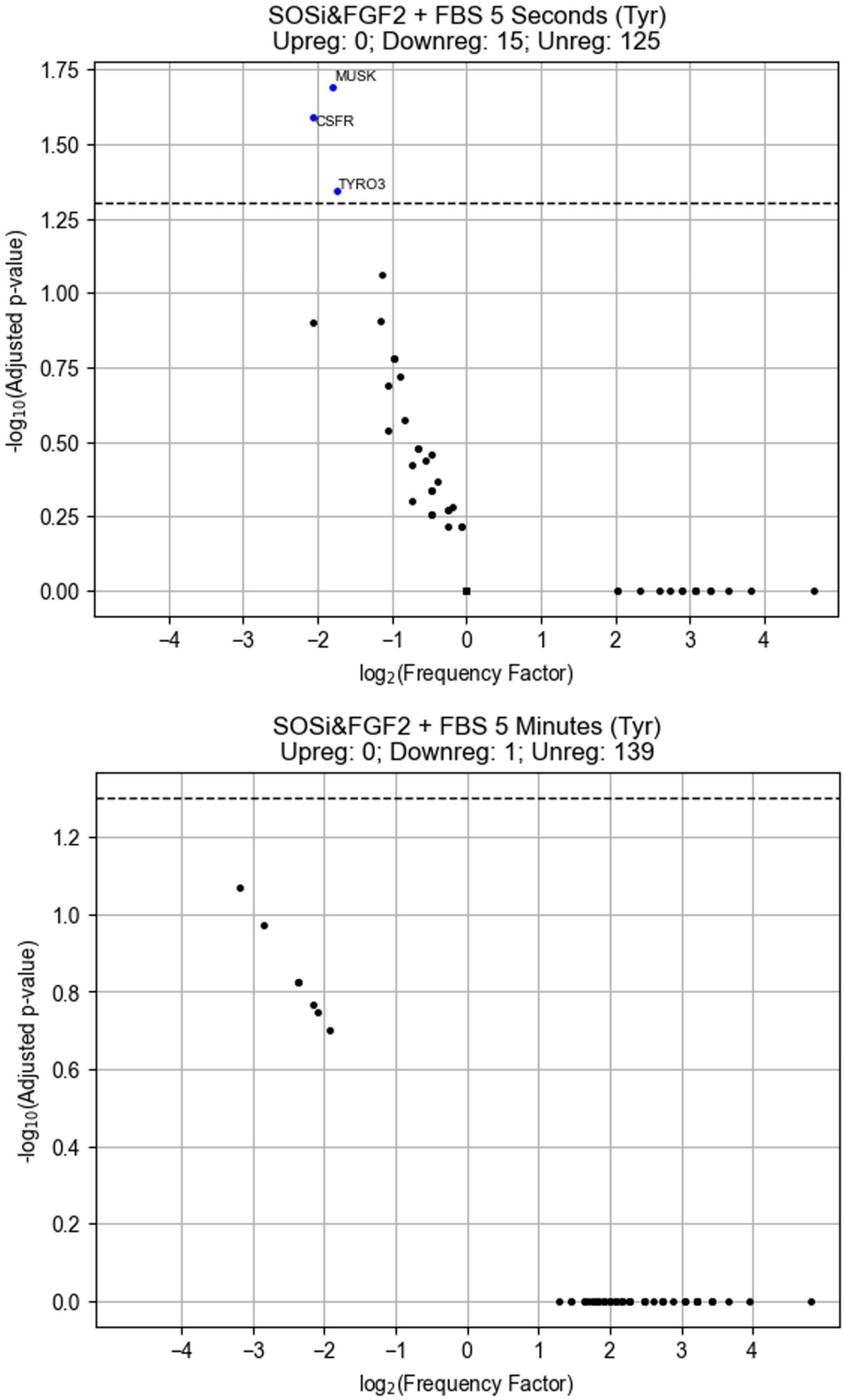


**Figure S33:** Tyrosine kinase activity (SOSi/ FGF2 plus FBS 5 Seconds and 5 minutes). Kinase library, FDR 0.05%, kin_type = 'tyr', method = 'percentile_rank', thresh = 8.

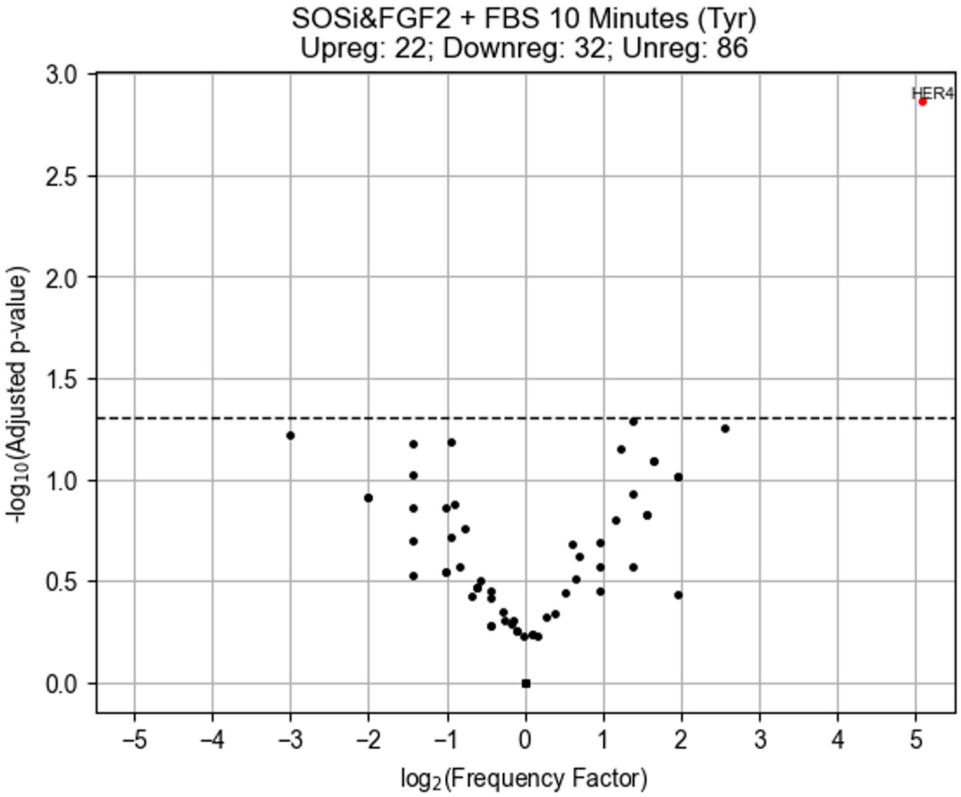


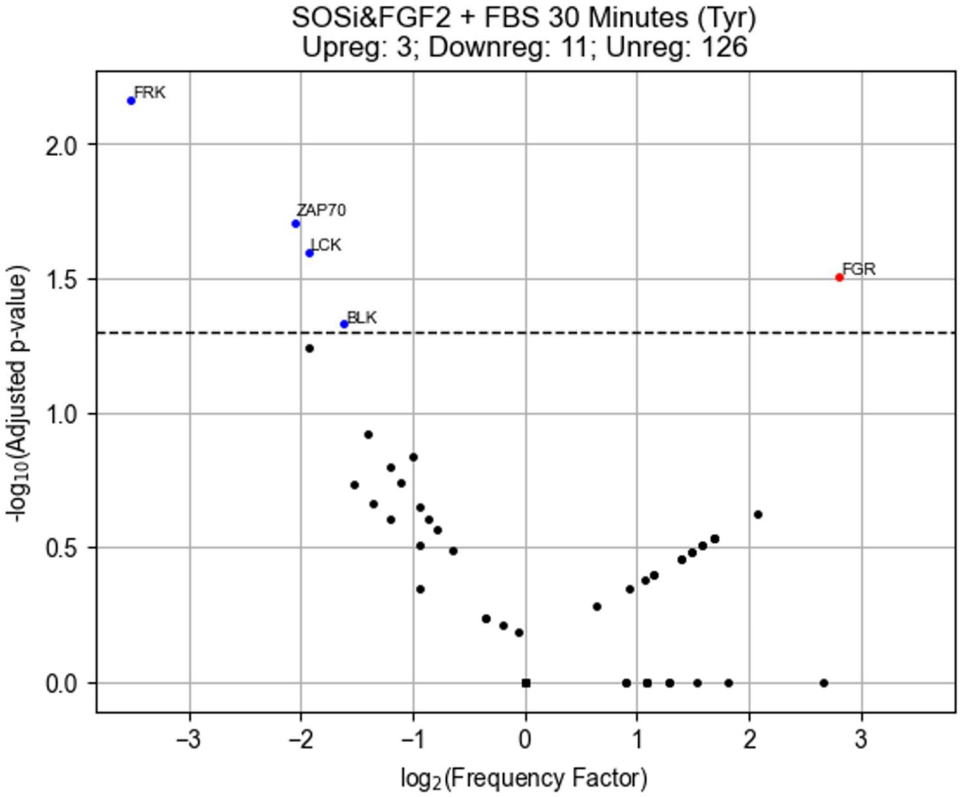


**Figure S34:** Tyrosine kinase activity (SOSi/ FGF2 plus FBS 10 minutes and 30 minutes). Kinase library, FDR 0.05%, kin_type = ’tyr’, method = ’percentile_rank’, thresh = 8.
